# Virtual screening for small molecule pathway regulators by image profile matching

**DOI:** 10.1101/2021.07.29.454377

**Authors:** Mohammad H. Rohban, Ashley M. Fuller, Ceryl Tan, Jonathan T. Goldstein, Deepsing Syangtan, Amos Gutnick, Ann DeVine, Madhura P. Nijsure, Megan Rigby, Joshua R. Sacher, Steven M. Corsello, Grace B. Peppler, Marta Bogaczynska, Andrew Boghossian, Gabrielle E. Ciotti, Allison T. Hands, Aroonroj Mekareeya, Minh Doan, Jennifer P. Gale, Rik Derynck, Thomas Turbyville, Joel D. Boerckel, Shantanu Singh, Laura L. Kiessling, Thomas L. Schwarz, Xaralabos Varelas, Florence F. Wagner, Ran Kafri, T.S. Karin Eisinger-Mathason, Anne E. Carpenter

## Abstract

Identifying chemical regulators of biological pathways is a time-consuming bottleneck in developing therapeutics and research compounds. Typically, thousands to millions of candidate small molecules are tested in target-based biochemical screens or phenotypic cell-based screens, both expensive experiments customized to each disease. Here, our uncustomized, virtual profile-based screening approach instead identifies compounds that match to pathways based on phenotypic information in public cell image data, created using the Cell Painting assay. Our straightforward correlation-based computational strategy retrospectively uncovered the expected, known small molecule regulators for 32% of positive-control gene queries. In prospective, discovery mode, we efficiently identified new compounds related to three query genes, and validated them in subsequent gene-relevant assays, including compounds that phenocopy or pheno-oppose YAP1 overexpression and kill a Yap1-dependent sarcoma cell line. This image profile-based approach could replace many customized labor- and resource-intensive screens and accelerate the discovery of biologically and therapeutically useful compounds.

**One sentence summary:** If a genetic perturbation impacts cell morphology, a computational query can reveal compounds whose morphology “matches”.

## Introduction

The pace of defining new diseases is rapidly accelerating ^1^, as is the cost and time required to develop novel therapeutics ^2^, creating huge unmet need. The dominant drug-discovery strategies in the pharmaceutical industry and academia are target-based (biochemical) and phenotypic (cell-based) screening. Both require significant setup time, are tailored to a specific target, pathway, or phenotype, and involve physically screening thousands to millions of candidate compounds at great expense ^3^. Computational approaches that allow virtual screening of small molecule modulators of pathways using the published literature or existing experimental data are beginning to emerge to accelerate drug discovery ^4,5^, but their predictive power is rarely evaluated systematically and prospectively.

Here we test a straightforward, image-based computational matching strategy; querying a compound library using image-based profiles of genes has not to our knowledge been previously attempted systematically. We use existing public data capturing the complex morphological responses of cells to a genetic perturbation (in the microscopy assay, Cell Painting ^6^), then identify small molecules (i.e., chemical compounds) that produce the same (or opposite) response. In this assay, more than a thousand quantitative morphology features (including size, shape, intensity, texture, correlation, and neighbor relationships) are extracted from five-color images of cells stained with six fluorescent stains that label eight cellular components or organelles (Figure 1b), to create an image-based profile of the sample. Conceptually similar to transcriptional profiling ^7^, image-based profiling assays like Cell Painting are cheaper and already proven in many applications ^8,9^.

**Figure 1:**
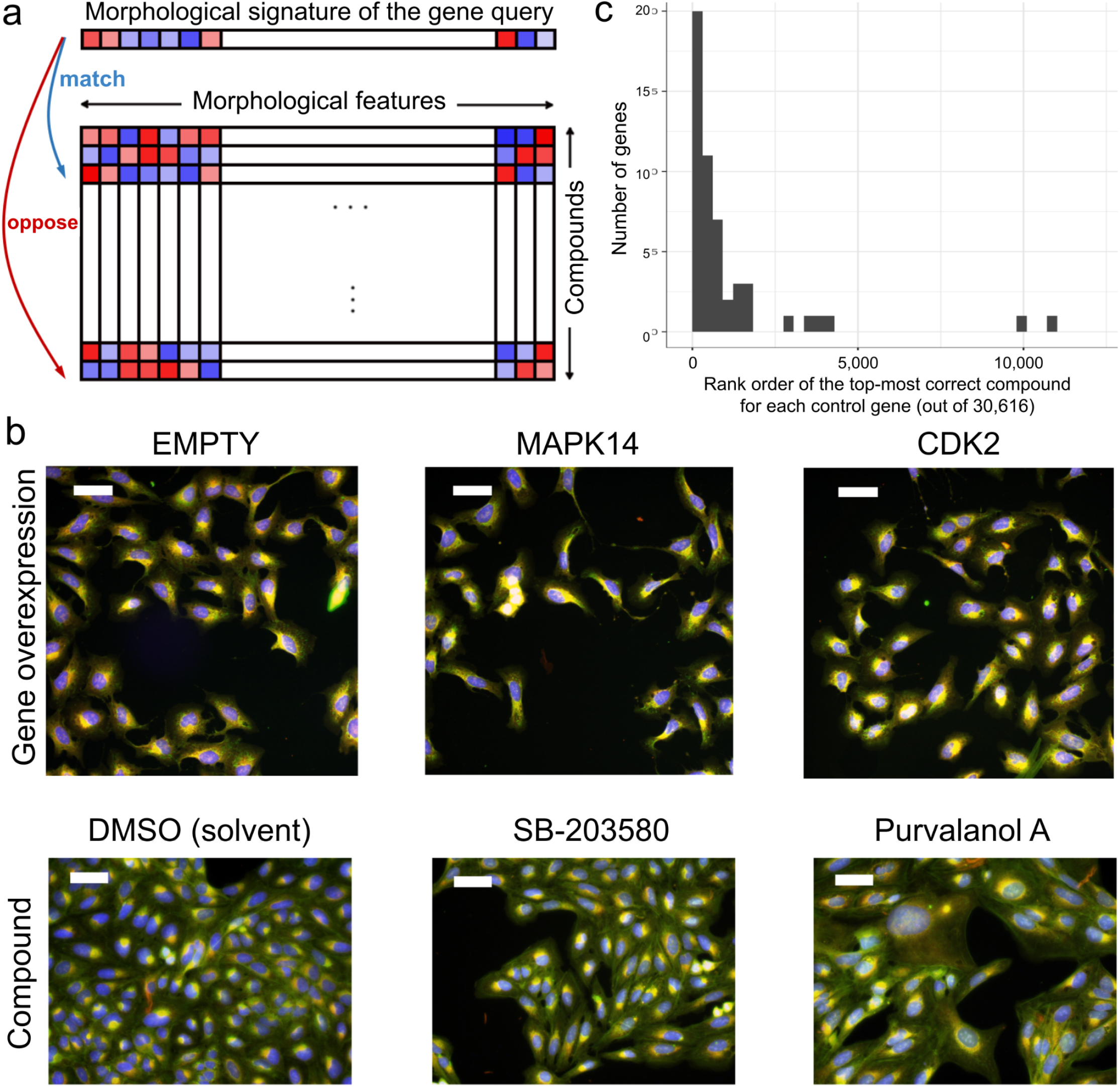
Image profile-based drug discovery offers efficient, virtual screening for pathway modulators. a) If an overexpressed gene changes the morphology of cells, its image-based profile can be used as a query in a database of small molecule profiles, looking for those that match (positively correlate) or oppose (negatively correlate). b) Cell Painting images for two positive control gene-compound matches that yield morphological phenotypes observable by eye (not all are expected to). EMPTY and DMSO are the negative controls in the gene overexpression and compound experiments, respectively; they differ in confluency and image acquisition conditions. The phenotype of p38a (MAPK14) overexpression matches (correlates to) that of SB-203580, a known p38 inhibitor; in both, elongated cells are over-represented. The phenotype of CDK2 overexpression (small cells) negatively correlates to that of purvalanol-a, a known CDK inhibitor, which induces an opposite phenotype (huge cells). Scale bars= 60 μm. c) The rank of the top correct compound for the control genes. Out of 63 genes with known compound matches in the set, 20 genes (32%) had a correct compound match in the top 1% of the 30,616 possible compounds (13 had a correct compound match in the top 0.5% (21%) and 36 genes (57%) had a correct compound match in the top 2.5%). Breakpoints in the histogram are set to [0, 306, 612, …] to correspond with 1% increments. The x-axis does not extend to all 30,616 compounds because all data points are in the top ~12,000.

Using image-based data to identify compounds that match a gene query has never been tested across a set of genes, making it difficult to estimate its potential. It is unknown whether gene perturbations yield profiles specific enough such that correlating compounds are likely to impact the function of a gene. It is also unknown whether anti-correlations of image-based profiles could reveal biological relationships. Evidence in favor of the strategy include anecdotal cases where perturbations of genes phenocopy the compounds targeting them ^10^, though these usually were done in the context of having a compound of interest and looking for the target instead of vice versa. Almost two decades of research indicates that image profiles can identify compounds inducing phenotypes that match other *compounds* ^11^ (reviewed in ^9^); this can be helpful in determining a compound’s mechanism of action (if the matching compounds are annotated)^12–16^ and in identifying novel chemical structures with behavior similar to known compounds with desired bioactivity. This latter case can be useful for finding compounds with better physical-chemical or physiological properties, or working around intellectual property concerns, but it requires having a compound with the desired biological impact already. Finding compounds that impact a gene/pathway of interest where existing compounds are *not* already known is a much harder task, and much more likely to yield groundbreaking medicines.

Recent decades have given rise to an appealing, reductive ideal in the pharmaceutical industry: one drug that targets one protein to target one disease ^17^. However, diseases often involve many interacting proteins and successful drugs often impact multiple targets ^18–20^. There is therefore a renewed appreciation for identifying small molecules that can modulate pathways or networks in living cell systems to yield a desired phenotypic effect ^17^. Because genes in a pathway often show similar morphology ^21^ and compounds often show similar morphology based on their mechanism of action as summarized above ^9^, we examined image profile matching as a promising but untested route to capturing perturbations at the pathway level and accelerating the screening step prior to identifying useful therapeutics and research tool compounds.

## Image-based gene-compound matching: validation

We began with 69 unique genes whose overexpression yields a distinctive morphological phenotype by Cell Painting, from our prior study in U2OS cells ^21^; roughly 50% of overexpression reagents in that study passed this criterion as did 45-54% in a more recent test of 160 overexpressed genes (Chandrasekaran et al.). We matched their image-based profiles to our public Cell Painting profiles of 30,616 small molecules ^22^ (Figure 1a), using simple Pearson correlation of population-averaged profiles (see “Scoring gene-compound and compound-compound connections” in Methods), avoiding machine learning methods due to their potential for overfitting and potential heightened sensitivity to experimental batch effects. We restricted matching to the 15,863 tested compounds (52%) whose profiles are distinguishable from negative controls, and confirmed that the profiles show variety rather than a single uniform toxic phenotype (Supplementary Figures S1 and S2).

We first verified that image-based profiles allow compounds to be matched with other compounds that share the same mechanism of action (for the subset that is annotated with this information). Consistent with past work ^9^, top-matching compound pairs share a common annotated mechanism-of-action four times more often than for the remainder of pairs (p-value < 5×10^−5^ permutation test on the Odds ratio, Supplementary Figure S3 and “Permutation test for validation of known compound-compound pairs” in Methods).

We next attempted gene-compound matching. We did not expect a given compound to produce a profile that matches that of its annotated gene target in all cases, nor even the majority. Expecting simple gene-compound matching takes a reductionist view that does not reflect the complexity of typical drug action (see Introduction). We therefore included genes annotated as physically interacting with the gene of interest (using BioGRID; see “Compound annotations” in Methods) as a correct match, given our goal of identifying compounds with the same functional impact in the cell and not only directly targeting the protein product of the gene of interest. In addition, existing annotations are imperfect, particularly given the prevalence of under-annotation, mis-annotation, off-target effects, and polypharmacology, where small molecules modulate protein functions beyond the intended primary target ^18,19^. Finally, technical reasons can also confound matching. The genetic and compound experiments were conducted years apart and by different laboratory personnel, yielding batch effects. They were performed in U2OS cells which may not be relevant for observing the annotated gene-compound interaction. In addition, the negative controls in a gene overexpression experiment (untreated cells), and a small molecule experiment (treated with the solvent control DMSO), do not produce identical profiles (left column, Figure 1b), and must therefore be normalized to align the negative controls in the feature space (see “Feature set alignment” in Methods). Despite these concerns, we persisted because even if the strategy worked in only a small fraction of cases, *dozens* of virtual screens and small validations of shortlisted predicted compounds could be done for less than the cost of a single traditional screening campaign - testing each compound in singlicate typically costs within an order of magnitude of $1 USD, and companies may screen millions of compounds. Even though downstream characterization, optimization, and target deconvolution of compounds will still be necessary, as for any drug discovery campaign, this virtual pre-screening could still translate to substantial resource savings.

Assessing the 63 genes out of 69 that had a compound annotated as targeting that gene in the set, we found that 32% of the genes successfully matched a compound, using an analysis of the highest-matching compounds per gene, described next. Because the method is intended as a virtual pre-screen that would feed a small proportion of predicted compounds to physical testing, our analysis mirrored this strategy by counting that 20 of the 63 genes had a ‘correct’ compound in the top 1% best-matching/opposing compounds in the experiment (Figure 1c; 1% = 306 out of 30,616 compounds in the library, with the 14,753 compounds not expressing a Cell Painting phenotype placed at the end of the rank order; p-value = 0.026 using 844 random permutations; see “Scoring gene-compound and compound-compound connections” and “Permutation test for validation of number of genes with a relevant compound match” in Methods); 1% represents a typical screen hit rate and a practical proportion for a confirmation screen of shortlisted compounds. Here, “correct” matches include cases where the corresponding protein of a gene interacts physically with at least one known annotated target of the compound (via BioGrid, 98.6% physical interactions). While 32% of genes were successful at the 1% shortlist level, the success rate increased to 36 genes (57%) having a correct compound match in the top 2.5% (Figure 1c).

We performed an alternate analysis, focusing not on one compound list per gene as in the two analyses above, but rather on the single ranked list of all 69 × 1177 gene-compound pairs together (69 genes with a phenotype and 1177 compounds that target any of them). Top-matching gene-compound pairs are correct matches 2.5 times more often than for the remainder of pairs (p-value = 0.002 one-sided Fisher test; Supplementary Figure S4 and Supplementary Table S1).

For some matches, we visually confirmed that gene overexpression phenocopies or pheno-opposes the matching/opposing compound (Figure 1b), although we emphasize that computationally-discovered phenotypes are not always visible to the human eye, particularly given cell heterogeneity. Examining the potential impact of polypharmacology, we found no relationship between the strength of correct gene-compound matches and the number of annotated targets a compound has (Supplementary Figure S5 and S6).

Throughout this study, we looked for compounds that both match (positively correlate) and oppose (negatively correlate) each overexpressed gene profile for several reasons. First, inhibitors and activators of a given pathway may both be of interest. Second, it is known that negative correlations among profiles can be biologically meaningful ^21^. Third, overexpression may not increase activity of a given gene product in the cell; it could be neutral or even decrease it via a dominant-negative or feedback loop effect. Fourth, the impact of a gene or compound perturbation could be cell-type specific. Supporting these theoretical arguments, we found that, empirically, both positively- and negatively-correlating matches were seen in our validation set (Supplementary Figure S4e). Furthermore, among the top 12 known gene-compound matches in our validation set, six showed correlation of the opposite directionality than expected (where expected is that an inhibitor’s profile would have the opposite correlation to its overexpressed target gene, Supplementary Figure S7).

Framing our approach as a virtual pre-screen, success even for 32% of Cell Painting-compatible genes would eliminate the need to carry out many dozens of large-scale customized screens in the pharmaceutical setting, instead advancing a few hundred compounds immediately to disease-relevant assays and saving hundreds of millions of dollars (see also Discussion for elaboration on success rates).

## Image-based gene-compound matching: discovery

We next searched virtually for novel small molecule regulators of pathways (defined loosely here as groups of genes whose disruption has a similar impact on biological outcomes). For each of the 69 genes, we created a rank-ordered list of compounds (from the 15,863 impactful compounds of the 30,616 set) based on the absolute value of correlation to that gene, enforcing a minimum of 0.35 (https://github.com/carpenterlab/2021_Rohban_submitted/blob/1de4fe928c30b8118a19be0736d15c3adae0d7d9/corr_mat.csv.zip). Because there is no systematic experiment to validate compounds impacting diverse pathways, we took a customized expert-guided approach to ensure the results are biologically meaningful rather than just statistically significant. We found seven experts studying pathways with strong hits who were willing to conduct experiments; they chose the most relevant biological systems and readouts, rather than simply attempting to validate the original image-based finding. The experts decided on the number of compounds reasonable for them to test in their experimental assay setup, and we provided them with the top matches from our list (subject to availability of sufficient compound). In a pharmaceutical context, we would recommend choosing the top 1% of compounds to test (n=306 in this case); due to throughput limitations here, we tested < the top 0.1% of the compounds (n = 9-33, except for RAS where n = 236).

Two cases yielded no confirmation (data not shown): SMAD3 and RAS. Nine compounds matching or opposing the SMAD3 overexpression profile failed to yield activity in a transcription reporter assay in A549 lung carcinoma cells involving tandem Smad binding elements, with and without Transforming growth factor beta 1 (TGF-β1). 236 compounds with positive or negative correlations to the wild-type RAS or oncogenic HRAS G12V differential profile (see Methods) failed to elicit a RAS-specific response in a 72-hour proliferation assay using isogenic mouse embryonic fibroblast (MEF) cell lines driven by human KRAS4b G12D, HRAS WT, or BRAF V600E alleles but otherwise devoid of RAS isoforms ^23^. We cannot distinguish whether the compounds were inactive due to major differences in the cell types or readouts, or whether these represent a failure of morphological profiling to accurately identify modulators of the pathway of interest.

A third case affirmed the approach but the novel compound identified was not very potent. We tested 17 compounds that negatively correlated with CSNK1E overexpression in a biochemical assay for the closely related kinase CSNK1A1. Three (SB 203580, SB 239063, and SKF-86002) had inhibitory IC_50_ concentrations in the nanomolar range at K_m_ ATP. Inhibition of CSNK1 family members by these compounds is supported by published kinase profiling studies ^24–26^. A fourth compound, BRD-K65952656, failed to bind any native kinases in a full KINOMEscan panel, suggesting it mimics CSNK1A1 inhibition via another molecular target. We chose not to pursue the expensive step of target deconvolution given its weak inhibition of CSNK1A1 (IC_50_ 12 μM).

A fourth case affirmed the approach but the novel compound failed to replicate following compound resynthesis, suggesting the desired activity, although validated, was not due to the expected structure, perhaps due to breakdown. We tested 16 compounds that positively correlated and 17 compounds that negatively correlated to GSK3B overexpression, for impact on GSK3α and GSK3β (which generally overlap in function) in a non-cell-based, biochemical assay. This yielded four hits with GSK3α IC50s ≤ 10 μM; the two most potent failed to show activity following resynthesis and hit expansion (testing of similarly-structured compounds) (Supplementary Table S2).

We did not pursue these cases further in light of the success for the three other cases, described next.

## Discovery of hits modulating the p38a (MAPK14) pathway

p38a (MAPK14) inhibitors are sought for a variety of disorders, including cancers, dementia, asthma, and COVID-19 ^27,28^. We chose 20 compounds whose Cell Painting profile matched (n=9) or opposed (n=11) that of p38α overexpression in U2OS cells. In a single-cell p38 activity reporter assay in retinal pigment epithelial (RPE1) cells ^29,30^, we identified many activating compounds; these are less interesting given that the p38 pathway is activated by many stressors but rarely inhibited. We also found several inhibiting compounds and confirmed their activity (Figure 2, Supplementary Figure S8), including a known p38 MAPK inhibitor; most had diverse chemical structures (Supplementary Figure S9a). Although the novel compounds are relatively weak, they nevertheless prove the principle that p38 pathway modulators can be found by image-profile matching, without a specific assay for the gene’s function.

**Figure 2:**
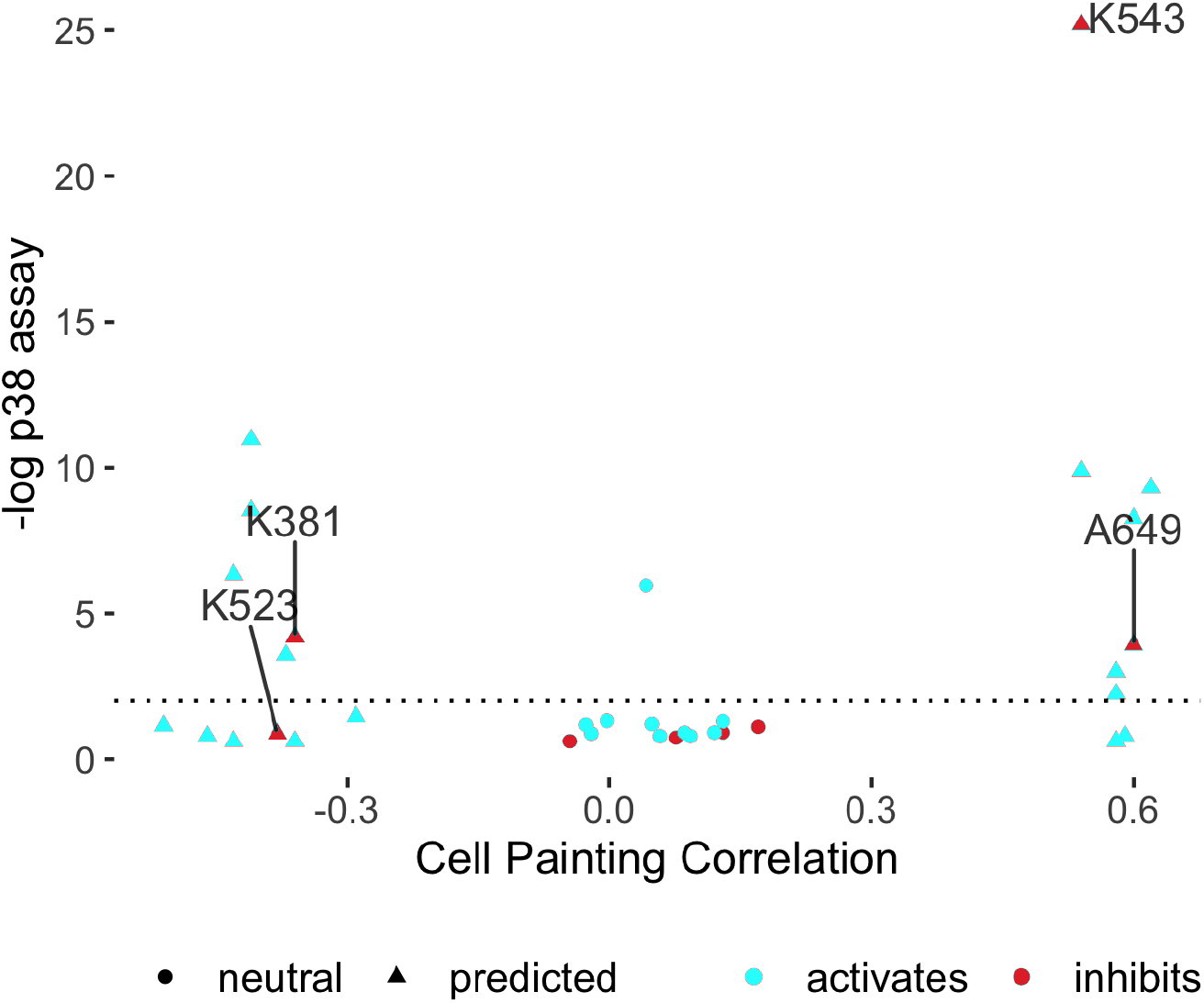
Cell Painting profiles identify compounds impacting the p38 pathway. Compounds predicted to perturb p38 activity (triangles) and a set of 14 neutral compounds (Cell Painting profile correlations to p38α between −0.2 to 0.2; circles) were tested for their influence on p38 activity at 1 μM using a two-sided t-test on the single cell distributions of a p38 activity reporter^31^ (FDR-adjusted -log_10_ p-values shown). Two potential inhibitors were found (BRD-K38197229 <K381> and BRD-A64933752 <A649>); an additional one (BRD-K52394958 <K523>) was identified via an alternative statistical test (Supplementary Figure S8a, h-i). K543 (BRD-K54330070) denotes SB-202190, a known p38 inhibitor found as a match (other known inhibitors such as SB-203580 from Figure 1 were strong matches but excluded from this experiment because they were already known).

## Discovery of hits impacting PPARGC1A (PGC-1α) overexpression phenotypes

We next identified compounds with strong morphological correlation to overexpression of peroxisome proliferator-activated receptor gamma coactivator 1-alpha (PGC1α, encoded by the PPARGC1A gene). We found that these compounds tend to be hits in a published, targeted screen for PGC1α activity (p=7.7e-06, Fisher’s exact test) ^32^, validating our image profile-based matching approach. The dominant matching phenotype is mitochondrial blobbiness, which can be quantified as the high standard deviation of the MitoTracker staining at the edge of the cell (Figure 3a,b) without major changes to cell proliferation, size, or overall protein content. Cell subpopulations that are large, multi-nucleate, and contain fragmented mitochondria are over-represented when PGC-1α is overexpressed while subpopulations whose organelles are asymmetric are under-represented (Supplementary Figure S10). More symmetric organelle morphology has been associated with reduced motility and PGC-1α overexpression ^33^. The role of PGC-1α in mitochondrial biogenesis is well-appreciated ^34^. The phenotype uncovered here using image profile matching is consistent with other recently discovered mitochondrial phenotypes associated with this gene ^35^.

**Figure 3:**
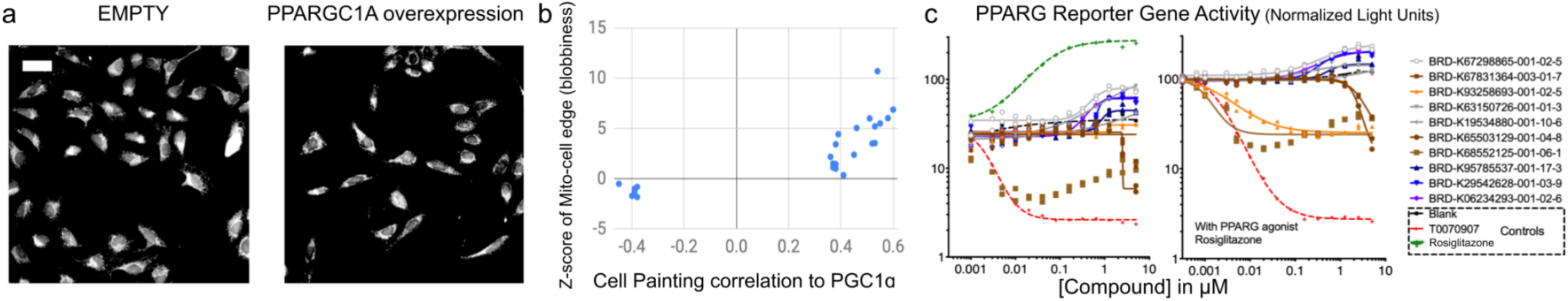
Cell Painting profiles identify compounds impacting PPARGC1A (PGC-1α) overexpression phenotypes. a) Cell Painting images for PPARGC1A (PGC-1α) overexpression compared to negative control (EMPTY, same field as in Figure 1a). Scale bar = 60 μm. b) Compounds with high or low correlations of their Cell Painting profiles to PGC-1α overexpression were chosen for further study (hence all samples are below ~ −0.35 or above ~0.35 on the X axis). Correlation to PGC-1α overexpression is dominated by one feature, the standard deviation of the MitoTracker staining intensity at the edge of the cell, which we term blobbiness (displayed on the Y axis as a z-score with respect to the negative controls). c) PPARG reporter gene assay dose-response curves in the absence (left) or presence (right) of added PPARG agonist, Rosiglitazone. Representative data of the ten most active compounds is shown and reported as normalized light units. Compounds highlighted in blue/purple are structurally related pyrazolo-pyrimidines.

We chose 24 compounds whose Cell Painting profiles correlated positively or negatively with PGC-1α overexpression in U2OS cells (Supplementary Table S3); they have generally diverse chemical structures (Supplementary Figure S9b) and one is a known direct ligand for PPAR gamma, GW-9662 (BRD-K93258693). PGC-1α is a transcriptional coactivator of several nuclear receptors including PPAR gamma and ERR alpha ^36^. We therefore tested compounds in a reporter assay representing FABP4, a prototypical target gene of the nuclear receptor, PPARG ^37^, in a bladder cancer cell line (Figure 3c). Three of the five most active compounds leading to reporter activation (among all the 24 compounds tested) were structurally related and included two annotated SRC inhibitors, PP1 and PP2, which have a known link to PGC1a ^38^, as well as a novel analog thereof. Inhibitors uncovered were CCT018159 (BRD-K65503129) and Phorbol 12-myristate 13-acetate (BRD-K68552125). Many of the same compounds also showed activity in a ERRalpha reporter assay in 293T cells, albeit with differing effects (Supplementary Figure S11).

Encouraged by these results, we tested the impact of the compounds on mitochondrial motility, given the mitochondrial phenotype we observed and the role of PGC1a in mitochondrial phenotypes and neurodegenerative disorders ^39^. In an automated imaging assay of rat cortical neurons ^40^, we found several compounds decreased mitochondrial motility; none increased motility (Supplementary Figure S12). Although the latter is preferred due to therapeutic potential, this result suggests that the virtual screening strategy, applied to a larger set of compounds, might identify novel motility-promoting compounds. We found 3 of the tested compounds suppress motility but do not decrease mitochondrial membrane potential; this is a much higher hit rate (13.0%) than in our prior screen of 3,280 bioactive compounds, which yielded two such compounds (0.06%)^40^.

## Discovery of small molecules impacting YAP1-related phenotypes

The Hippo pathway affects development, organ size regulation, and tissue regeneration. Small molecule regulators are highly sought for research and as potential therapeutics for cancer and other diseases; the pathway has been deemed relatively undruggable ^41,42^. We tested 30 compounds (Supplementary Table S4) whose Cell Painting profile matched (25 compounds) or opposed (5 compounds) the overexpression of the Hippo pathway effector Yes-associated protein 1 (YAP1), which we previously explored ^21^ (Supplementary Table S5, images shown in Supplementary Figure S13). The compounds have generally diverse chemical structures (Supplementary Figure S9c). One hit, fipronil, has a known tie to the Hippo pathway: its impact on mRNA profiles matches that of another calcium channel blocker, ivermectin, a potential YAP1 inhibitor ^43^ (99.9 connectivity score in the Connectivity Map^7^). After identifying five promising compounds from the 30 compounds tested in a cell proliferation assay in KP230 cells (described later), we focused on the three strongest in various assays and cell contexts, as follows.

N-Benzylquinazolin-4-amine (NB4A, BRD-K43796186) is annotated as an EGFR inhibitor and shares structural similarity with kinase inhibitors. NB4A showed activity in 30 of 606 assays recorded in PubChem, one of which detected inhibitors of TEAD-YAP interaction in HEK-TIYL cells. Its morphological profile positively correlated with that of YAP1 overexpression (0.46) and, consistently, negatively correlated with overexpression of STK3/MST2 (−0.49), a known negative regulator of YAP1.

Because the Hippo pathway can regulate the pluripotency and differentiation of human pluripotent stem cells (hPSCs) ^44,45^, we investigated the effect of NB4A in H9 hPSCs. NB4A did not affect *YAP1* mRNA expression but increased the expression of YAP1 target genes (*CTGF* and *CYR61*) in a dose-dependent manner (Figure 4a), confirming it impacts YAP1 phenotypes. Accordingly, NB4A increased YAP1 nuclear localization (Figure 4b). While decreasing total YAP1 protein levels, NB4A also reduced YAP1 S127 phosphorylation (Figure 4c and Supplementary Figure S14a), which promotes YAP1 cytoplasmic sequestration ^46^.

**Figure 4:**
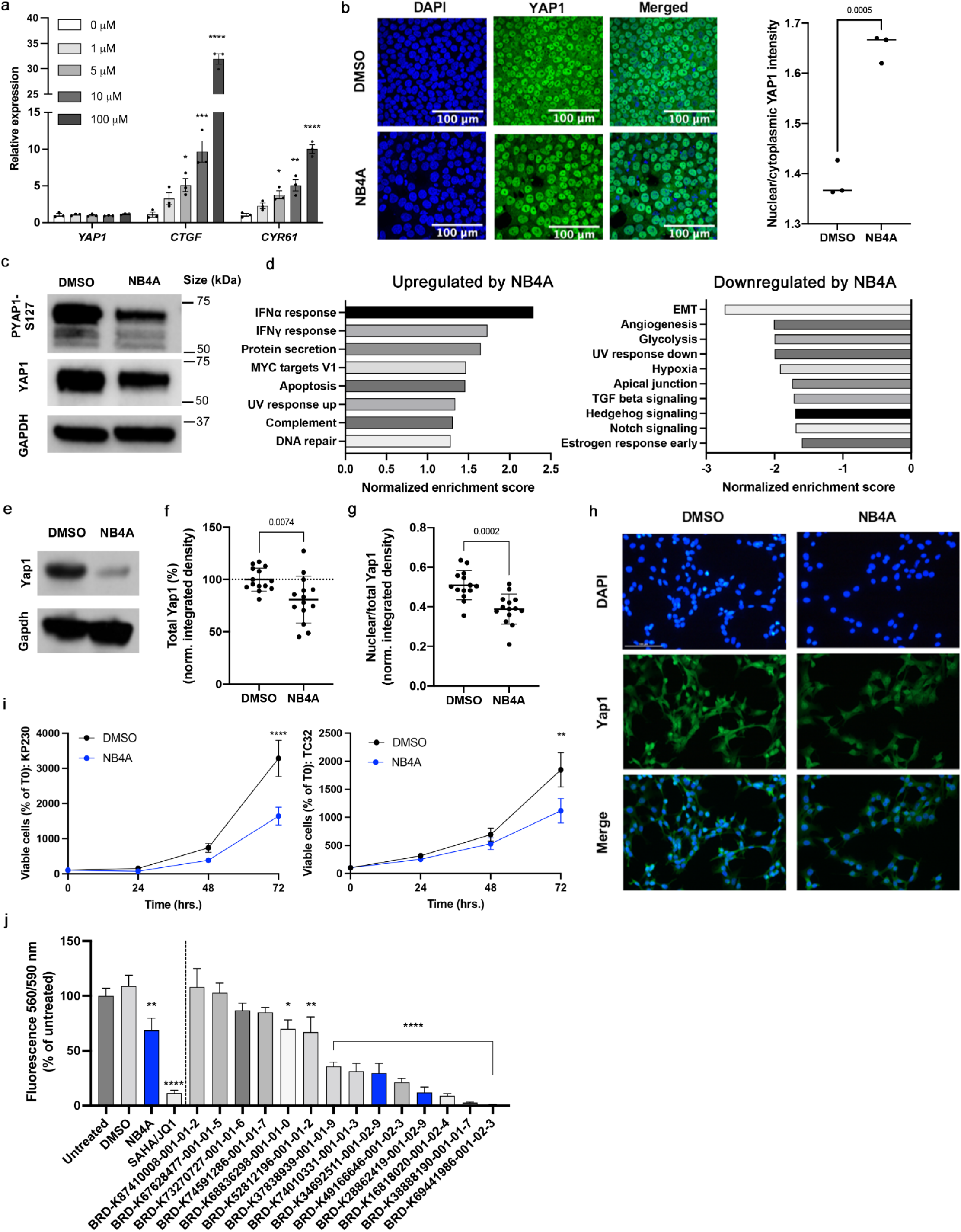
Cell Painting profiles identify compounds impacting YAP1 phenotypes. a) Relative transcript levels of YAP1, CTGF, and CYR61 in H9 human pluripotent stem cells treated with NB4A or DMSO control for 24 hrs. *P<0.05; **P<0.01; ***P<0.001; ****P<0.0001 (one-way ANOVA with Dunnett’s multiple comparisons test). Mean ± SEM. n = 3 biologically independent experiments. b) Representative images of YAP1 immunofluorescence (left) and quantification of nuclear/cytoplasmic YAP1 mean intensity (right) in H9 cells after treatment with 10 μM NB4A or DMSO control for 24 hours. Two-tailed student’s t-test. n = 3 biologically independent experiments; an average of mean intensities from 3 fields of each biological replicate is calculated. c) Representative blot of n = 3 biologically independent experiments for phospho-YAP1 (S127) and total YAP1 from H9 cells treated with DMSO or 10 μM NB4A for 24 hrs, with GAPDH as loading control (quantified in Supplementary Figure S14a). d) Normalized enrichment scores of GSEA show up to 10 of the most significant Hallmark pathways up- and down-regulated in NB4A-treated vs. control KP230 cells (FDR-adjusted P<0.25). n = 3. IFNα = interferon alpha; IFNγ = interferon gamma; EMT = epithelial-mesenchymal transition. e) Representative western blot for Yap1 in NB4A-treated and control KP230 cells. f) Immunofluorescence-based analysis of total Yap1 in NB4A-treated and control KP230 cells. Two-tailed student’s t-test. Mean ± SEM. n = 3. g) Immunofluorescence-based analysis of nuclear Yap1 in NB4A-treated and control KP230 cells (normalized to total Yap1). Two-tailed student’s t-test. Mean ± SEM. n = 3. For f and g, the Y axis is integrated density normalized to cell number and representative images are shown in (h), out of 5 fields acquired per condition. Scale bar (top left panel) = 100 μM. i) Growth curves of NB4A-treated and control KP230 and TC32 sarcoma cells. **P<0.01; ****P<0.0001 DMSO vs. NB4A (72 hrs.; 2-way ANOVA with Sidak’s multiple comparisons test). Mean ± SEM. n = 3. For panels d-i, cells were treated with 10 μM NB4A daily for 72 hours. j) PrestoBlue screen of KP230 cells treated with various analogs of BRD-K28862419 and BRD-K34692511 (10 μM daily for 48 hours). *P<0.05; **P<0.01; ****P<0.0001 vs. DMSO (one-way ANOVA with Dunnett’s multiple comparisons test). Mean ± SEM. n = 2 biological replicates, each with 3 technical replicates. SAHA (2 μM)/JQ1 (0.5 μM), which inhibits Yap1 expression and transcriptional activity in sarcoma cells ^47^, was used as a positive control. Blue bars indicate the original hits (NB4A, BRD-K28862419 and BRD-K34692511). All test compounds were verified at purity > 75%.

Effects of NB4A on YAP1 mRNA expression were not universal across cell types, consistent with the Hippo pathway’s known context-specific functions. In most cell types represented in the Connectivity Map, *YAP1* mRNA is unaffected, but in HT29 cells, *YAP1* mRNA is up-regulated after six hours of NB4A treatment (z-score = 3.16; also z-score = 2.04 for TAZ) and in A375 cells, *YAP1* mRNA is slightly down-regulated (at 6 and 24 hours; z-score ~ −0.7) ^7^. NB4A had no effect in a YAP1-responsive reporter assay following 48h of YAP overexpression in HEK-293 cells (Supplementary Figure S14b).

Compounds influencing the Hippo pathway might be therapeutic for undifferentiated pleomorphic sarcoma (UPS), an aggressive mesenchymal tumor that lacks targeted treatments ^47^. In UPS, YAP1 promotes tumorigenesis and is inversely correlated with patient survival ^47^. In KP230 cells, derived from a mouse model of UPS 47, Yap1 protein levels were reduced after 72 hours of NB4A treatment (Figure 4e-f, h). NB4A also significantly attenuated Yap1 nuclear localization (Figure 4g-h), which is known to reduce its ability to impact transcription. Interestingly, NB4A did not directly alter transcription of *Yap1*, its sarcoma target genes (*Foxm1, Ccl2, Hbegf, Birc5*, and *Rela*), nor Yap1’s negative regulator, angiomotin (*Amot*) (data not shown). Instead, pathways such as interferon alpha and gamma responses were up-regulated, whereas pathways such as the epithelial-mesenchymal transition, angiogenesis, and glycolysis were down-regulated, according to RNA sequencing and gene set enrichment analysis (Figure 4d; Supplementary Table S6). This indicates a potentially useful mechanism distinct from transcriptional regulation of *YAP1*.

Genetic and pharmacologic inhibition of Yap1 is known to suppress UPS cell proliferation *in vitro* and tumor initiation and progression *in vivo* ^47^. Consistent with being a Hippo pathway regulator, NB4A inhibited the proliferation of two YAP1-dependent cell lines: KP230 cells and TC32 human Ewing’s family sarcoma cells ^48^ (Figure 4i). NB4A did not affect the proliferation of two other YAP1-dependent lines, STS-109 human UPS cells (Supplementary Figure S15a) and HT-1080 fibrosarcoma cells (Supplementary Figure S15b) ^47,49^, nor YAP1-independent HCT-116 colon cancer cells (Supplementary Figure S15c-e). Interestingly, NB4A treatment did not exhibit overt toxicity by trypan blue staining in any of these (not shown), suggesting it inhibits cell proliferation by a mechanism other than eliciting cell death.

Next, we investigated two structurally similar compounds (BRD-K28862419 and BRD-K34692511, distinct from NB4A’s structure) whose Cell Painting profiles negatively correlated with YAP1’s overexpression profile (−0.43 for BRD-K28862419 and −0.45 for BRD-K34692511) and positively correlated with TRAF2 overexpression (0.41 for BRD-K28862419 and 0.29 for BRD-K34692511) (Supplementary Figure S13). These compounds are not commercially available, limiting our experiments and past literature.

We assessed the compounds’ impact on mesenchymal lineage periosteal cells isolated from 4-day old femoral fracture callus from mice with DOX-inducible YAP-S127A. BRD-K34692511 substantially upregulated mRNA levels of relevant Hippo components including *Yap1* and *Cyr61* after 48 hours of treatment, but not at 1 and 4 hours (Supplementary Figure S14c-f). By contrast, the compounds had no effect on *YAP1* or its target genes in H9 hPSCs (Supplementary Figure S14g), nor in a 48 h YAP-responsive reporter assay following YAP overexpression in HEK-293 cells (Supplementary Figure S14b).

Like NB4A, the effects of these compounds on proliferation varied across cell types. In the U2OS Cell Painting images, BRD-K28862419 reduced proliferation (−2.0 st dev). Per PubChem, it inhibits cell proliferation in HEK293, HepG2, A549 cells (AC50 5-18 μM) and it inhibits PAX8, which is known to influence TEAD/YAP signaling^50^. BRD-K34692511 had none of these impacts.

Both compounds had the desired effect of inhibiting KP230 cell proliferation (Supplementary Figure S15f). Also noteworthy, BRD-K28862419 modestly yet significantly reduced KP230 cell viability (Supplementary Figure S15g), indicating its mechanism of action and/or therapeutic index may differ from that of NB4A and BRD-K34692511.

Finally, we resynthesized both of these non-commercially available compounds to ensure their integrity and identity, and selected a set of close analogs of such compounds to investigate structure-activity relationship. Using a high-throughput resazurin-based screen (Presto Blue assay), we confirmed the activity of the two resynthesized compounds to reduce proliferation of KP230 cells, and furthermore found eight additional analogs with activity (Figure 4J). In fact, four of the analogs were more potent than one or both of the original compounds, reducing KP230 cell growth by 78.70-98.95% vs. untreated cells (Figure 4J). Structural analysis reveals that the macrocycle is quite robust to changes in stereochemistry, and that compounds with an amide linker tend to outperform those with a urea linker (Supplementary Figure S16).

In summary, although deconvoluting the targets and behaviors of these compounds in various cell contexts remains to be further ascertained, we conclude that the strategy identified compounds that modulate YAP1-related phenotypes, in particular an unusual ability to reduce growth of certain aggressive sarcoma lines. This demonstrates that, although the directionality and cell specificity will typically require further study, image-based pathway profiling can identify modulators of a given pathway.

## Discussion

We found that hit-stage small molecule regulators of pathways of interest can be discovered by virtual matching of genes and compounds using Cell Painting profiles, which we term image profile-based compound screening. The approach provides a simple, minimal-resource approach to shortlist top candidates to target the pathway of a gene of interest, so long as the gene produces a detectable image-based profile. It can reduce the cost of compound screening by orders of magnitude by enriching the signal to noise ratio from compound libraries. We do not claim the particular compounds we uncovered in this study are sufficiently potent, specific, and non-toxic for human therapeutics. As with all screening approaches, significant further work is necessary to develop hits into useful therapeutics; this includes confirming activity and directionality of hits in a relevant cell type or model system (as we did successfully for three cases here), improving potency and specificity, and identifying the molecular target(s) (including so-called off-target effects that may or may not be useful to achieve the desired phenotypic effect); these latter are significant steps. Further, like all drug discovery, the eventual clinical success relies on the therapeutic hypothesis for the gene, pathway, and/or phenotype being correct, which is also challenging.

Even so, virtualizing a large-scale screen by computationally matching the phenotypic effect of compounds to that of gene manipulation will in many cases enable rapid and inexpensive identification of compounds with desired phenotypic impacts, and avoid an expensive large-scale physical screen. Our approach yielded hits with substantial chemical structure diversity, which is another advantage. Gene-compound matching might also be useful to identify which genes/pathways are targeted by novel small molecules of unknown mechanism of action, another significant bottleneck in the drug discovery process ^51^.

Other virtual drug screening methods have been proposed; to our knowledge, none have been systematically evaluated that take a gene name as input. For those using alternative strategies (e.g. predicting outcomes in a particular assay that is used to train the model), their success rates across a broad swath of biology are unavailable or difficult to compare to this study. Many report retrospective rates on known compound hits (which can unfortunately be prone to overfitting) but do not prospectively identify novel hits; others report success in identifying a novel compound prospectively but do not mention the failures, making it impossible to compute the success rate across multiple diverse genes. To our knowledge, ours is the only study to test a sufficient number of genes to report success rates at the gene level, both retrospectively (63 genes) and prospectively (7 genes). Other methods report predictions with ~8-70% success rates, but at the assay level ^52–54^ (and with varying degrees of stringency in how success is defined). In other words, they predict compounds’ assay activity rather than compounds impacting a given gene function. To study a new area of biology, these methods require (a) developing an assay to target the biological phenotype of interest (this criterion means these methods’ reported success rates have come from testing on only a subset of biological “space”, which is likely better-studied), and (b) testing a pilot set of compounds in the assay to train a machine learning model (typically thousands of compounds, which is resource intensive). This is much more intensive effort than what is needed for the strategy presented here: a morphological profile of a single gene perturbation (which will soon be publicly available at the genome-scale, see below, but can be done in any laboratory with a suitable microscope within a few days), and profiles for compounds (which are already publicly available in the tens of thousands). No other data is required (e.g. to pre-train models), nor special equipment, nor extensive computational expertise nor compute time.

As mentioned, most virtual compound prediction studies report only retrospective assay-level success rates - these are certainly valuable, but a lower bar that, unfortunately, can be inflated by unexpected sources of overfitting. For example, if too-similar assays or too-similar compounds are included in both training and test sets, success rates will be inflated; only some studies control for this well. To clarify success rates for this study in particular, using any gene as an input, roughly 50% will pass at the first step (having a distinguishable morphological phenotype under the assay conditions, though other cell types or stains could be attempted), and the remainder are estimated to succeed at a rate of 32-71% as follows: our study showed a 32% retrospective success rate for known gene-compound connections (for a 1% shortlist); given these known pairs include highly optimized chemical matter, we wondered whether we would see a lower success rate in actual prospective validation. Nevertheless, we discovered a 43% success rate (albeit with low n: 3 out of 7 genes tested); we also note the rate might even be considered 71% (5 of 7) if we consider the other two genes as successes whose hits were abandoned due to chemistry and potency. Due to resource limits, we tested ~0.1% of the best-matching compounds rather than the recommended 1%, which would likely also improve results (we tested for SMAD3: 9 compounds, RAS: 236, CSNK1A1: 17, GSK3B: 33, p38 MAPK14: 20, PPARGC1A: 24, YAP1: 30). Overfitting is not a concern in our study; we used simple metrics of correlation rather than machine learning that runs that risk; avoiding machine learning also may make matching of profiles across datasets created under very different conditions more successful. As a side note, we do not emphasize success rates in terms of how many of the selected compounds for each gene were validated in the followup assays, because in practice, hundreds of compounds should be shortlisted for testing and the major metric of interest to the screener is whether good chemical matter is in that set, not whether the rate within the set is 1% or 100%.

We expect future iterations of this strategy to be more successful. First, we would expect better-quality chemical matter from larger libraries; only 30,000 were screened in this work whereas a pharmaceutical screening campaign can test millions ^55^. Large-scale data production efforts are underway that will increase the potential for matching profiles against public data: the JUMP-Cell Painting Consortium is producing a public dataset of 140,000 chemical and genetic perturbations. It is remarkable that the image-based profile matching strategy worked for two datasets created years apart by different personnel and equipment; we expect improvements if the query gene and compound library were created more consistently. The limits of the approach remain to be tested: how different could image sets be, in terms of resolution, confluency, imaging modality, and even cell type? Creating data using other staining sets or more complex biological models, such as co-cultures, primary cells, or organoids could increase the probability of success for some pathways, as could assessing whether gene knockdown profiles (e.g. by CRISPR) yield better results in practice than gene overexpression. Pathways where overexpression and knockdown give opposite profiles may be even better starting points for virtual screening, as might merged profiles based on several pathway members of interest rather than a single gene.

Another major direction for the future is making better predictions by integrating morphology profiles with other data sources when available at scale, such as transcriptomic^56^, proteomic, and metabolomic^57^ profiles, or historical assay data^52^. Chemical structure information might also be useful^53^, though this would require significant adaptation to incorporate because it is not a property one can obtain for the genes used as queries in our matching approach, and the goal is not to identify compounds of similar structure (diversity is usually preferred). More advanced computational methods are also on the horizon, from feature extraction ^58^ to machine learning on new benchmark datasets of gene-compound pairs ^59^; we would expect supervised machine learning to work better than our unsupervised correlation-based approach ^9^. We hope that image profile-based virtual screening will be a new accelerant toward meeting the pressing need for novel therapeutics.

## Acknowledgements

The authors thank the researchers who originally helped produce the published data used in this analysis, including the Broad Institute LINCS team, Cancer Program and PRISM team. We appreciate helpful discussions with our colleagues, including Pere Puigserver, Evan Rosen, and Amit Majithia.

## Data and materials availability

All data, code, and materials used in the analysis are available to any researcher for purposes of reproducing or extending the analysis.

