## Supplementary material for "Virtual screening for small molecule pathway regulators by image profile matching": Supp Tables

***Supplementary tables***

**Supplementary Table 1: Gene-compound pairs that have morphological profiles with absolute correlation of at least 0.35 are around 2.5 more enriched in being relevant and trustworthy compared to the remainder of the pairs.** Odds ratio is 2.53 (=(16/6546)/(71/73545)) and p-value = 0.002 in the one sided Fisher's test. Note that this table considers all gene-compound pairs together in the ranked list (rankings are not done on a per-gene basis).

| | **Gene is among the compound targets** | **Gene is not among the compound targets** |
|---|---|---|
| $\|corr.\| \geq 0.35$ | 16 | 6,546 |
| $\|corr.\| < 0.35$ | 71 | 73,545 |

### Supplementary Table 2: GSK3 assay data and compound structures, as described in Methods

| Compound ID | Compound Name | SMILES | Structure | GSK3a IC$_{50}$, µM | GSK3b IC$_{50}$, µM | GSK3a selectivity ratio | Note |
|---|---|---|---|---|---|---|---|
| BRD-K00760705-001-05-1 | BRD0705 | CC[C@@]1(c2c[nH]nc2NC2=C1C(=O)CC(C)(C)C2)c1ccccc1 | 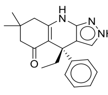 | 0.131 | 1.05 | 8.04 | Control |
| BRD-K21263731-001-06-7 | BRD3731 | CC(C)(C)Cc1[nH]nc2NC3=C(C(=O)CC(C)(C)C3)[C@](C)(c12)c1ccccc1 | 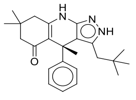 | 0.145 | 0.0109 | 0.07 | Control |
| BRD-K87550320-001-04-0 | BRD0320 | CC1(C)CC(=O)C2=C(C1)Nc1n[nH]c(C3CC3)c1[C@@]2(C)c1cccc(F)c1 | 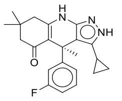 | 0.00828 | 0.00338 | 0.41 | Control |
| BRD-K16189898-001-12-8 | Chiron 99021 | Cc1c[nH]c(n1)-c1cnc(NCCNc2ccc(cn2)C#N)nc1-c1ccc(Cl)cc1Cl | 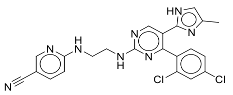 | 0.00294 | 0.00176 | 0.60 | Control |
| BRD-K55000304-001-11-9 | GW8510 | O=C1Nc2ccc3ncsc3c2\C1=C\Nc1ccc(cc1)S(=O)(=O)Nc1ccccn1 | 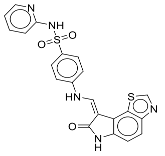 | 0.00710 | 0.00432 | 0.61 | Control |
| BRD-K59184148-001-23-2 | SB 216763 | Cn1cc(C2=C(C(=O)NC2=O)c2ccc(Cl)cc2Cl)c2ccccc12 | 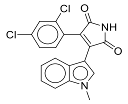 | 0.00424 | 0.00554 | 1.31 | Control |
| BRD-K26994486-001-01-0 | | C[C@@H](CO)N1C[C@H](C)[C@@H](CN(C)C(=O)Nc2c(C)noc2C)OCCCC[C@H](C)Oc2ccc(cc2C1=O)N(C)C | 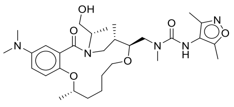 | 2.20 | 35.0 | 15.89 | active GSK3a |
| BRD-K91354313-001-01-6 | | CCC(=O)Nc1ccc2OC[C@H](C)N(C[C@H](C)[C@@H](CN(C)C(=O)c2c1)OC)C(C)=O | 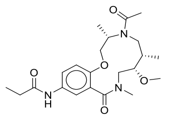 | 2.48 | 35.0 | 14.12 | active GSK3a |
| BRD-K02827191-001-02-4 | | C[C@@H](CO)N1C[C@@H](C)[C@H](CN(C)C(C)=O)Oc2ncc(cc2C1=O)C#Cc1ccc(F)cc1 | 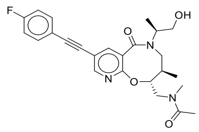 | 3.80 | 35.0 | 9.21 | active GSK3a |
| BRD-K52043064-001-01-3 | | CO[C@@H]1CN(C)C(=O)c2ccc(NC(=O)CN(C)C)cc2OC[C@@H](C)N(C[C@H]1C)C(=O)C1CCCCC1 | 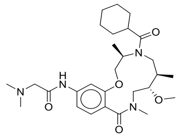 | 3.80 | 35.0 | 9.21 | active GSK3a |
| BRD-A62505706-001-03-1 | Edoxudine | CCc1cn(C2OCC(O)C2CO)c(=O)nc1O | 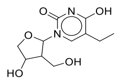 | 11.0 | 35.0 | 3.18 | borderline active GSK3a |
| BRD-K50011338-001-01-2 | | CO[C@H]1CN(C)C(=O)c2ccc(NC(=O)Nc3ccccc3)cc2OC[C@H](C)N(C[C@H]1C)C(C)=O | 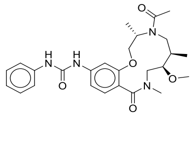 | 11.0 | 35.0 | 3.18 | borderline active GSK3a |

| | | | | | | |
|---|---|---|---|---|---|---|
| BRD-K87995704-001-01-9 | | CO[C@@H]1CN(C)C(=O)c2cc(NC(=O)NC3CCCCC3)ccc2OC[C@@H](C)N(CC2CC2)C[C@@H]1C | 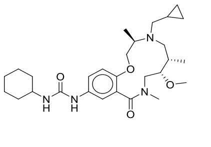 | 25.5 | 35.0 | 1.37 |
| BRD-K79711234-001-11-8 | Thiamphenicol | CS(=O)(=O)c1ccc(cc1)[C@@H](O)[C@@H](CO)NC(=O)C(Cl)Cl | 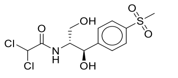 | 27.0 | 30.6 | 1.13 |
| BRD-K87264606-001-01-9 | | C[C@H](CO)N1C[C@H](C)[C@@H](CN(C)S(=O)(=O)c2cn(C)cn2)OCCCC[C@@H](C)Oc2ccc(cc2C1=O)N(C)C | 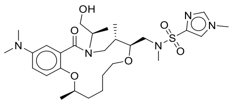 | 30.0 | 30.0 | 1.00 |
| BRD-K04407772-001-01-2 | | OC[C@@H]1O[C@H](CC(O)=O)C[C@@H]2[C@H]1Oc1ccc(NC(=O)CC3CC3)cc21 | 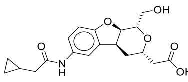 | 35.0 | 35.0 | 1.00 |
| BRD-K30491662-001-01-7 | | CO[C@H]1CN(C)C(=O)c2ccc(NC(=O)Nc3cccc(OC)c3)cc2OC[C@@H](C)NC[C@@H]1C | 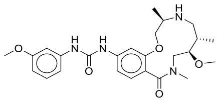 | 35.0 | 35.0 | 1.00 |
| BRD-K50710722-001-01-3 | | CCCN1C[C@H](C)[C@@H](CN(C)C(=O)c2cc(NC(=O)COC)ccc2OC[C@H]1C)OC | 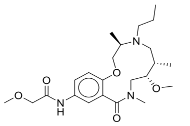 | 35.0 | 35.0 | 1.00 |
| BRD-K54148510-001-01-8 | | C[C@@H](CO)N1C[C@H](C)[C@H](CN(C)Cc2ccc3OCOc3c2)Oc2ccc(NC(=O)CCC(F)(F)F)cc2C1=O | 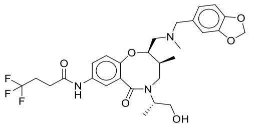 | 35.0 | 35.0 | 1.00 |
| BRD-K68074655-001-01-5 | | C[C@@H](CO)N1C[C@@H](C)[C@H](CN(C)S(=O)(=O)c2ccc(F)cc2)Oc2ccc(NC(=O)CCN3CCOCC3)cc2C1=O | 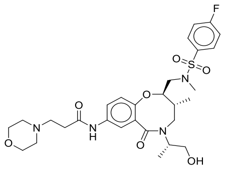 | 35.0 | 35.0 | 1.00 |
| BRD-K68303333-001-01-8 | | C[C@H](CO)N1C[C@@H](C)[C@H](CN(C)C)OCCCC[C@@H](C)Oc2ccc(NC(=O)c3ccncc3)cc2C1=O | 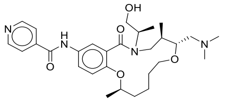 | 35.0 | 35.0 | 1.00 |
| BRD-K80298065-001-01-8 | | CC(C)NC(=O)N(C)C[C@H]1Oc2ccc(NS(=O)(=O)c3ccc(C)cc3)cc2C(=O)N(C[C@H]1C)[C@@H](C)CO | 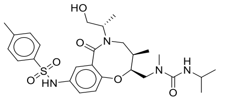 | 35.0 | 35.0 | 1.00 |

| | | | | | | | |
|---|---|---|---|---|---|---|---|
| BRD-K84480302-001-01-1 | | C[C@@H](CO)N1C[C@H](C)[C@@H](CN(C)CC2CC2)Oc2ccc(NS(=O)(=O)c3cn(C)cn3)cc2C1=O | 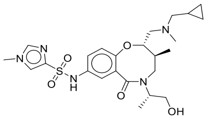 | 35.0 | 35.0 | 1.00 | |
| BRD-K88156935-001-01-8 | | C[C@@H](CO)N1C[C@H](C)[C@H](CN(C)C(=O)Nc2ccc(F)cc2)Oc2ccc(cc2C1=O)N(C)C | 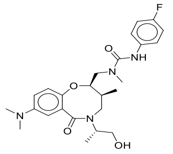 | 35.0 | 35.0 | 1.00 | |
| BRD-K93051667-001-01-8 | | CO[C@@H]1CN(C)C(=O)c2ccc(NS(=O)(=O)c3cn(C)cn3)cc2OC[C@H](C)N(Cc2cccc(F)c2)C[C@@H]1C | 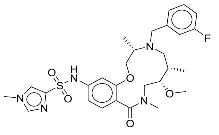 | 35.0 | 35.0 | 1.00 | |
| BRD-K99195544-001-01-6 | | CCC(=O)Nc1ccc2c(OC[C@H](C)N(C[C@H](C)[C@@H](CN(C)C2=O)OC)C(=O)CN2CCOCC2)c1 | 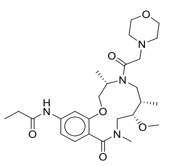 | 35.0 | 35.0 | 1.00 | |
| BRD-A01636364-003-08-6 | Bupivacane hydrochloride | CCCCN1CCCCC1C(=O)Nc1c(C)cccc1C | 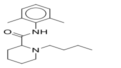 | 35.0 | 35.0 | 1.00 | |
| BRD-A31916785-103-01-0 | Timolol maleate | CC(C)(C)NCC(O)COc1nsnc1N1CCOCC1 | 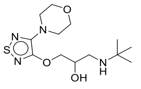 | 35.0 | 35.0 | 1.00 | |
| BRD-K00662280-001-01-1 | CL 218872 | Cc1nnc2ccc(nn12)-c1cccc(c1)C(F)(F)F | 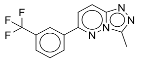 | 35.0 | 11.0 | 0.31 | borderline active GSK3b |
| BRD-K30984264-001-06-2 | Harmine | COc1ccc2c(c1)[nH]c1c(C)nccc21 | 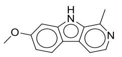 | 35.0 | 35.0 | 1.00 | |
| BRD-K37270826-001-03-7 | Mifepristone | CC#C[C@]1(O)CC[C@H]2[C@@H]3CCC4=CC(=O)CCC4=C3[C@H](C[C@]12C)c1ccc(cc1)N(C)C | 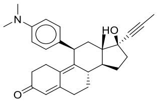 | 35.0 | 35.0 | 1.00 | |
| BRD-K50388907-001-15-5 | Fenofibrate | CC(C)OC(=O)C(C)(C)Oc1ccc(cc1)C(=O)c1ccc(Cl)cc1 | 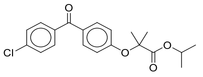 | 35.0 | 35.0 | 1.00 | |
| BRD-K59419204-001-01-9 | AM 281 | Cc1c(nn(c1-c1ccc(I)cc1)-c1ccc(Cl)cc1Cl)C(=O)NN1CCOCC1 | 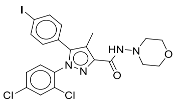 | 35.0 | 35.0 | 1.00 | |
| BRD-K79684402-300-01-0 | Ro 10-5824 | Cc1ncc(CN2CCC(=CC2)c2ccccc2)c(N)n1 | 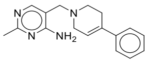 | 35.0 | 35.0 | 1.00 | |
| BRD-K84358317-001-01-2 | | CC(C)C#Cc1ccc2c(O[C@H](CN(C)C(=O)c3ccncc3)[C@H](C)CN([C@H](C)CO)S2(=O)=O)c1 | 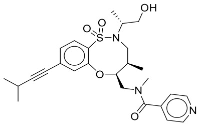 | 35.0 | 35.0 | 1.00 | |
| BRD-K97564742-103-01-9 | Mepyramine maleate | COc1ccc(cc1)CN(CCN(C)C)c1ccccn1 | 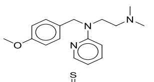 | 35.0 | 35.0 | 1.00 | |
| BRD-K33710385-001-05-4 | Ethionamide | CCc1cc(ccn1)C(N)=S | 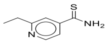 | 38.0 | 38.0 | 1.00 | |

| | | | | | | |
|---|---|---|---|---|---|---|
| BRD-K90524085-001-05-2 | MY-5445 | Clc1cccc(Nc2nnc(-c3ccccc3)c3ccccc23)c1 | 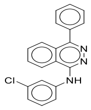 | 50.0 | 33.8 | 0.68 |
| BRD-K97181089-001-02-7 | Amiloride | NC(=N)NC(=O)c1nc(Cl)c(N)nc1N | 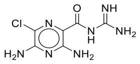 | 75.0 | 75.0 | 1.00 |

**Supplementary Table 3: PGC-1α-related compound structures, whose behavior is shown in Figure 3c and Supplementary Figures 11 and 12**

| Compound ID | Structure | Name |
|---|---|---|
| BRD-K67298865 | 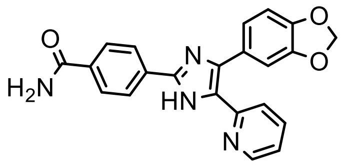 | 4-(4-(benzo[d][1,3]dioxol-5-yl)-5-(pyridin-2-yl)-1H-imidazol-2-yl)benzamide |
| BRD-K67831364 | 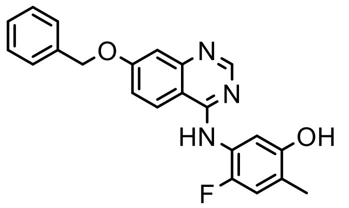 | 5-((7-(benzyloxy)quinazolin-4-yl)amino)-4-fluoro-2-methylphenol |
| BRD-K93258693 | 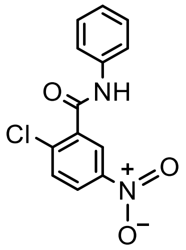 | 2-chloro-5-nitro-N-phenylbenzamide |
| BRD-K63150726 | 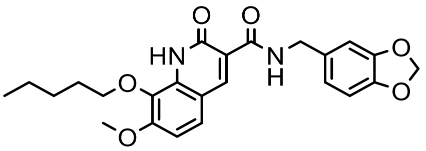 | n-(1,3-benzodioxol-5-ylmethyl)-1,2-dihydro-7-methoxy-2-oxo-8-(pentyloxy)-3-quinolinecarboxamide |
| BRD-K19534880 | 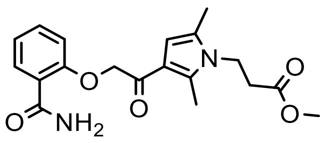 | methyl 3-(3-(2-(2-carbamoylphenoxy)acetyl)-2,5-dimethyl-1H-pyrrol-1-yl)propanoate |
| BRD-K65503129 | 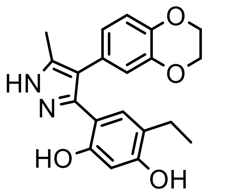 | 4-[4-(2,3-dihydro-1,4-benzodioxin-6-yl)-5-methyl-1H-pyrazol-3-yl]-6-ethylbenzene-1,3-diol |

| | | |
|---|---|---|
| BRD-K68552125 | 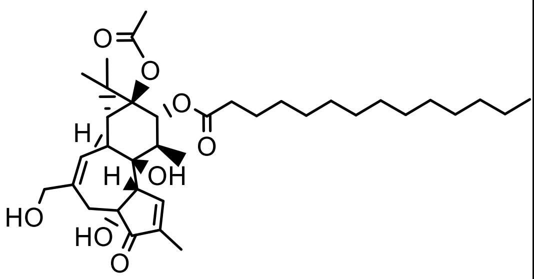 | 12-O-Tetradecanoylphorbol-13-acetate |
| BRD-K95785537 | 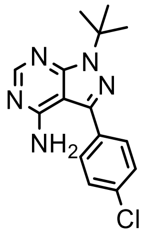 | 1-(tert-butyl)-3-(4-chlorophenyl)-1H-pyrazolo[3,4-d]pyrimidin-4-amine |
| BRD-K29542628 | 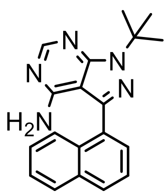 | 1-(tert-Butyl)-3-(naphthalen-1-yl)-1H-pyrazolo[3,4-d]pyrimidin-4-amine |
| BRD-K06234293 | 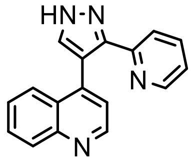 | 4-(3-(pyridin-2-yl)-1H-pyrazol-4-yl)quinoline |
| BRD-K02862004 | 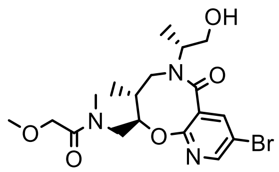 | *N*-[[(2*R*,3*R*)-8-bromo-5-[(2*R*)-1-hydroxypropan-2-yl]-3-methyl-6-oxo-3,4-dihydro-2*H*-pyrido[2,3-b][1,5]oxazocin-2-yl]methyl]-2-methoxy- *N*-methylacetamide |
| BRD-A70407468 | 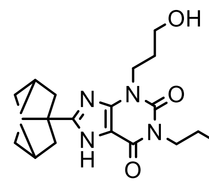 | 1-butyl-3-(3-hydroxypropyl)-8-(3-tricyclo[3.3.1.$0^{3,7}$]nonanyl)-7*H*-purine-2,6-dione |
| BRD-K74094800 | 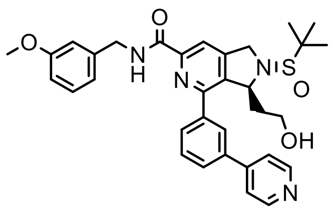 | (3*S*)-2-[(*S*)-*tert*-butylsulfinyl]-3-(2-hydroxyethyl)-*N*-[(3-methoxyphenyl)methyl]-4- (3-pyridin-4-ylphenyl)-1,3-dihydropyrrolo[3,4-c]pyridine-6-carboxamide |

| | | |
|---|---|---|
| BRD-K69705756 | 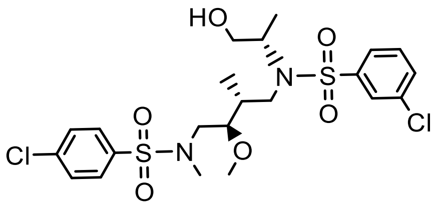 | 3-chloro-N-((2R,3R)-4-((4-chloro-N-methylphenyl)sulfonamido)-3-methoxy-2-methylbutyl)-N-((S)-1-hydroxypropan-2-yl)benzenesulfonamide |
| BRD-K17133642 | 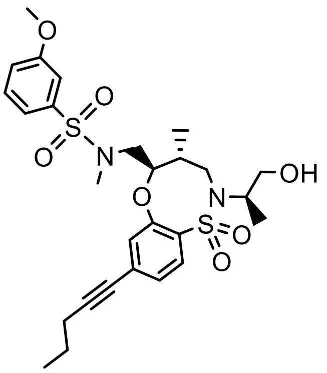 | N-(((4R,5R)-2-((R)-1-hydroxypropan-2-yl)-4-methyl-1,1-dioxido-8-(pent-1-yn-1-yl)-2,3,4,5-tetrahydrobenzo[b][1,4,5]oxathiazocin-5-yl)methyl)-3-methoxy-N-methylbenzenesulfonamide |
| BRD-K10449938 | 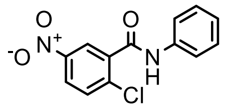 | 3-chloro-*N*-[(2*R*,3*R*)-4-[(4-chlorophenyl)sulfonyl-methylamino]-3-methoxy-2-methylbutyl]-*N*-[(2*R*)-1-hydroxypropan-2-yl]benzenesulfonamide |
| BRD-K35458079 | 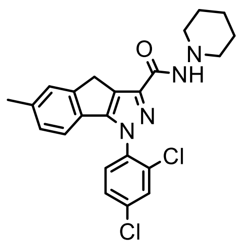 | 5-methyl-2-phenyl-4*H*-pyrazol-3-one |
| BRD-K65285700 | 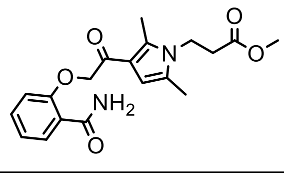 | 1-(2,4-dichlorophenyl)-6-methyl-*N*-piperidin-1-yl-4*H*-indeno[1,2-c] pyrazole-3-carboxamide |
| BRD-K19534880 | 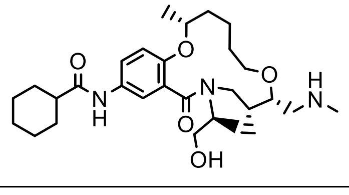 | methyl 3-[3-[2-(2-carbamoylphenoxy)acetyl]-2,5-dimethylpyrrol-1-yl]propanoate |
| BRD-K45142472 | 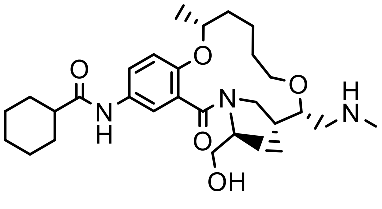 | *N*-[(3*R*,9*S*,10*R*)-12-[(2*S*)-1-hydroxypropan-2-yl]-3,10-dimethyl-9-(methylaminomethyl)-13-oxo-2,8-dioxa-12-azabicyclo[12.4.0]octadeca- |

| | | |
|---|---|---|
| | | 1(14),15,17-trien-16-yl]cyclohexanecarboxamide |
| BRD-K68223954 | 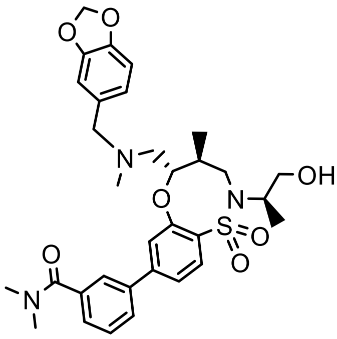 | 3-((4S,5S)-5-(((benzo[d][1,3]dioxol-5-ylmethyl)(methyl)amino)methyl)-2-((R)-1-hydroxypropan-2-yl)-4-methyl-1,1-dioxido-2,3,4,5-tetrahydrobenzo[b][1,4,5]oxathiazocin-8-yl)-N,N-dimethylbenzamide |
| BRD-K14309706 | 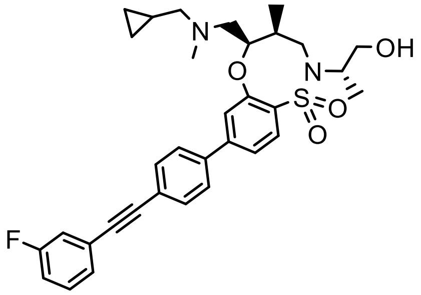 | (4S,5R)-5-(((cyclopropylmethyl)(methyl)amino) methyl)-8-(4-((3-fluorophenyl) ethynyl)phenyl)-2-((S)-1-hydroxypropan-2-yl)-4-methyl-2,3,4,5-tetrahydrobenzo[b][1,4,5] oxathiazocine 1,1-dioxide |
| BRD-K43556160 | 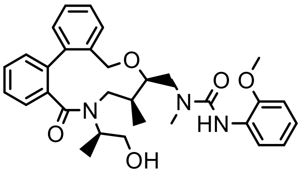 | 1-[[(10*R*,11*S*)-13-[(2*R*)-1-hydroxypropan-2-yl]-11- methyl-14-oxo-9-oxa-13-azatricyclo[13.4.0.0$^{2,7}$] nonadeca-1(19),2,4,6,15,17-hexaen-10-yl]methyl]-3-(2-methoxyphenyl)-1-methylurea |
| BRD-K59605310 | 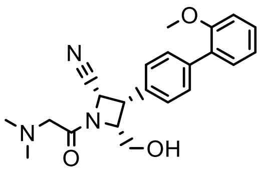 | (2*S*,3*S*,4*R*)-1-[2-(dimethylamino)acetyl]-4-(hydroxymethyl)-3-[4-(2-methoxyphenyl)phenyl] azetidine-2-carbonitrile |
| T0070907 (control) | 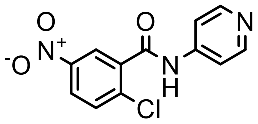 | 2-chloro-5-nitro-N-(pyridin-4-yl)benzamide |

**Supplementary Table 4: YAP1-related compound structures**

| Compound ID | Structure | Name |
|---|---|---|
| BRD-K96698997 (Cmpd. 1) | 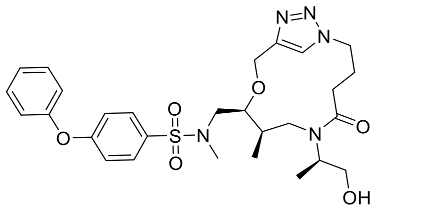 | N-[[(8R,9S)-6-[(2R)-1-hydroxypropan-2-yl]-8-methyl-5-oxo-10-oxa-1,6,13,14-tetrazabicyclo[10.2.1]pentadeca-12(15),13-dien-9-yl]methyl]-N-methyl-4-phenoxybenzenesulfonamide |
| BRD-K13719685 (Cmpd. 2) | 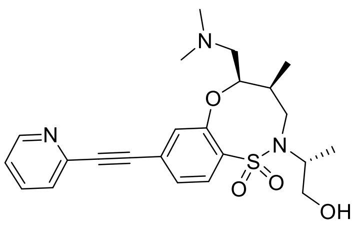 | (4S,5R)-5-((dimethylamino)methyl)-2-((R)-1-hydroxypropan-2-yl)-4-methyl-8-(pyridin-2-ylethynyl)-2,3,4,5-tetrahydrobenzo[b][1,4,5]oxathiazocine 1,1-dioxide |
| **BRD-K34692511 (Cmpd. 3)** | 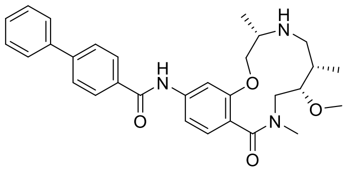 | N-[(4S,7S,8S)-8-methoxy-4,7,10-trimethyl-11-oxo-2-oxa-5,10-diazabicyclo[10.4.0]hexadeca-1(12),13,15-trien-15-yl]-4-phenylbenzamide |
| **BRD-K28862419 (Cmpd. 4)** | 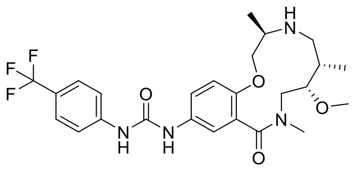 | 1-[(4R,7S,8S)-8-methoxy-4,7,10-trimethyl-11-oxo-2-oxa-5,10-diazabicyclo[10.4.0]hexadeca-1(12),13,15-trien-14-yl]-3-[4-(trifluoromethyl)phenyl]urea |
| BRD-K70003473 (Cmpd. 5) | 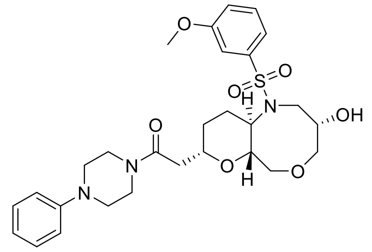 | 2-[(3S,6aR,8S,10aR)-3-hydroxy-1-(3-methoxyphenyl)sulfonyl-3,4,6,6a,8,9,10,10a-octahydro-2H-pyrano[2,3-c][1,5]oxazocin-8-yl]-1-(4-phenyl-1-piperazinyl)ethanone |
| BRD-K46678324 (Cmpd. 6) | 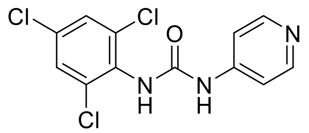 | 1-pyridin-4-yl-3-(2,4,6-trichlorophenyl)urea |

| | | |
|---|---|---|
| BRD-K06593056 (Cmpd. 7) | 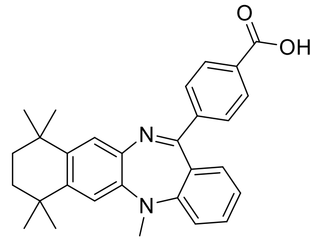 | 4-(5,7,7,10,10-pentamethyl-8,9-dihydronaphtho[2,3-b][1,4]benzodiazepin-13-yl)benzoic acid |
| BRD-A61154809 (Cmpd. 8) | 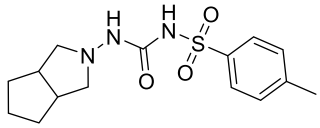 | 1-(3,3a,4,5,6,6a-hexahydro-1H-cyclopenta[c]pyrrol-2-yl)-3-(4-methylphenyl)sulfonylurea |
| BRD-K77793136 (Cmpd. 9) | 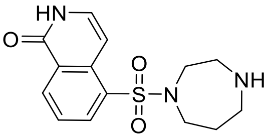 | 5-(1,4-diazepan-1-ylsulfonyl)-2H-isoquinolin-1-one |
| BRD-K15567136 (Cmpd. 10) | 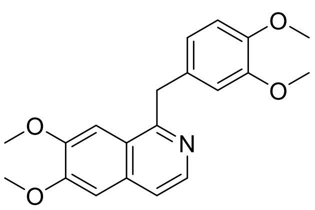 | 1-[(3,4-dimethoxyphenyl)methyl]-6,7-dimethoxyisoquinoline |
| BRD-K88429204 (Cmpd. 11) | 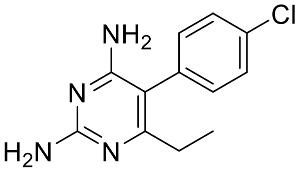 | 5-(4-chlorophenyl)-6-ethylpyrimidine-2,4-diamine |
| BRD-K42095107 (Cmpd. 12) | 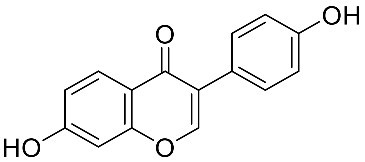 | 7-hydroxy-3-(4-hydroxyphenyl)chromen-4-one |
| **BRD-K43796186 (Cmpd. 13)** | 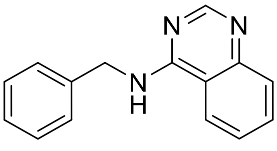 | N-benzylquinazolin-4-amine |
| BRD-K37451830 (Cmpd. 14) | 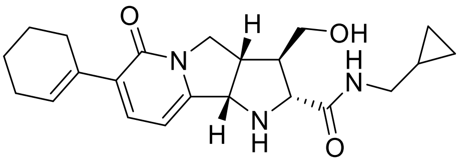 | (2R,3R,3aS,9bS)-7-(1-cyclohexenyl)-N-(cyclopropylmethyl)-3-(hydroxymethyl)-6-oxo-1,2,3,3a,4,9b-hexahydropyrrolo[2,3-a]indolizine-2-carboxamide |

| | | |
|---|---|---|
| BRD-K03953354 (Cmpd. 15) | 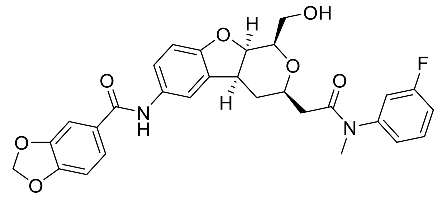 | N-[(1R,3R,4aS,9aR)-3-[2-[(3-fluorophenyl)methylamino]-2-oxoethyl]-1-(hydroxymethyl)-3,4,4a,9a-tetrahydro-1H-pyrano[3,4-b]benzofuran-6-yl]-1,3-benzodioxole-5-carboxamide |
| BRD-K39839146 (Cmpd. 16) | 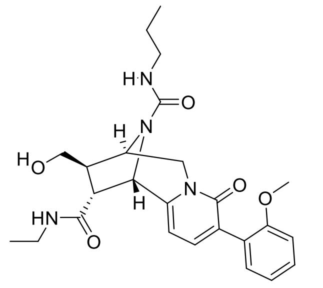 | (1S,9R,10R,11R)-11-N-ethyl-10-(hydroxymethyl)-5-(2-methoxyphenyl)-6-oxo-12-N-propyl-7,12-diazatricyclo[7.2.1.02,7]dodeca-2,4-diene-11,12-dicarboxamide |
| BRD-K62768599 (Cmpd. 17) | 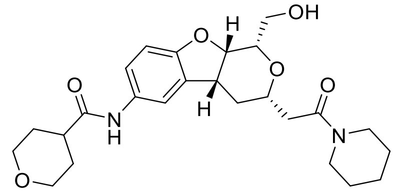 | N-[(1S,3S,4aR,9aS)-1-(hydroxymethyl)-3-[2-oxo-2-(1-piperidinyl)ethyl]-3,4,4a,9a-tetrahydro-1H-pyrano[3,4-b]benzofuran-6-yl]-4-oxanecarboxamide |
| BRD-K42367391 (Cmpd. 18) | 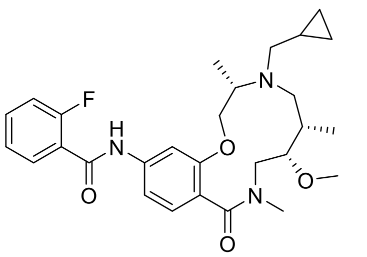 | N-[(5S,6S,9S)-8-(cyclopropylmethyl)-5-methoxy-3,6,9-trimethyl-2-oxo-11-oxa-3,8-diazabicyclo[10.4.0]hexadeca-1(12),13,15-trien-14-yl]-2-fluorobenzamide |
| BRD-K22874335 (Cmpd. 19) | 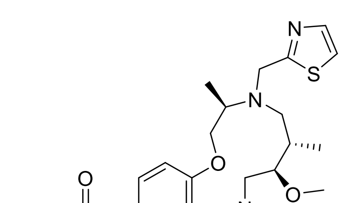 | N-[(4R,7S,8R)-8-methoxy-4,7,10-trimethyl-11-oxo-5-(1,3-thiazol-2-ylmethyl)-2-oxa-5,10-diazabicyclo[10.4.0]hexadeca-1(12),13,15-trien-14-yl]cyclohexanecarboxamide |
| BRD-K41723088 (Cmpd. 20) | 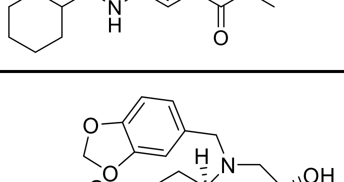 | 2-[(3S,6aR,8R,10aR)-1-(1,3-benzodioxol-5-ylmethyl)-3-hydroxy-3,4,6,6a,8,9,10,10a-octahydro-2H-pyrano[2,3-c][1,5]oxazocin-8-yl]-1-piperidin-1-ylethanone |
| BRD-K68530167 (Cmpd. 21) | 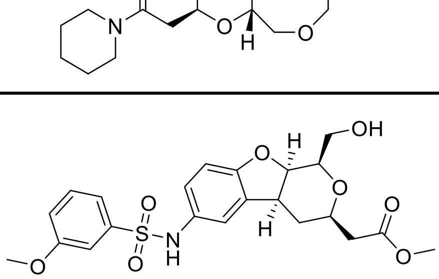 | 2-[(1R,3R,4aS,9aR)-1-(hydroxymethyl)-6-[(3-methoxyphenyl)sulfonylamino]-3,4,4a,9a-tetrahydro-1H-pyrano[3,4- |

| | | |
|---|---|---|
| | | b]benzofuran-3-yl]acetic acid methyl ester |
| BRD-K22754756 (Cmpd. 22) | 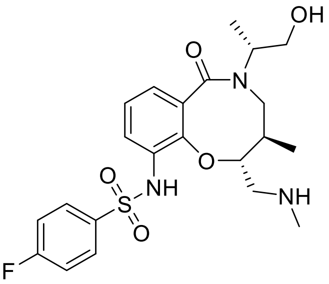 | 4-fluoro-N-[(2R,3R)-5-[(2R)-1-hydroxypropan-2-yl]-3-methyl-2-(methylaminomethyl)-6-oxo-3,4-dihydro-2H-1,5-benzoxazocin-10-yl]benzenesulfonamide |
| BRD-K11266478 (Cmpd. 23) | 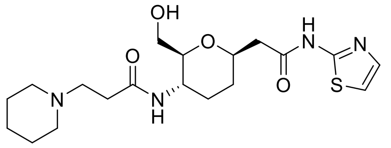 | N-[(2S,3S,6R)-2-(hydroxymethyl)-6-[2-oxo-2-(1,3-thiazol-2-ylamino)ethyl]oxan-3-yl]-3-piperidin-1-ylpropanamide |
| BRD-K00135177 (Cmpd. 24) | 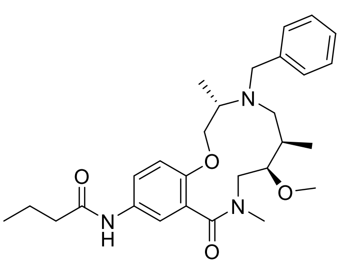 | N-[(4S,7R,8R)-8-methoxy-4,7,10-trimethyl-11-oxo-5-(phenylmethyl)-2-oxa-5,10-diazabicyclo[10.4.0]hexadeca-1(12),13,15-trien-14-yl]butanamide |
| BRD-K40143134 (Cmpd. 25) | 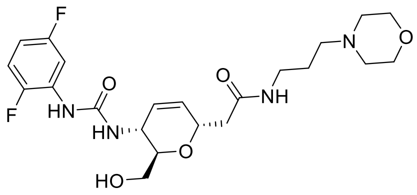 | 2-[(2R,3R,6S)-3-[[(2,5-difluoroanilino)-oxomethyl]amino]-2-(hydroxymethyl)-3,6-dihydro-2H-pyran-6-yl]-N-[3-(4-morpholinyl)propyl]acetamide |
| BRD-K11758216 (Cmpd. 26) | 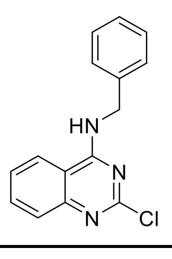 | N-benzyl-2-chloroquinazolin-4-amine |
| BRD-K48052543 (Cmpd. 27) | 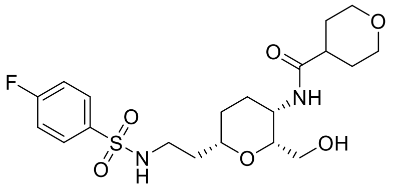 | N-[(2R,3S,6S)-6-[2-[(4-fluorophenyl)sulfonylamino]ethyl]-2-(hydroxymethyl)oxan-3-yl]oxane-4-carboxamide |
| BRD-A50675702 (Cmpd. 28) | 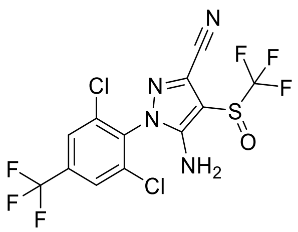 | 5-amino-1-[2,6-dichloro-4-(trifluoromethyl)phenyl]-4-(trifluoromethane)sulfinyl-1H-pyrazole-3-carbonitrile |

| | | |
|---|---|---|
| BRD-K28043081 (Cmpd. 29) | 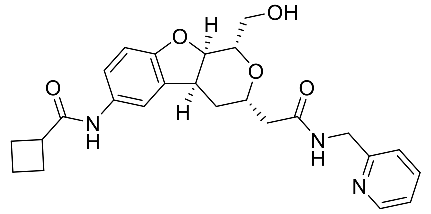 | N-[(1S,3S,4aS,9aR)-1-(hydroxymethyl)-3-[2-oxo-2-(pyridin-2-ylmethylamino)ethyl]-3,4,4a,9a-tetrahydro-1H-pyrano[3,4-b][1]benzofuran-6-yl]cyclobutanecarboxamide |
| BRD-K19969618 (Cmpd. 30) | 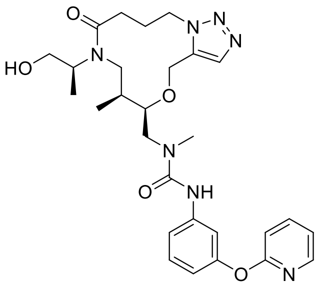 | 1-[[(8S,9R)-6-[(2S)-1-hydroxypropan-2-yl]-8-methyl-5-oxo-10-oxa-1,6,14,15-tetrazabicyclo[10.3.0]pentadeca-12,14-dien-9-yl]methyl]-1-methyl-3-(3-pyridin-2-yloxyphenyl)urea |

**Supplementary Table 5: Thirty compounds whose Cell Painting profile matched (25 compounds) or opposed (5 compounds) the overexpression of the Hippo pathway effector Yes-associated protein 1 (YAP1)**

| | Broad ID | MOA | Compound Name | Known Targets | Corr. to YAP cluster | Avg. Cell Count z-score | Corr. to TRAF2 |
|---|---|---|---|---|---|---|---|
| 1 | BRD-K96698997-001-01-4 | | | | -0.451816097371088 | -1.09876751317736 | 0.483378769177758 |
| 11 | BRD-K88429204-001-04-7 | dihydrofolate reductase inhibitor | pyrimethamine | DHFR, SLC47A1 | 0.450273057324876 | -0.21652845532613 | -0.349609799358579 |
| 9 | BRD-K77793136-003-01-4 | rho associated kinase inhibitor | hydroxyfasudil | PKIA, PRKACA, ROCK1 | 0.445765052349004 | -1.05491865344321 | -0.466745148730574 |
| 5 | BRD-K70003473-001-01-0 | | | | -0.419476423486351 | 0.57275101988869 | 0.309863968937689 |
| 21 | BRD-K68530167-001-02-5 | | | | 0.493953439051083 | -0.141108416583381 | -0.461903561022211 |
| 17 | BRD-K62768599-001-01-7 | | | | 0.488328497910833 | -0.739206863357278 | -0.52723612634195 |
| 27 | BRD-K48052543-001-01-2 | | | | 0.509868432726129 | 0.0980140318335538 | -0.413168132822434 |
| 6 | BRD-K46678324-001-03-7 | rho associated kinase inhibitor | RHO-kinase inhibitor II | | 0.428418602602677 | -0.725175228242348 | -0.339745295430873 |
| 13 | BRD-K43796186-001-01-1 | EGFR inhibitor | benzyl-quinazolin-4-yl-amine | EGFR | 0.462214494435451 | -0.570827241978117 | -0.377843726025219 |
| 18 | BRD-K42367391-001-01-3 | | | | 0.488715954293843 | -0.937403709355667 | -0.272772324093652 |

| | Broad ID | MOA | Compound Name | Known Targets | Corr. to YAP cluster | Avg. Cell Count z-score | Corr. to TRAF2 |
|---|---|---|---|---|---|---|---|
| 12 | BRD-K42095107-001-02-3 | estrogen receptor agonist | daidzein | ESRRA, ESRRB, ESRRG, TRPC5 | 0.458309143762898 | -0.881277168895946 | -0.371958320674078 |
| 20 | BRD-K41723088-001-01-6 | | | | 0.491645117137496 | -0.0709502410087301 | -0.43890527237543 |
| 25 | BRD-K40143134-001-01-2 | | | | 0.50094854898695 | -0.114799100742887 | -0.342751435013124 |
| 16 | BRD-K39839146-001-01-4 | | | | 0.483549276460743 | 0.218452233236704 | -0.428362405612274 |
| 14 | BRD-K37451830-001-01-1 | | | | 0.480168234528875 | -0.00429997421281192 | -0.416740972979351 |
| 3 | BRD-K34692511-001-01-9 | | | | -0.439820984888043 | 0.478037482862912 | 0.286882376707942 |
| 4 | BRD-K28862419-001-01-9 | | | | -0.431674202259506 | -2.04414892904578 | 0.406783833287405 |
| 29 | BRD-K28043081-001-01-3 | | | | 0.526963070335337 | -0.553287698084454 | -0.406037778061057 |
| 19 | BRD-K22874335-001-01-1 | | | | 0.490400094569485 | -1.70388177750873 | -0.362018046441038 |
| 22 | BRD-K22754756-001-01-5 | | | | 0.496607051682064 | -1.40921744009519 | -0.297027575600364 |
| 30 | BRD-K19969618-001-01-2 | | | | 0.548319649662317 | 0.77971763783391 | -0.466667305651416 |

| | Broad ID | MOA | Compound Name | Known Targets | Corr. to YAP cluster | Avg. Cell Count z-score | Corr. to TRAF2 |
|---|---|---|---|---|---|---|---|
| 10 | BRD-K15567136-001-01-1 | phosphodiesterase inhibitor | papaverine hydrochloride | PDE10A, PDE4B, PDE5A | 0.44641427678599 | -0.656771007057064 | -0.493557215874781 |
| 2 | BRD-K13719685-001-01-5 | | | | -0.442645220271481 | 2.10395320180544 | 0.359126741534968 |
| 26 | BRD-K11758216-001-01-3 | | | | 0.507801351616454 | -1.20225082214997 | -0.440615345966013 |
| 23 | BRD-K11266478-001-01-0 | | | | 0.498791551452305 | 0.485053300420377 | -0.439765555665776 |
| 7 | BRD-K06593056-001-01-4 | retinoid receptor agonist | LE-135 | RARB | 0.435680365509664 | -0.790948517843583 | -0.451237141338472 |
| 15 | BRD-K03953354-001-01-2 | | | | 0.483314867234996 | -1.39693975936963 | -0.481607208500513 |
| 24 | BRD-K00135177-001-01-4 | | | | 0.500357831891246 | 0.400863489730796 | -0.364867410280201 |
| 8 | BRD-A61154809-001-03-5 | ATP channel blocker\|insulin secretagogue | gliclazide | ABCC8, VEGFA | 0.440145834465088 | 0.820309153702101 | -0.33133766243123 |
| 28 | BRD-A50675702-001-03-0 | chloride channel blocker\|GABA gated chloride channel blocker | fipronil | | 0.51019424419505 | -0.118808139347153 | -0.475169203024327 |

### Supplementary Table 6: RNA-sequencing-based enrichment analysis of Hallmark gene sets up- and down-regulated in KP230 cells by NB4A

| NAME | GS<br> follow link to MSigDB | GS DETAILS | SIZE | ES | NES | NOM p-val | FDR q-val | FWER p-val | RANK AT MAX | LEADING EDGE | UPREGULATED OR DOWNREGULATED |
|---|---|---|---|---|---|---|---|---|---|---|---|
| HALLMARK_INTERFERON_ALPHA_RESPONSE | HALLMARK_INTERFERON_ALPHA_RESPONSE | Details ... | 86 | 0.5941053 | 2.2873087 | 0 | 0 | 0 | 1819 | tags=38%, list=12%, signal=43% | Upregulated in NB4A-treated |
| HALLMARK_INTERFERON_GAMMA_RESPONSE | HALLMARK_INTERFERON_GAMMA_RESPONSE | Details ... | 166 | 0.3945195 | 1.7322907 | 0 | 0.003686826 | 0.008 | 2131 | tags=29%, list=14%, signal=33% | Upregulated in NB4A-treated |
| HALLMARK_PROTEIN_SECRETION | HALLMARK_PROTEIN_SECRETION | Details ... | 92 | 0.41896117 | 1.6522279 | 0.002403846 | 0.00996725 | 0.032 | 2513 | tags=36%, list=17%, signal=43% | Upregulated in NB4A-treated |
| HALLMARK_MYC_TARGETS_V1 | HALLMARK_MYC_TARGETS_V1 | Details ... | 182 | 0.3305301 | 1.4714037 | 0.005181347 | 0.04932377 | 0.189 | 2181 | tags=26%, list=15%, signal=30% | Upregulated in NB4A-treated |
| HALLMARK_APOPTOSIS | HALLMARK_APOPTOSIS | Details ... | 150 | 0.34260988 | 1.4562898 | 0.002717391 | 0.04559618 | 0.214 | 1928 | tags=27%, list=13%, signal=31% | Upregulated in NB4A-treated |
| HALLMARK_UV_RESPONSE_UP | HALLMARK_UV_RESPONSE_UP | Details ... | 146 | 0.30856675 | 1.3382866 | 0.024523161 | 0.11366022 | 0.524 | 3312 | tags=36%, list=22%, signal=46% | Upregulated in NB4A-treated |
| HALLMARK_COMPLEMENT | HALLMARK_COMPLEMENT | Details ... | 158 | 0.29993588 | 1.3087488 | 0.036931816 | 0.12590155 | 0.617 | 2019 | tags=20%, list=13%, signal=23% | Upregulated in NB4A-treated |
| HALLMARK_DNA_REPAIR | HALLMARK_DNA_REPAIR | Details ... | 143 | 0.3008566 | 1.2843143 | 0.035326086 | 0.13475496 | 0.689 | 3426 | tags=40%, list=23%, signal=51% | Upregulated in NB4A-treated |
| HALLMARK_PI3K_AKT_MTOR_SIGNALING | HALLMARK_PI3K_AKT_MTOR_SIGNALING | Details ... | 97 | 0.29323217 | 1.168757 | 0.15869017 | 0.30128673 | 0.948 | 2927 | tags=32%, list=20%, signal=39% | Upregulated in NB4A-treated |
| HALLMARK_HEME_METABOLISM | HALLMARK_HEME_METABOLISM | Details ... | 169 | 0.26993614 | 1.1590812 | 0.13533835 | 0.2913993 | 0.958 | 2760 | tags=24%, list=18%, signal=29% | Upregulated in NB4A-treated |
| HALLMARK_ALLOGRAFT_REJECTION | HALLMARK_ALLOGRAFT_REJECTION | Details ... | 149 | 0.26266745 | 1.1286 | 0.17435898 | 0.33098337 | 0.984 | 2168 | tags=19%, list=14%, signal=22% | Upregulated in NB4A-treated |
| HALLMARK_OXIDATIVE_PHOSPHORYLATION | HALLMARK_OXIDATIVE_PHOSPHORYLATION | Details ... | 192 | 0.2433627 | 1.0804513 | 0.26944444 | 0.42189047 | 0.999 | 3242 | tags=30%, list=22%, signal=38% | Upregulated in NB4A-treated |
| HALLMARK_REACTIVE_OXYGEN_SPECIES_PATHWAY | HALLMARK_REACTIVE_OXYGEN_SPECIES_PATHWAY | Details ... | 46 | 0.30948043 | 1.072884 | 0.35746607 | 0.4090726 | 0.999 | 2204 | tags=26%, list=15%, signal=31% | Upregulated in NB4A-treated |
| HALLMARK_IL2_STAT5_SIGNALING | HALLMARK_IL2_STAT5_SIGNALING | Details ... | 178 | 0.2425146 | 1.0618333 | 0.30357143 | 0.40563384 | 1 | 2168 | tags=21%, list=14%, signal=24% | Upregulated in NB4A-treated |
| HALLMARK_IL6_JAK_STAT3_SIGNALING | HALLMARK_IL6_JAK_STAT3_SIGNALING | Details ... | 75 | 0.27034676 | 1.0383954 | 0.378866 | 0.4350973 | 1 | 3312 | tags=31%, list=22%, signal=39% | Upregulated in NB4A-treated |
| HALLMARK_ADIPOGENESIS | HALLMARK_ADIPOGENESIS | Details ... | 189 | 0.23059528 | 1.0254962 | 0.3837535 | 0.4401859 | 1 | 2798 | tags=25%, list=19%, signal=30% | Upregulated in NB4A-treated |
| HALLMARK_PANCREAS_BETA_CELLS | HALLMARK_PANCREAS_BETA_CELLS | Details ... | 25 | 0.33470038 | 1.0112545 | 0.4312115 | 0.4494147 | 1 | 1269 | tags=16%, list=8%, signal=17% | Upregulated in NB4A-treated |
| HALLMARK_SPERMATOGENESIS | HALLMARK_SPERMATOGENESIS | Details ... | 109 | 0.23348868 | 0.9675974 | 0.5630027 | 0.53295434 | 1 | 4079 | tags=32%, list=27%, signal=44% | Upregulated in NB4A-treated |
| HALLMARK_EPITHELIAL_MESENCHYMAL_TRANSITION | HALLMARK_EPITHELIAL_MESENCHYMAL_TRANSITION | Details ... | 190 | -0.6542706 | -2.734083 | 0 | 0 | 0 | 1536 | tags=43%, list=10%, signal=47% | Downregulated in NB4A-treated |
| HALLMARK_ANGIOGENESIS | HALLMARK_ANGIOGENESIS | Details ... | 31 | -0.6635637 | -2.009867 | 0 | 0 | 0 | 1291 | tags=35%, list=9%, signal=39% | Downregulated in NB4A-treated |
| HALLMARK_GLYCOLYSIS | HALLMARK_GLYCOLYSIS | Details ... | 184 | -0.4841484 | -2.004427 | 0 | 0 | 0 | 1519 | tags=28%, list=10%, signal=31% | Downregulated in NB4A-treated |
| HALLMARK_UV_RESPONSE_DN | HALLMARK_UV_RESPONSE_DN | Details ... | 137 | -0.504841 | -2.002119 | 0 | 0 | 0 | 2807 | tags=40%, list=19%, signal=49% | Downregulated in NB4A-treated |
| HALLMARK_HYPOXIA | HALLMARK_HYPOXIA | Details ... | 186 | -0.4647971 | -1.924311 | 0 | 1.94E-04 | 0.001 | 2715 | tags=40%, list=18%, signal=48% | Downregulated in NB4A-treated |
| HALLMARK_APICAL_JUNCTION | HALLMARK_APICAL_JUNCTION | Details ... | 185 | -0.421712 | -1.741695 | 0 | 0.005009011 | 0.029 | 2672 | tags=32%, list=18%, signal=39% | Downregulated in NB4A-treated |
| HALLMARK_TGF_BETA_SIGNALING | HALLMARK_TGF_BETA_SIGNALING | Details ... | 54 | -0.5039986 | -1.720936 | 0.003412969 | 0.005356684 | 0.036 | 1462 | tags=31%, list=10%, signal=35% | Downregulated in NB4A-treated |
| HALLMARK_HEDGEHOG_SIGNALING | HALLMARK_HEDGEHOG_SIGNALING | Details ... | 34 | -0.5514501 | -1.700231 | 0.005235602 | 0.005728535 | 0.044 | 3166 | tags=41%, list=21%, signal=52% | Downregulated in NB4A-treated |
| HALLMARK_NOTCH_SIGNALING | HALLMARK_NOTCH_SIGNALING | Details ... | 31 | -0.5646099 | -1.685015 | 0.010948905 | 0.006704273 | 0.055 | 3007 | tags=45%, list=20%, signal=56% | Downregulated in NB4A-treated |
| HALLMARK_ESTROGEN_RESPONSE_EARLY | HALLMARK_ESTROGEN_RESPONSE_EARLY | Details ... | 184 | -0.3867249 | -1.59521 | 0.00155521 | 0.015935246 | 0.142 | 1426 | tags=20%, list=10%, signal=22% | Downregulated in NB4A-treated |
| HALLMARK_WNT_BETA_CATENIN_SIGNALING | HALLMARK_WNT_BETA_CATENIN_SIGNALING | Details ... | 38 | -0.4959761 | -1.583363 | 0.018656716 | 0.015919015 | 0.151 | 3007 | tags=42%, list=20%, signal=53% | Downregulated in NB4A-treated |
| HALLMARK_MTORC1_SIGNALING | HALLMARK_MTORC1_SIGNALING | Details ... | 188 | -0.3639692 | -1.516257 | 0.00312989 | 0.026771687 | 0.269 | 1382 | tags=21%, list=9%, signal=23% | Downregulated in NB4A-treated |
| HALLMARK_MYOGENESIS | HALLMARK_MYOGENESIS | Details ... | 186 | -0.3648306 | -1.504344 | 0.001557632 | 0.02799793 | 0.299 | 1336 | tags=18%, list=9%, signal=20% | Downregulated in NB4A-treated |
| HALLMARK_CHOLESTEROL_HOMEOSTASIS | HALLMARK_CHOLESTEROL_HOMEOSTASIS | Details ... | 69 | -0.4184438 | -1.493341 | 0.028169014 | 0.029403457 | 0.333 | 1705 | tags=30%, list=11%, signal=34% | Downregulated in NB4A-treated |
| HALLMARK_MITOTIC_SPINDLE | HALLMARK_MITOTIC_SPINDLE | Details ... | 193 | -0.3635059 | -1.488942 | 0.010736196 | 0.028197076 | 0.341 | 3281 | tags=36%, list=22%, signal=45% | Downregulated in NB4A-treated |
| HALLMARK_APICAL_SURFACE | HALLMARK_APICAL_SURFACE | Details ... | 39 | -0.4417995 | -1.415298 | 0.060763888 | 0.049216405 | 0.551 | 2354 | tags=31%, list=16%, signal=36% | Downregulated in NB4A-treated |
| HALLMARK_KRAS_SIGNALING_UP | HALLMARK_KRAS_SIGNALING_UP | Details ... | 181 | -0.3398556 | -1.398792 | 0.017377567 | 0.053476032 | 0.61 | 2296 | tags=23%, list=15%, signal=26% | Downregulated in NB4A-treated |
| HALLMARK_ESTROGEN_RESPONSE_LATE | HALLMARK_ESTROGEN_RESPONSE_LATE | Details ... | 179 | -0.3390946 | -1.382303 | 0.009584664 | 0.05974807 | 0.662 | 1426 | tags=18%, list=10%, signal=20% | Downregulated in NB4A-treated |
| HALLMARK_INFLAMMATORY_RESPONSE | HALLMARK_INFLAMMATORY_RESPONSE | Details ... | 163 | -0.3420927 | -1.378494 | 0.020733653 | 0.05833988 | 0.675 | 1185 | tags=16%, list=8%, signal=17% | Downregulated in NB4A-treated |
| HALLMARK_TNFA_SIGNALING_VIA_NFKB | HALLMARK_TNFA_SIGNALING_VIA_NFKB | Details ... | 188 | -0.3291668 | -1.373827 | 0.013740458 | 0.05704531 | 0.688 | 1189 | tags=20%, list=8%, signal=21% | Downregulated in NB4A-treated |
| HALLMARK_KRAS_SIGNALING_DN | HALLMARK_KRAS_SIGNALING_DN | | 138 | -0.3369457 | -1.336891 | 0.051948052 | 0.07507158 | 0.807 | 2429 | tags=19%, list=16%, signal=22% | Downregulated in NB4A-treated |
| HALLMARK_COAGULATION | HALLMARK_COAGULATION | | 107 | -0.3364556 | -1.28073 | 0.08777969 | 0.11387649 | 0.92 | 2896 | tags=30%, list=19%, signal=37% | Downregulated in NB4A-treated |
| HALLMARK_UNFOLDED_PROTEIN_RESPONSE | HALLMARK_UNFOLDED_PROTEIN_RESPONSE | | 103 | -0.3199706 | -1.226251 | 0.13128039 | 0.16439793 | 0.978 | 2247 | tags=29%, list=15%, signal=34% | Downregulated in NB4A-treated |
| HALLMARK_G2M_CHECKPOINT | HALLMARK_G2M_CHECKPOINT | | 190 | -0.2905551 | -1.219829 | 0.097826086 | 0.16501382 | 0.983 | 3942 | tags=39%, list=26%, signal=52% | Downregulated in NB4A-treated |
| HALLMARK_ANDROGEN_RESPONSE | HALLMARK_ANDROGEN_RESPONSE | | 91 | -0.2986016 | -1.129188 | 0.21680672 | 0.2863506 | 0.999 | 1382 | tags=19%, list=9%, signal=20% | Downregulated in NB4A-treated |
| HALLMARK_P53_PATHWAY | HALLMARK_P53_PATHWAY | | 187 | -0.2723642 | -1.106566 | 0.25832012 | 0.3154989 | 0.999 | 1024 | tags=14%, list=7%, signal=15% | Downregulated in NB4A-treated |
| HALLMARK_XENOBIOTIC_METABOLISM | HALLMARK_XENOBIOTIC_METABOLISM | | 169 | -0.2588365 | -1.063456 | 0.31388012 | 0.38838965 | 0.999 | 1382 | tags=14%, list=9%, signal=15% | Downregulated in NB4A-treated |
| HALLMARK_MYC_TARGETS_V2 | HALLMARK_MYC_TARGETS_V2 | | 54 | -0.2797416 | -0.968363 | 0.4856661 | 0.59060425 | 1 | 3804 | tags=37%, list=25%, signal=49% | Downregulated in NB4A-treated |
| HALLMARK_FATTY_ACID_METABOLISM | HALLMARK_FATTY_ACID_METABOLISM | | 148 | -0.2398035 | -0.964624 | 0.53543305 | 0.58000576 | 1 | 2867 | tags=24%, list=19%, signal=29% | Downregulated in NB4A-treated |
| HALLMARK_E2F_TARGETS | HALLMARK_E2F_TARGETS | | 192 | -0.2091263 | -0.868751 | 0.8031496 | 0.79996806 | 1 | 2856 | tags=25%, list=19%, signal=31% | Downregulated in NB4A-treated |
| HALLMARK_PEROXISOME | HALLMARK_PEROXISOME | | 90 | -0.2161393 | -0.804066 | 0.84477127 | 0.902477 | 1 | 1405 | tags=12%, list=9%, signal=13% | Downregulated in NB4A-treated |
| HALLMARK_BILE_ACID_METABOLISM | HALLMARK_BILE_ACID_METABOLISM | | 98 | -0.2020332 | -0.754092 | 0.93811077 | 0.9386531 | 1 | 1464 | tags=13%, list=10%, signal=15% | Downregulated in NB4A-treated |

**Supplementary Table 7: RT-qPCR primer sequences used in the study.**

| Gene name | Forward primer | Reverse primer |
|---|---|---|
| *GAPDH* | GTGGTCTCCTCTGACTTCAAC | CCTGTTGCTGTAGCCAAATTC |
| *YAP1* | GCTGCCACCAAGCTAGATAA | GTGCATGTGTCTCCTTAGATCC |
| *CTGF* | GTGCATCCGTACTCCCAAA | CTCCACAGAATTTAGCTCGGTAT |
| *CYR61* | AGCCTCGCATCCTATACAACC | TTCTTTCACAAGGCGGCACTC |
| *Yap1* | GATGTCTCAGGAATTGAGAAC | CTGTATCCATTTCATCCACAC |
| *Cyr61* | CTGCGCTAAACAACTCAACGA | GCAGATCCCTTTCAGAGCGG |
