## Supplementary material for "Virtual screening for small molecule pathway regulators by image profile matching": Supp Methods and Figures

**This PDF file includes:**

Materials and Methods
Figs. S1 to S15
Tables S1 to S7

### Materials and Methods

##### *Data availability*

The large-scale Cell Painting datasets used in this paper are publicly available and their details and locations are described in publications (gene overexpression dataset[1] and compound dataset[2]). RNA-sequencing data have been deposited into the NCBI Gene Expression Omnibus (GEO; accession number GSE198909).

##### *Code availability*

The code used in this study is available at https://github.com/broadinstitute/GeneCompoundImaging. It is available for use under the BSD 3-clause license, a permissive open-source license.

##### *Cell line and DNA construct availability*

Cell lines and DNA constructs are available from the laboratories that performed the experiments using them, or where restricted by licensing, from commercial sources.

##### *Research animals*

*Boerckel lab:* Mouse experiments were conducted in compliance with all relevant regulations. All animal experiments were performed at the University of Pennsylvania under IACUC review and in compliance with IACUC protocol #806482.

##### *Feature set alignment*

As the two experiments (gene overexpression and small molecule treatmen6=t) were analyzed by a slightly different CellProfiler pipeline, and also the phenotype of the negative controls are quite different (Figure 1b), an extra data preprocessing step is needed to make the feature sets comparable. To achieve this, we first took the intersection of features in the two datasets, which resulted in 605 features (1399 features in the genetic screen, without any feature selection; and 729 features in the compound screen, obtained using the findCorrelation with threshold of 0.90 on the original 1,783 dimensional feature set). In order to compare values of the corresponding features across experiments, each feature is standardized (mean-centered and scaled by standard deviation) with respect to the negative control. This was done platewise based on the mean and standard deviation of the controls at profile level for the compound dataset. The normalization parameters are slightly different for the genetic screen, where median and median absolute deviation (MAD) are used instead, to remove outlier effects[1]. The code repository for all the analyses are publicly available as described in Code Availability.

##### *Compound annotations*

Compound MOAs and target annotations were mainly acquired from the "Repurposing hub" [3] and then curated to include missing annotations from other sources, such as DrugBank [4]. This results in 747 compounds annotated with the gene(s) that they target. The protein interaction data, which was used to assess relevance of a protein to compound targets, was collected from BioGRID [5]. We note that 98.6% of BioGrid human gene annotations are based on physical interaction rather than genetic (Oughtred et al. 2021). For the Gene Ontology-based functional similarity, we used the R package ontologyIndex and ontologySimilarity (Greene et al. 2017), which were previously developed to calculate semantic similarity between ontological objects. The similarity score is calculated based on the GO annotations under biological processes through the get_sim_grid function.

***Scoring gene-compound and compound-compound connections***

We use Pearson correlation on aligned profiles of a gene and compound to score their connection. The profiles are obtained by averaging the replicate profiles feature-wise. We empirically found that for gene-compound connections, an absolute score value greater than 0.35 indicates similar/opposite phenotypes in the two profiles and used this for validation experiments. For the follow-up experiments of a gene, we typically used a more stringent filter of 0.40 and picked a balanced number of strongest-correlating compounds and strongest anti-correlating, subject to availability in the chemical library.

To validate how many genes find a compound with relevant targets among their top matches, we obtain the rank of the first relevant compound match for each gene, and simply count the number of genes for which the rank is within the top 306 compounds (1% of compounds, out of 30,616).

The 69 genes used as queries in the experiment were taken from the 110 reagents scored as expressing a Cell Painting phenotype in (Rohban et al. 2017) by excluding engineered variants of genes, and by selecting a single reference / wild-type construct when multiple were available (e.g. isoforms).

***Permutation test for validation of known compound-compound pairs***

The odds ratio of how likely compound pairs that are strongly correlated have the same MoA is considered as the test statistic. The compound names are then shuffled and the odds ratio is re-calculated. This yields a sample from the null hypothesis. The re-calculation is repeated n times. The actual odds ratio, without any shuffling of compound names, is ranked against the null distribution samples to produce a p-value estimation.

***Permutation test for validation of number of genes with a relevant compound match***

We run a permutation test to test the significance of the number of genes (20 out of 63) that have a match in the top 1% of their rank ordered list of compounds (with the rank based on their correlation to the gene). To generate a sample from the null distribution, the compounds are shuffled randomly, and the number of such genes is calculated. The p-value is calculated based on the empirical sampled null distribution. Note that (1) we perform a single test, where the test statistic is the number of genes, and therefore no correction for multiple testing is required here, and (2) that this procedure automatically accounts for the fact that some genes have many associated compounds (many matches) in the set.

***Enrichment analysis plots***

We follow the same logic as the Gene Set Enrichment Analysis (GSEA) [6]. The set of gene-compound pairs are sorted based on their profile correlations on the x-axis. On the y-axis, the plot goes up by a certain fixed amount if the corresponding gene-compound is a valid pair. Otherwise, the plot goes down by the same fixed amount. Scanning the x-axis from left to right, early existence of abundant valid pairs results in a rapid jump of the plot, and illustrates higher enrichment of profile correlation for being indicative of biological relevance between the gene-compound pair.

***Relationship between strength of gene-compound matching and the number of correct gene annotations***

For the analysis shown in Supplementary Figure S5, we calculated the spearman correlation between the absolute correlation of correct gene-compound pairs (including BioGrid-extended genes) and the number of direct protein targets of the compound (including only the direct annotated target genes of each compound, to better capture true polypharmacology).

***Chemical structure similarity (calculating Tanimoto coefficients for hit compounds)***

Similarity matrices were calculated using the Pipeline Pilot (Biovia, version 18.1.0) "Molecular Similarity NxN" function with ECFP4 fingerprints (Rogers and Hahn 2010) and Tanimoto similarity coefficients (Bajusz et al. 2015). Heatmaps were generated using the Seaborn library (0.11.2) for Python.

***Assessing compound purity and identity***

Compound purity and identity were determined by ultra performance liquid chromatography coupled with mass spectroscopy (UPLC-MS). Purity was measured by UV absorbance at 210 nm. Identity was determined on a SQ mass spectrometer by positive and/or negative electrospray ionization. Mobile phase A consisted of either 0.1% ammonium hydroxide or 0.05% trifluoroacetic acid in water, while mobile phase B consisted of either 0.1% ammonium hydroxide or 0.06% trifluoroacetic acid in acetonitrile. The gradient ran from 5% to 95% mobile phase B over 2.65 min at 0.9 mL/min. An Acquity BEH C18, 1.7 µm, 2.1x50 mm column was used with column temperature maintained at 65 degrees C. Compounds were dissolved in DMSO at a nominal concentration of 1 mM, and 1.0 µL of this solution was injected.

***SMAD3 experiments***

For SMAD3 compounds, given a limit of 10 compounds to study, we chose the top five positive matches and the top two negative matches (which were somewhat cytotoxic based on cell count in the Cell Painting assay), along with three additional negative matches (among the top 15) which were less cytotoxic. One was unavailable.

A549 lung carcinoma cells were transfected with the luciferase reporter plasmids 4xSBE-Luc to measure TGF-b/Smad3-activated transcription [7] and pRL-TK (low expressing, constitutively active Renilla luciferase under the HSV-thymidine kinase promoter) (Promega cat# E224A) to normalize for the 4xSBE Firefly luciferase values. The transfected cell lysates were processed for luciferase assays as described[56] and per manufacturer's protocol (Promega). In brief, the cells were seeded in 24-well plates at 80% confluency and, after adhering, the media was changed to growth or starvation media (RPMI-1640 with 10% or 2% FBS respectively) for 6 hours. The cells were then transfected with 200 ng 4xSBE-Luc and 100ng Rl-Tk-Luc reporter plasmids per well using Lipofectamine 2000 per manufacturer recommendations (Thermo Fisher cat# 11668019). 12 hours after transfection cells were treated for 24 hours with 5 ng/ml TGF-β1 or 5 µM SB431542 to inhibit TGF-b-induced Smad activation, and either of 9 compounds at 10µM in triplicate. All cells were harvested with 200 µl of passive lysis buffer (Promega). Luciferase assays were performed using a Dual-Luciferase assay kit (Promega), and luciferase activities were quantified with a SpectraMax M5 plate luminometer (Molecular Devices) and normalized to the internal Renilla luciferase control and DMSO control.

***Ras experiments***

Isogenic RAS-less mouse embryonic fibroblast cell lines driven by human KRAS4b G12D, HRAS WT, or BRAF V600E alleles were plated in 384-well plates and dosed with compound or DMSO 18 hours later using an Echo acoustic liquid handler in a 10 point, 2-fold dilution in 0.2% DMSO, with 10µM as the top concentration. After

72 hours, Promega CellTiter-Glo® reagent was added, and the signal was read using Envision software. Values were normalized using day zero and DMSO control readings. Hits were determined by a one log difference in IC50 values between BRAF V600E and RAS-driven cell line responses.

***Casein-kinase 1 alpha experiments***

CSNK1A1 enzymatic assays were performed by mobility shift assay using the Labchip EZ Reader II (Perkin Elmer). GST-tagged human CSNK1A1 (Carna Biosciences) protein was incubated with ATP, substrate, and assay buffer (20 mM Hepes - pH 7.5, 5 mM $MgCl_2$, and 0.01% Triton X-100). The assay reaction was initiated with 5 μM ATP, 2 mM DTT, and 1 μM Profiler Pro FL-Peptide 16 substrate (Perkin Elmer). Curve fitting and determination of AC50 values for phosphorylation inhibition were performed using Genedata.

***GSK3B experiments***

The compounds with a Cell Painting profile matching or opposing GSK3 overexpression were tested against GSK3α and GSK3β as previously reported [8]. Purified GSK3β or GSK3α was incubated with tested compounds in the presence of 4.3 μM of ATP (at or just below Km to study competitive inhibitors) and 1.5 μM peptide substrate (Peptide 15, Caliper) for 60 minutes at room temperature in 384-well plates (Seahorse Bioscience) in assay buffer that contained 100 mM HEPES (pH 7.5), 10 mM $MgCl_2$, 2.5 mM DTT, 0.004% Tween-20, and 0.003% Brij-35. Reactions were terminated with the addition of 10 mM ethylenediaminetetraacetic acid (EDTA). Substrate and product were separated electrophoretically, and fluorescence intensity of the substrate and product was determined by Labchip EZ Reader II (Perkin Elmer). The kinase activity was measured as percent conversion to product. The reactions were performed in duplicate for each sample. The positive control, CHIR99021, was included in each plate and used to scale the data in conjunction with “in-plate” DMSO controls. The results were analyzed by Genedata Assay Analyzer. The percent inhibition was plotted against the compound concentration, and the IC50 value was determined from the logistic dose-response curve fitting. Values are the average of at least three experiments. Compounds were tested using a 12-point dose curve with 3-fold serial dilution starting from 33 μM. The two most active compounds were resynthesized for validation and tested along with closely related analogs (Supplementary Methods).

***p38 experiments***

Cell Painting profiles for two wild-type variants of p38α (MAPK14) were averaged to create a p38α Cell Painting profile. 20 compounds whose Cell Painting profile correlated positively or negatively to that of p38α overexpression were selected; we also chose 14 "non-correlated" compounds (i.e. absolute value of correlation <0.2) as negative/neutral controls. The compounds were tested for their influence on p38 activity using the RPE1-p38 kinase translocation reporter (KTR) line that was previously generated[9]. This cell line has been tested and confirmed to be negative for mycoplasma contamination, but not authenticated. p38 activity is measured by phosphorylation of its substrate, MEF2C, which is preferentially phosphorylated by p38α, while p38β and p38δ contribute less [10]. RPE1-p38KTR cells were cultured in DMEM/F12 medium supplemented with 10% Fetal Bovine Serum at 37C in a humidified atmosphere with 5% $CO_2$. 1000 cells were plated per well in 96-well plates and treated with 1μM and 10μM of each compound (n=4 well per concentration per compound, no replicates) for 48 hours. Only the middle 60 wells were used to prevent potential confounds from the edge effect. Cells were then fixed in 4% paraformaldehyde for 10min, followed by permeabilization in cold methanol at -20C for 5min. Cells were stained with 0.4 μg/mL Alexa Fluor 647 carboxylic acid, succinimidyl ester for 2hr at RT, followed by 1μg/mL DAPI for 10min at RT to facilitate the segmentation of individual cells.

p38 activity in single cells was calculated using the ratio of the median intensity of the p38-KTR in a 5-pixel-wide cytoplasmic ring around the nucleus to the median intensity of the p38-KTR in the nucleus. p38 activity measurements were normalized to DMSO within the same plate and column. The Student's t-test or Kolomogorov-Smirnov (KS) test was used to assess the significance of changes in the single cell distributions of p38 activity for each compound relative to control; we note that even for the positive control known inhibitor the effect sizes are small. When reporting hits from the assay, KS test and t-test p-values were adjusted to control the false discovery rate using the Benjamini-Hochberg method, using the p.adjust(method='BH") method in R.

***PPARGC1A (PGC-1α) experiments***

*Reporter assays:* To measure PGC-1α activity related to PPARG, RT112/84 cells were obtained from the Cancer Cell Line Encyclopedia (Broad Institute, Cambridge, MA), which obtained them from the original source and performed cell line authentication. The cell line was engineered with the NanoLuc gene cloned into the 3' UTR of the FABP4 (previously described[11]) followed by stable expression of nuclear GFP (pTagGFP2-H2B, Evrogen) and tested negative for mycoplasma (MycoAlert, Lonza). Cells were plated in 384-well plates at ~10,000 cells/well and dosed with indicated compounds in the absence or presence of EC50 of PPARG agonist, rosiglitazone, using an HP D300 digital dispenser. The following day nuclei were counted for normalization (IncuCyte S3, Essen Bioscience) and the reporter activity was evaluated using the NanoGlo Luciferase Assay System (Promega). Normalized data is reported as NanoGlo arbitrary light units divided by cell number. PPARG agonist, rosiglitazone, and inhibitor, T0070907, were obtained from Tocris and included as controls.

To measure effects on PGC1a/ERRalpha, HEK293T cells purchased from ATCC were co-transfected with Gal4-ERRalpha, with and without PGC1a (pCDNA3.1-Flag-HA-PGC-1alpha [12]), kind gifts from Pere Puigserver, in combination with the Gal4 UAS reporter construct, pGL4.35 [luc2P/9XGAL4UAS/Hygro] (Promega) modified by subcloning the HSV-TK promoter into the unique HindIII site that is downstream of the 9xGal4 UAS sites, in addition to a Renilla luciferase expression vector pRL (Promega) for normalization. Cells were dosed with compounds and 24 hours later, plates were analyzed using Dual-Glo Luciferase Assay System (Promega). Normalized light units are reported as Firefly luciferase divided by Renilla luciferase. ERRalpha modulators XCT790, Daidzein, and Biochanin A (Cayman Chemical) were included as controls. 293T cells were not authenticated nor tested for mycoplasma.

*High content mitochondrial motility screen:* We used our previously published assay to assess mitochondrial motility[13]. Briefly, we plated E18 rat cortical neurons in the middle 60 wells of 96 well plates (Greiner) – 40,000 cells per well in 150 µl enriched Neurobasal media. Neurons were transfected with mito-DsRed at DIV7 using Lipofectamine2000 (Life Technologies). Plating and transfection were all done using an Integra VIAFLO 96/384 automated liquid handler. At DIV9, test compounds were added into wells to achieve a final concentration of 10 µM each (4 wells per compound), as well at 10 µM calcimycin for neg. control, and DMSO only for mock treatment. Following a 1-2 hour incubation, plates were imaged on a ArrayScan XTI (Thermo Fisher). Mitochondrial motility data was extrapolated from imaging data using a MATLAB and CellProfiler based computational pipeline. Compounds A01-A12 were tested on one plate; B01-B11 were tested separately on another plate on the same day. The experiment was repeated twice in different weeks. In the second week, TMRE was added to all wells after imaging was completed (20min, then 2 washes) and imaged to measure mitochondrial membrane potential in order to determine mitochondrial and cell health.

***YAP1-related compounds***

For the initial experiments, quality control of the compounds revealed that purity was 88% for A15 (BRD-K34692511-001-01-9), 81% for A05 (BRD-K28862419-001-01-9), and > 99% for E07 (BRD-K43796186-001-01-1). For subsequent experiments in the Eisinger lab, BRD-K43796186 (NB4A) was ordered from MuseChem (cat. #M189943) and for the Kiessling lab, from Ambinter (Cat # Amb2554311).

##### *YAP1 cell culture and treatments*

Eisinger lab: Murine KP230 cells, a Yap1-dependent cancer cell line, were derived from a tumor from the KP mouse model (*Kras*$^{G12/D}$; *Trp53*$^{fl/fl}$*),* as described in[14]. STS-109 UPS cells were derived from a human UPS tumor and validated by Rebecca Gladdy, MD (Sinai Health System, Toronto, Ontario, Canada). TC32 cells were a gift from Patrick Grohar, MD, PhD (Children's Hospital of Philadelphia). HT-1080, HCT-116, and HEK293T cells were purchased from ATCC. KP230, HT-1080, and HEK-293T cells were grown in DMEM with 10% FBS, 1% L-glutamine, and 1% penicillin/streptomycin (P/S). STS-109 cells were cultured in DMEM with 20% FBS, 1% L-glutamine, and 1% P/S. TC32 cells were grown in RPMI with 10% FBS, 1% L-glutamine, and 1% P/S. HCT-116 cells were cultured in McCoy's 5A medium with 10% FBS and 1% P/S. All cells were confirmed to be negative for mycoplasma contamination and were maintained in an incubator at 37C with 5% $CO_2$. For experimental purposes, cells were cultured for up to 20 passages before being discarded, and were grown to approximately 50% confluence to circumvent the effects of high cell density on Yap1 expression and activity. All cell lines in the Eisinger laboratory were treated with 10 μM of each inhibitor or an equivalent volume of DMSO every 24 hours for 3 days, except for STS-109 cells, which were treated daily for 8 days.

Kiessling lab: H9 hPSCs (WiCell) were maintained on vitronectin (Thermo Fisher)-coated plates in Essential 8 (E8) medium. The cells were routinely passaged using 0.5mM EDTA and treated with 5μM Y-27632 dihydrochloride (Tocris) on the first day. For testing the effects of the small molecules, H9 hPSCs were seeded at 50K cells/cm$^2$ on vitronectin-coated plates in E8 medium supplemented with 5μM Y-27632 dihydrochloride (day 0). On day 1, the medium was switched to E8 medium. On day 2, the medium was switched to E8 medium supplemented with the small molecules. Following overnight incubation, the cells were collected for subsequent analysis on day 3. The cells were regularly checked for Mycoplasma contaminations (Sigma Aldrich - Lookout Mycoplasma PCR Detection Kit) but were not authenticated.

Boerckel lab: Murine periosteal cells were isolated from a transgenic mouse model (CMV-Cre;R26R-rtTA$^{fl}$; tetO-YAP$^{S127A}$; C57Bl/6 strain/background) in which YAP1 can be activated in a doxycycline inducible manner (Camargo 2011). This mouse model expresses a mutant form of YAP1 (YAP$^{S127A}$) that escapes degradation. Cells were isolated from 3 female mice (age 15 weeks) from a 4-day old femoral fracture callus. Cells were cultured in a-MEM with 15% Fetal Bovine serum (S11550, R&D Systems), 1% GlutaMAX-I (Gibco, 35050-061) and 1% Penicillin/Streptomycin (Gibco, 15140-122).

##### *YAP1-related lentiviral production*

Knockdown of *YAP1* in HCT-116 cells was performed with shRNAs (TRC clone IDs: TRCN0000107266 and TRCN0000107267); a scrambled shRNA was used as a negative control. shRNA plasmids (Dharmacon) were packaged using the third-generation lentiviral vector system (pVSV-G, pMDLG, and pRSV-REV; Addgene) and expressed in HEK-293T cells using Fugene 6 transfection reagent (Promega). Virus-containing supernatants were collected 24 and 48 hours after transfection and concentrated 40-fold by centrifugation with polyethylene glycol 8000.

##### *YAP1-related proliferation assays*

NB4A treatment: Cells were treated with 10 µM of each inhibitor or an equivalent volume of DMSO every 24 hours for 3-8 days, and counted with a hemocytometer with trypan blue exclusion daily (KP230, HT-1080, TC32, HCT-116), or every 2 days (STS-109).

BRD-K28862419 and BRD-K34692511 analog screen: KP230 cells were treated with 10 µM of each compound or an equivalent volume of DMSO every 24 hours for 2 days. Treated cells were then incubated with PrestoBlue viability reagent (ThermoFisher Scientific) for 2 hours according to the manufacturer's recommendations. Fluorescence (560/590 nm) was read on a Spectramax M2e plate reader (Molecular Devices).

shRNA-mediated *YAP1* knockdown: HCT-116 cells were infected with *YAP1* shRNA-encoding lentiviruses in the presence of 8 µg/mL polybrene (Sigma). Antibiotic selection (3 µg/mL puromycin) was performed after 48 hours, after which cells were cultured for an additional 48 hours. Cells were then trypsinized, seeded under puromycin-selection conditions, and counted with a hemocytometer with trypan blue exclusion on days 7, 8, and 9 post-infection.

***YAP1-related qRT-PCR***

For the Eisinger lab, total RNA from cultured cells was isolated with the QIAGEN RNeasy mini kit, and cDNA was synthesized with the High-Capacity RNA-to-cDNA kit (Life Technologies). qRT-PCR analysis was performed with TaqMan "best coverage" probes on a ViiA7 instrument. Hypoxanthine phosphoribosyltransferase (*HPRT*) and succinate dehydrogenase subunit A ( *SDHA*) were used as endogenous controls. Relative expression was calculated using the ddCt method.

For the Kiessling lab, the RNA was extracted using TRIzol (Life Technologies) and Direct-zol™ RNA MiniPrep kit (Zymo Research) as per manufacturer instructions. The RNA was reverse transcribed using iScript cDNA synthesis kit (Bio-Rad). The qPCR was performed on CFX Connect (Bio-Rad) using iTaq Universal SYBR Green Supermix (Bio-Rad). GAPDH was used as a reference gene for normalization. The relative gene expression levels were determined using the ddCt method. The primer sequences used are listed in Supplementary Table S7.

For the Boerckel lab, to induce $YAP^{S127A}$, 1µM doxycycline was added to the cell culture medium for 48 hours. This was used as a positive control to compare YAP1 mRNA expression. Cells were also treated with BRD-K34692511-001-01-9 at 5µM. mRNA was isolated from cells (n=3/group/time point) at 1, 4 or 48 hours after treatment using Qiagen RNeasy Mini kit (Qiagen, 74106). cDNA was prepared as per the manufacturer's protocol using the High-Capacity Reverse Transcription kit (Thermofisher scientific, 4368814). qPCR analysis was performed using the QuantStudio 6 Pro Real-Time PCR System.

***YAP1-related reporter assay***

Varelas lab: HEK293T cells purchased from ATCC were co-transfected using Lipofectamine 3000 (Thermo Fisher) with a TEAD luciferase reporter construct, 8xGTIIC-luciferase (gift from Stefano Piccolo, Addgene plasmid # 34615), a plasmid expressing Renilla Luciferase from a CMV promoter as a transfection control, along with a plasmid expressing 3xFlag-tagged wild-type YAP1 from a CMV promoter (pCMV5 backbone). Following transfection the cells were immediately treated with 0.2% DMSO, 10µM NB4A, BRD-K34692511 or BRD-K28862419 and then lysed 48 hours later. Lysates were examined using the Dual-Luciferase Reporter Assay System (Promega) according to the manufacturer's protocol and measured using a SpectraMax iD3 plate reader (Molecular Devices). Firefly Luciferase activity from the TEAD reporter was normalized to Renilla

Luciferase activity and then plotted as relative values. Mycoplasma tests are routinely performed, but cells were not recently authenticated.

***YAP1-related RNA-sequencing and data analysis***

Total RNA from cultured cells was isolated with the QIAGEN RNeasy Mini Kit with on-column DNase digestion. RNA quality checks were performed with an Agilent 2100 Bioanalyzer (Eukaryotic Total RNA Nano kit). Library preparation (500 ng input RNA) was performed with the NEBNext Poly(A) mRNA Magnetic Isolation Module (#E7490) with SPRIselect Beads (Beckman Coulter), the NEBNext Ultra II Single-End RNA Library Prep kit (#7775S), and the NEBNext Multiplex Oligos for Illumina (Index Primers Set 1) according to the manufacturer's instructions. Library size was confirmed with an Agilent 2100 Bioanalyzer (DNA1000 chip). Pooled libraries were diluted to 1.8 pM (concentrations checked with the Qubit Fluorometer high-sensitivity assay, Thermo Fisher), and sequenced on an Illumina NexSeq 500 instrument with the NexSeq 500 75-cycle high-output kit.

For data analysis, FASTQ files were generated with the *bcl2fastq* command line program (Illumina). Transcript alignment was performed with Salmon [15]. Differential expression analysis (NB4A- vs. DMSO-treated cells) was performed with the DESeq2 R package. DESeq2 "stat" values for each gene were used as inputs to pre-ranked GSEA, where enrichment was tested against the Hallmark gene sets from the Molecular Signatures Database (MSigDB). Access to sequencing data is discussed in the data availability section.

***YAP1-related Western blotting***

For the Kiessling lab, the cells were lysed in RIPA buffer (Pierce) supplemented with Halt Protease inhibitor cocktail and Halt Phosphatase inhibitor cocktail (Thermo Fisher). The Eisinger lab lysed cells in hot Tris-SDS buffer (pH 7.6) and boiled for 5 minutes at 95ºC. The protein concentration of each sample was quantified using the Pierce BCA protein assay (Thermo Fisher). The proteins were resolved by SDS-PAGE and transferred to PVDF membranes using the Trans-Blot Turbo Transfer system (Bio-Rad). The membranes were blocked in 5% non-fat milk in TBS-T for up to 1 hour at room temperature and incubated with primary antibodies in 5% bovine serum albumin in TBS-T overnight at 4ºC. Then, the membranes were incubated with HRP-conjugated anti-rabbit IgG secondary antibodies at 1:10000 (Kiessling lab; Jackson ImmunoResearch Laboratories, #111-035-003) or 1:2500 (Eisinger lab; Cell Signaling Technology [CST] #7074) for 1 hour at RT and developed in the ChemiDoc MP Imaging system (Kiessling lab) or on autoradiography film (Eisinger lab) using ECL Prime reagent (Amersham). The band intensities in immunoblots were quantified with Image Lab software. The primary antibodies and dilutions used are: anti-YAP1 (CST 4912S and CST 14074 [clone D8H1X]) at 1:1000, anti-phospho-YAP1-S127 (CST 4911S) at 1:1000, and anti-GAPDH (CST 5174 and CST 2118 [clone 14C10]) at 1:15000 and 1:1000, respectively. Primary antibodies were validated commercially in cells both wild-type and deficient (e.g., knockout) for the gene/protein of interest. YAP1-related immunofluorescence and image analysis

For the Eisinger lab, cells grown on poly-L-lysine-coated chamber slides were fixed in 4% PFA (15 minutes at room temperature), permeabilized with 0.5% Triton-X100/PBS (15 minutes at room temperature), and blocked with 5% goat serum (Vector Laboratories S-1000; 1 hour at room temperature). Cells were then incubated with anti-Yap1 primary antibodies (CST #14074 [clone D8H1X]; 1:1000) diluted in blocking buffer overnight at 4ºC. Subsequently, cells were incubated with Alexa Fluor 488-conjugated secondary antibodies (4 ug/mL in blocking buffer; Thermo Fisher Scientific #A-11008) for 1 hour at room temperature. Coverslip mounting was performed with ProLong Gold Antifade reagent with DAPI. Images (5 fields per condition for each of 3 independent experiments) were acquired with a Nikon Eclipse Ni microscope and Nikon NES Elements software. Image analysis was performed with Fiji as follows: For nuclear staining intensity, watershed analysis of DAPI channel images (8-bit) was performed to “separate” nuclei that appeared to be touching. Nuclei were then converted to regions of interest (ROIs) that were “applied” to the corresponding GFP channel image (8-bit format). Analysis of staining intensity in these nuclear ROIs was then performed, excluding objects smaller than 100 pixels$^2$ (integrated density normalized to number of nuclei). A similar process was followed to determine whole-cell staining intensity: using 8-bit GFP channel images, cells were distinguished from background via thresholding, and converted to ROIs that were applied back to the 8-bit GFP channel images. Analysis of staining intensity (integrated density normalized to number of nuclei) was then performed in these ROIs, excluding objects smaller than 500 pixels$^2$. The ratio of nuclear to total Yap1 expression was determined after subtracting out background GFP signal from no-primary antibody controls.

For the Kiessling lab, the cells were fixed with 4% formaldehyde for 15 mins at room temperature. The cells were permeabilized and blocked with PBS containing 2% BSA and 0.1% Triton-X100. The cells were incubated with a primary antibody against YAP1 (Santa Cruz Biotechnology, sc-101199) at 1:200 dilution in a blocking buffer overnight at 4°C. Then, the cells were incubated with a goat anti-mouse Alexa Fluor 488 conjugated secondary antibody (Thermo Fisher, #A11001) at 1:1000 dilution for 1 hour at room temperature. The nuclei were counterstained with DAPI dilactate (Molecular Probes). Images were collected with Olympus FV1200 microscope and analyzed with CellProfiler. Briefly, nuclei and cell bodies were segmented using DAPI and YAP channels respectively. The cell cytoplasm was determined as the region outside nuclei but within the cell bodies. Then, the ratio of mean intensity of YAP in the nucleus to cytoplasm was calculated to determine YAP translocation.

***Chemical synthesis of BRD4486***

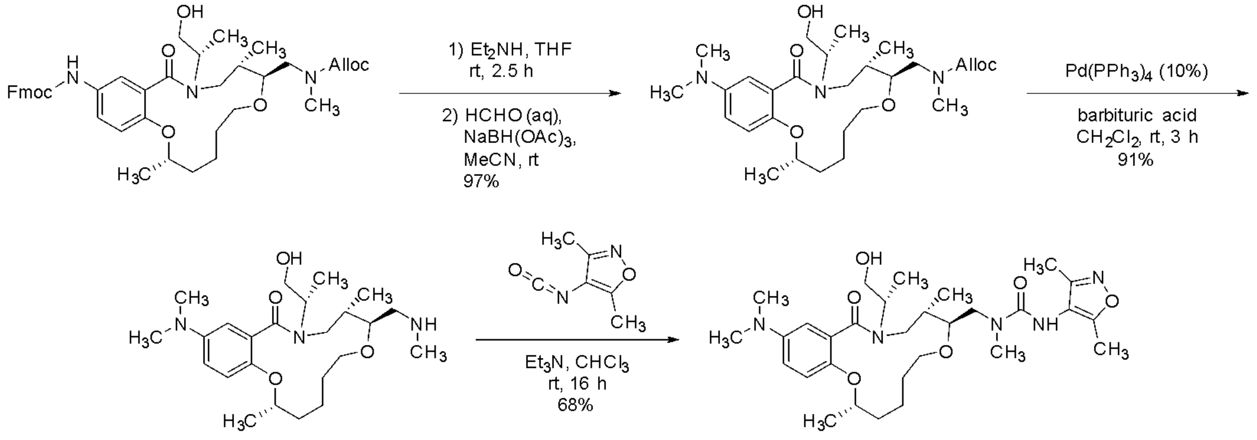


BRD4486

We followed prior work [16]. To a solution of the Fmoc-protected intermediate (100 mg, 0.140 mmol, 1 eq) in tetrahydrofuran (1 mL) was added diethylamine (150 µL, 1.44 mmol, 10.3 eq). The mixture was stirred 2.5 h, and the solvent and excess amine were evaporated.

The residue was dissolved in acetonitrile (1 mL), then formaldehyde (37% aqueous, 100 µL, 1.32 mmol, 9.4 eq), catalytic acetic acid (1 drop), and sodium triacetoxyborohydride (100 mg, 0.472 mmol, 3.4 eq) were added. The mixture was stirred 1 h, then quenched with saturated aqueous $NaHCO_3$ and diluted with $CH_2Cl_2$. The layers were separated, and the aqueous layer was extracted with $CH_2Cl_2$. The combined organic layers were dried over $MgSO_4$, filtered, and evaporated, and the residue was purified by flash column chromatography (0-10% MeOH/$CH_2Cl_2$) to provide the dimethylaniline (70.2 mg, 97% yield).

A yellow solution of the resulting Alloc-protected amine (68 mg, 0.131 mmol, 1 eq), 1,3-dimethyl-barbituric acid (200 mg, 1.28 mmol, 9.8 eq), and tetrakis(triphenylphosphine)palladium(0) (15 mg, 0.013 mmol, 0.1 eq) in dichloromethane (1 mL) was stirred at room temperature for 3 h. The reaction was quenched with 0.1 M HCl and stirred 15 min. The layers were separated, and the organic layer was extracted with 0.1 M HCl. The pH of the combined aqueous layers was adjusted to ~12 with 4 M NaOH, then extracted with $CH_2Cl_2$. The combined organic layers were dried over $Na_2CO_3$, filtered, and evaporated to provide the crude free amine (52 mg, 91% yield), which was used directly in the urea formation.

To a solution of the crude secondary amine (52 mg, 0.119 mmol, 1 eq) and 3,5-dimethylisoxazol-4-yl isocyanate (20 mg, 0.145 mmol, 1.2 eq) in chloroform (1 mL) was added triethylamine (16.5 µL, 0.119 mmol, 1 eq). The mixture was stirred at room temperature for 16 h, and the solvent was evaporated. The residue was purified by flash column chromatography (0-10% MeOH/$CH_2Cl_2$) to provide **BRD4486** (46.2 mg, 68% yield) as a white solid.

MS: 574.5 $[M+H]^+$; 572.3 $[M-H]^-$

#### *Chemical synthesis of BRD4313*

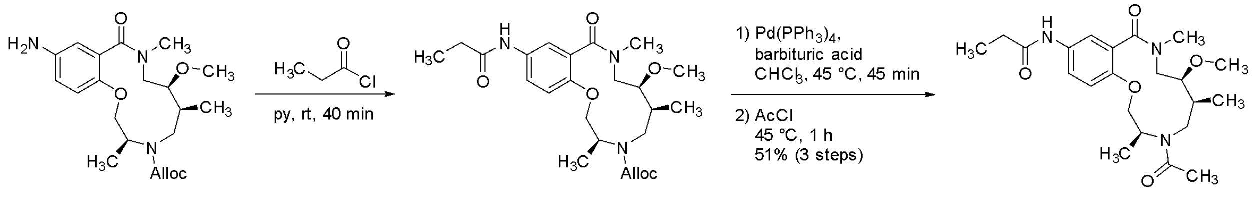


We followed prior work [17]. To the aniline intermediate (100 mg, 0.247 mmol, 1 eq) dissolved in pyridine (0.5 mL) was added propionyl chloride (45 µL, 0.514 mmol, 2.1 eq). The resulting mixture was stirred at room temperature for 40 min.

Chloroform (0.5 mL) and 1,3-dimethylbarbituric acid (385 mg, 2.46 mmol, 10 eq) were added, and, after 5 min, tetrakis(triphenylphenylphosphine)palladium(0) (28 mg, 0.024 mmol, 0.1 eq) was added. The yellow-green solution was stirred at 45 °C for 45 min ($CO_2$ evolution), then acetyl chloride (261 µL, 3.69 mmol, 15 eq) was added. After stirring 1 h, the mixture was diluted with $CH_2Cl_2$ and poured into 0.1 M HCl. The layers were separated, and the aqueous layer was extracted with $CH_2Cl_2$. The combined organic layers were washed with saturated aqueous $NaHCO_3$, dried over $MgSO_4$, filtered, and evaporated. The residue was purified by flash column chromatograph (2.5-10% MeOH/$CH_2Cl_2$) to provide the product (82.4 mg) in 84% purity. The material was further purified by mass-directed prep-HPLC (0-100% MeCN/water + 0.1% TFA) to provide **BRD4313** (54.6 mg, 51% yield over 3 steps) as a tan solid.

MS: 420.8 [M+H]+

#### *Analog selection for GSK experiments*

Compounds for (stereochemical) structure-activity relationship determination were selected from the Broad compound library in the following order until 24 analogs had been reached for each hit:

1) All available stereoisomers of the hit compound
2) Available regioisomers of the hit compound
3) Analogs with high structural similarity (Tanimoto ≥ 0.75) to the hit compound, selected for chemical diversity

Assay plates for each series (RCM/H2T) were prepared with 4 on-plate controls plus the hit compound from the opposing series. The controls included 3 dual GSK3α/β inhibitors (CHIR-99021, GW8510, and BRD0320) and one GSK3α-specific inhibitor (BRD0705). All compounds tested are shown below.

#### **BRD4486** (RCM) Series Assay Plate

| Core ID | Structure | Note | Core ID | Structure | Note | Core ID | Structure | Note |
|---|---|---|---|---|---|---|---|---|
| BRD-K00760705 | 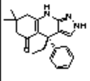 | control | BRD-K94445007 | 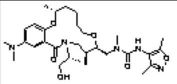 | RCM stereoisomer | BRD-K07274638 | 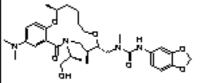 | RCM analog |
| **CHIR99021** BRD-K16189898 | 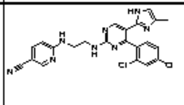 | control | BRD-K23628938 | 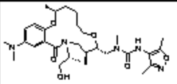 | RCM stereoisomer | BRD-K18142082 | 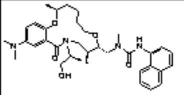 | RCM analog |
| **GW8510** BRD-K55000304 | 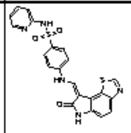 | control | BRD-K36176998 | 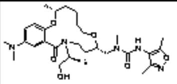 | RCM stereoisomer | BRD-K07891707 | 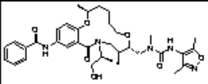 | RCM analog |
| BRD-K87550320 | 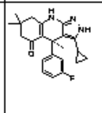 | control | BRD-K94445007 | 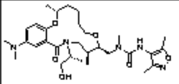 | RCM stereoisomer | BRD-K26973634 | 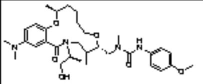 | RCM analog |
| BRD-K26994486 | 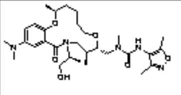 | RCM hit | BRD-K10781609 | 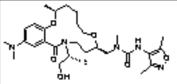 | RCM stereoisomer | BRD-K57195030 | 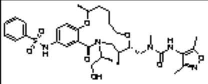 | RCM analog |
| BRD-K91354313 | 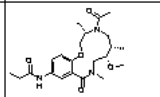 | H2T hit | BRD-K34254821 | 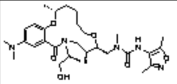 | RCM stereoisomer | BRD-K71568523 | 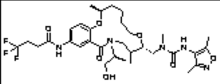 | RCM analog |
| BRD-K97242998 | 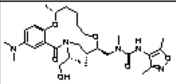 | RCM stereoisomer | BRD-K65054381 | 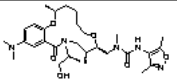 | RCM stereoisomer | BRD-K72681403 | 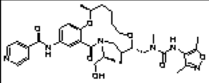 | RCM analog |
| BRD-K56373119 | 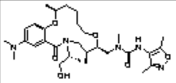 | RCM stereoisomer | BRD-K09376136 | 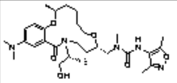 | RCM stereoisomer | BRD-K61284358 | 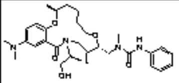 | RCM analog |
| BRD-K86938753 | 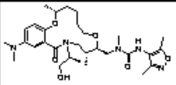 | RCM stereoisomer | BRD-K12710356 | 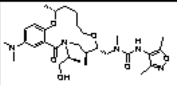 | RCM stereoisomer | BRD-K18599462 | 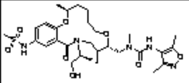 | RCM analog |
| BRD-K59522787 | 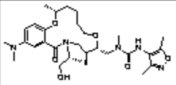 | RCM stereoisomer | BRD-K96280720 | 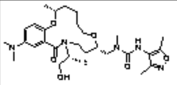 | RCM stereoisomer | BRD-K55276005 | 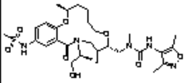 | RCM analog |

#### **BRD4313** (H2T) Series Assay Plate

| Core ID | Structure | Note | Core ID | Structure | Note | Core ID | Structure | Note |
|---|---|---|---|---|---|---|---|---|
| BRD-K00760705 | 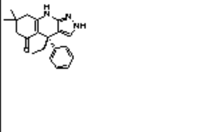 | control | BRD-K74022101 | 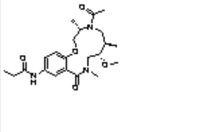 | H2T stereoisomer | BRD-K74198088 | 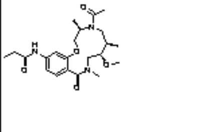 | H2T regio/stereoisomer |
| **CHIR99021** BRD-K16189898 | 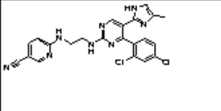 | control | BRD-K83538063 | 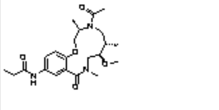 | H2T stereoisomer | BRD-K06631334 | 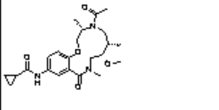 | H2T analog |
| **GW8510** BRD-K55000304 | 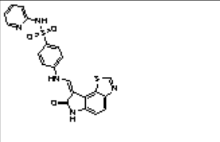 | control | BRD-K97378175 | 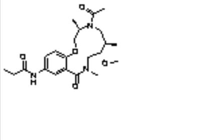 | H2T stereoisomer | BRD-K28720546 | 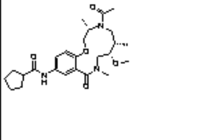 | H2T analog |
| BRD-K87550320 | 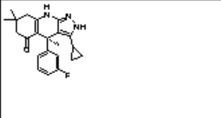 | control | BRD-K00411149 | 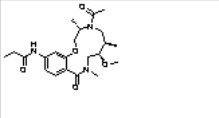 | H2T regio/stereoisomer | BRD-K31377965 | 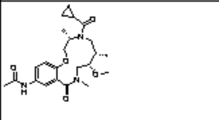 | H2T analog |
| BRD-K91354313 | 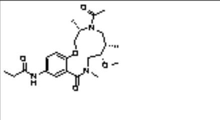 | H2T hit | BRD-K04403306 | 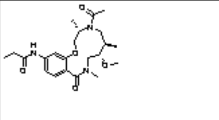 | H2T regio/stereoisomer | BRD-K33863933 | 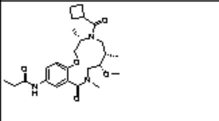 | H2T analog |
| BRD-K26994486 | 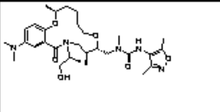 | RCM hit | BRD-K11844860 | 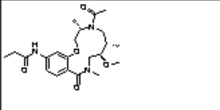 | H2T regio/stereoisomer | BRD-K44217085 | 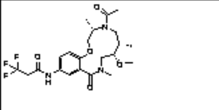 | H2T analog |
| BRD-K12299093 | 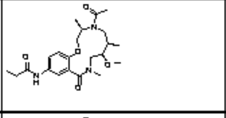 | H2T stereoisomer | BRD-K20077349 | 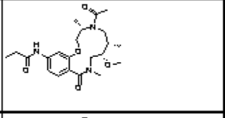 | H2T regio/stereoisomer | BRD-K48114548 | 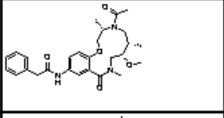 | H2T analog |
| BRD-K42803695 | 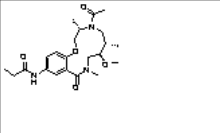 | H2T stereoisomer | BRD-K34274486 | 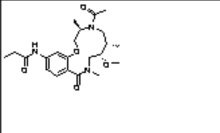 | H2T regio/stereoisomer | BRD-K58769468 | 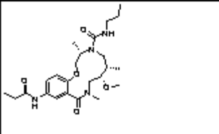 | H2T analog |
| BRD-K53538949 | 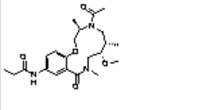 | H2T stereoisomer | BRD-K54036439 | 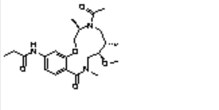 | H2T regio/stereoisomer | BRD-K76966915 | 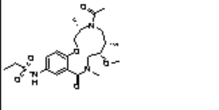 | H2T analog |
| BRD-K61699306 | 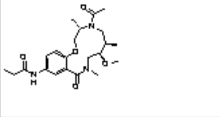 | H2T stereoisomer | BRD-K59865704 | 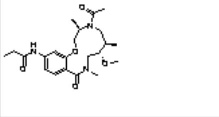 | H2T regio/stereoisomer | BRD-K80986480 | 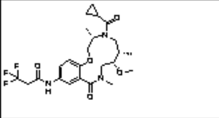 | H2T analog |

##### ***Mobility shift microfluidics assay protocol***

We followed previous protocol [8]. Purified GSK3β or GSK3α was incubated with tested compounds in the presence of 4.3 µM of ATP (at or just below Km to study competitive inhibitors) and 1.5 µM peptide substrate (Peptide 15, Caliper) for 60 minutes at room temperature in 384-well plates (Seahorse Bioscience) in assay buffer that contained 100 mM HEPES (pH 7.5), 10 mM $MgCl_2$, 2.5 mM DTT, 0.004% Tween-20, and 0.003% Briji-35. Reactions were terminated by the addition of 10 mM ethylenediaminetetraacetic acid (EDTA). Substrate and product were separated electrophoretically, and fluorescence intensity of the substrate and product was determined by Labchip EZ Reader II (Caliper Life Sciences). The kinase activity was measured as percent conversion. The reactions were performed in duplicate for each sample. The positive control, CHIR99021, was included in each plate and used to scale the data in conjunction with in-plate DMSO controls. The results were analyzed by Genedata Assay Analyzer. The percent inhibition was plotted against the compound concentration, and the $IC_{50}$ value was determined from the logistic dose-response curve fitting. Values are the average of at least three experiments. Compounds were tested using a 12-point dose curve with 3-fold serial dilution starting from 33 µM.

***Assay Results***

**RCM**

| Core ID | GSK3α $IC_{50}$ μM | GSK3β $IC_{50}$ μM |
|---|---|---|
| BRD-K00760705 | **0.110** | **0.517** |
| BRD-K16189898 | **0.00279** | **0.00254** |
| BRD-K55000304 | **0.0198** | **0.00947** |
| BRD-K87550320 | **0.0110** | **0.00530** |
| BRD-K26994486 | >33 | >33 |
| BRD-K91354313 | >33 | >33 |
| BRD-K97242998 | >33 | >33 |
| BRD-K56373119 | >33 | >33 |
| BRD-K86938753 | >33 | >33 |
| BRD-K59522787 | >33 | >33 |
| BRD-K94445007 | >33 | >33 |
| BRD-K23628938 | >33 | >33 |
| BRD-K36176998 | >33 | >33 |
| BRD-K94445007 | >33 | >33 |
| BRD-K10781609 | >33 | >33 |
| BRD-K34254821 | >33 | >33 |
| BRD-K65054381 | >33 | >33 |
| BRD-K09376136 | >33 | >33 |
| BRD-K12710356 | >33 | >33 |
| BRD-K96280720 | >33 | >33 |
| BRD-K07274638 | >33 | >33 |
| BRD-K18142082 | >33 | >33 |
| BRD-K07891707 | >33 | >33 |
| BRD-K26973634 | >33 | >33 |
| BRD-K57195030 | >33 | >33 |
| BRD-K71568523 | >33 | >33 |
| BRD-K72681403 | >33 | >33 |
| BRD-K61284358 | >33 | >33 |
| BRD-K18599462 | >33 | >33 |
| BRD-K55276005 | >33 | >33 |

**H2T**

| Core ID | GSK3α $IC_{50}$ μM | GSK3β $IC_{50}$ μM |
|---|---|---|
| BRD-K00760705 | **0.121** | **0.583** |
| BRD-K16189898 | **0.00538** | **0.00371** |
| BRD-K55000304 | **0.0125** | **0.00693** |
| BRD-K87550320 | **0.0195** | **0.00802** |
| BRD-K91354313 | >33 | >33 |
| BRD-K26994486 | >33 | >33 |
| BRD-K12299093 | >33 | >33 |
| BRD-K42803695 | >33 | >33 |
| BRD-K53538949 | >33 | >33 |
| BRD-K61699306 | >33 | >33 |
| BRD-K74022101 | >33 | >33 |
| BRD-K83538063 | >33 | >33 |
| BRD-K97378175 | >33 | >33 |
| BRD-K00411149 | >33 | >33 |
| BRD-K04403306 | >33 | >33 |
| BRD-K11844860 | >33 | >33 |
| BRD-K20077349 | >33 | >33 |
| BRD-K34274486 | >33 | >33 |
| BRD-K54036439 | >33 | >33 |
| BRD-K59865704 | >33 | >33 |
| BRD-K74198088 | >33 | >33 |
| BRD-K06631334 | >33 | >33 |
| BRD-K28720546 | >33 | >33 |
| BRD-K31377965 | >33 | >33 |
| BRD-K33863933 | >33 | **23.8** |
| BRD-K44217085 | >33 | >33 |
| BRD-K48114548 | >33 | >33 |
| BRD-K58769468 | >33 | >33 |
| BRD-K76966915 | >33 | >33 |
| BRD-K80986480 | >33 | >33 |

***Chemical synthesis of BRD-K28862419***

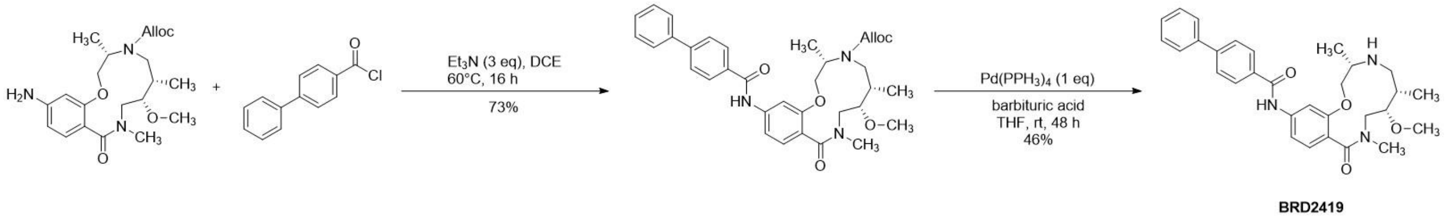


BRD2419

We followed prior work [17]. To a solution of the alloc protected aniline intermediate (45 mg, 0.111 mmol, 1.20 eq) and of 4-phenylbenzoyl chloride (20 mg, 0.0923 mmol, 1.00 eq) in CH2Cl2 (0.5 mL) was added triethylamine (0.277 mmol, 3 eq). The mixture was stirred at 60°C for 16 h. The solvent was evaporated, and the residue purified by normal phase chromatography (0-50% ethyl acetate in hexanes) to provide the diphenylamide (39 mg, 73% yield).

A vial of the alloc protected intermediate (51 mg) in tetrahydrofuran (0.5 mL) was sparged with Ag for 10 mins. In a separate vial Pd(PPH3)4 (102 mg, 0.0883 mmol, 1.00 eq) and barbituric acid (207 mg, 1.32 mmol, 15.0 eq) were combined and backfilled with Ag. Then, the THF solution was transferred (by syringe) to the powder. The reaction was stirred under nitrogen for 48 h. The reaction was concentrated then purified by normal phase chromatography (silica, 0 – 40% CH3OH in CH2Cl2) to provide N-((3S,6S,7S)-7-methoxy-3,6,9-trimethyl-10-oxo-3,4,5,6,7,8,9,10-octahydro-2H-benzo[k][1]oxa[4,9]diazacyclododecin-13-yl)-[1,1'-biphenyl]-4-carboxamide (21 mg, 0.0409 mmol, ?, 46% yield). MS: 502.96 [M+H] + 500.62 [M-H]-.

##### *Chemical re-synthesis of and BRD-K34692511*

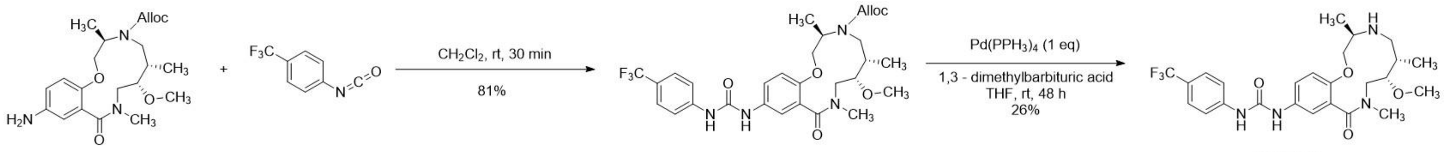


We followed prior work [17]. To a vial of the analine intermediate (42 mg, 0.104 mmol, 1.00 eq) in CH2Cl2 (0.5 mL) was added 1-isocyanato-4-(trifluoromethyl)benzene (0.01 mL, 0.104 mmol, 1.00 eq) and stirred at 25 °C for 30 min. The reaction was concentrated and the residue was purified by normal phase chromatography (0-100% ethyl acetate in hexanes) to provide the protected urea intermediate (52 mg, 81% yield).

A vial of the urea intermediate (52 mg, 0.0871 mmol, 1.00 eq) in tetrahydrofuran (0.4 mL) was sparged with Ag for 10 mins. In a separate vial Pd(PPH3)$_4$ (101 mg, 0.0871 mmol, 1.00 eq) and 1,3 - dimethylbarbituric acid (14 mg, 0.0871 mmol, 1.00 eq) were combined and backfilled with Ag. Then the THF solution was transferred (by syringe) to the powder. The reaction was stirred under nitrogen for 48 h. The reaction remained bright yellow. The reaction was concentrated then purifed by normal phase chromatography (0-100% CH3OH in CH2Cl2, product elutes). The collected fractions were concentrated and purified with a focused gradient of 35-45% CH3OH in CH2Cl2 was run to separate remaining impurities. Fractions containing product were collected to provide 1-[(4R,7S,8S)-8-methoxy-4,7,10-trimethyl-11-oxo-2-oxa-5,10-diazabicyclo[10.4.0]hexadeca-1(12),13,15-trien-14-yl]-3-[4-(trifluoromethyl)phenyl]urea, (13 mg, 0.0224 mmol, 26% yield). MS: 510.63 [M+H]+, 553.860 [M+CHOO-]-.

#### Supplementary Figures:

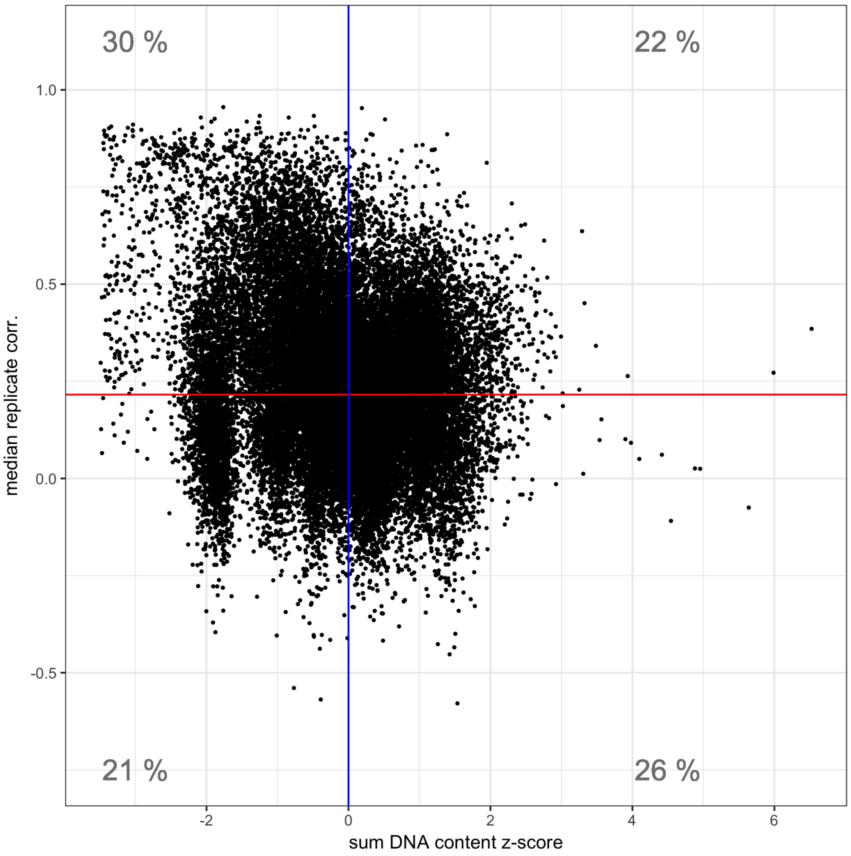


***Supplementary Figure S1: Relationship between detectable Cell Painting profiles and cell proliferation rules out toxicity being a single, dominant phenotype.*** *The Y axis shows the replicate correlation, which is high for compounds that produce detectable morphological phenotypes in the Cell Painting assay. 52% of the compounds have a replicate correlation higher than the 95th percentile of non-replicate correlations (red line) and thus are considered to have a detectable phenotype. The X axis shows the z-score for the sum of DNA content, where higher values represent higher cell proliferation. Although the ratio of low-proliferation samples (left of blue line) with a detectable phenotype (30% vs. 21%) is higher than for high-proliferation samples (right of blue line) (22% vs. 26%), it is clear that impact on cell proliferation does not explain the majority of detectable morphological phenotypes. This fact is also supported by observing no significant negative correlation between proliferation and replicate correlation.*

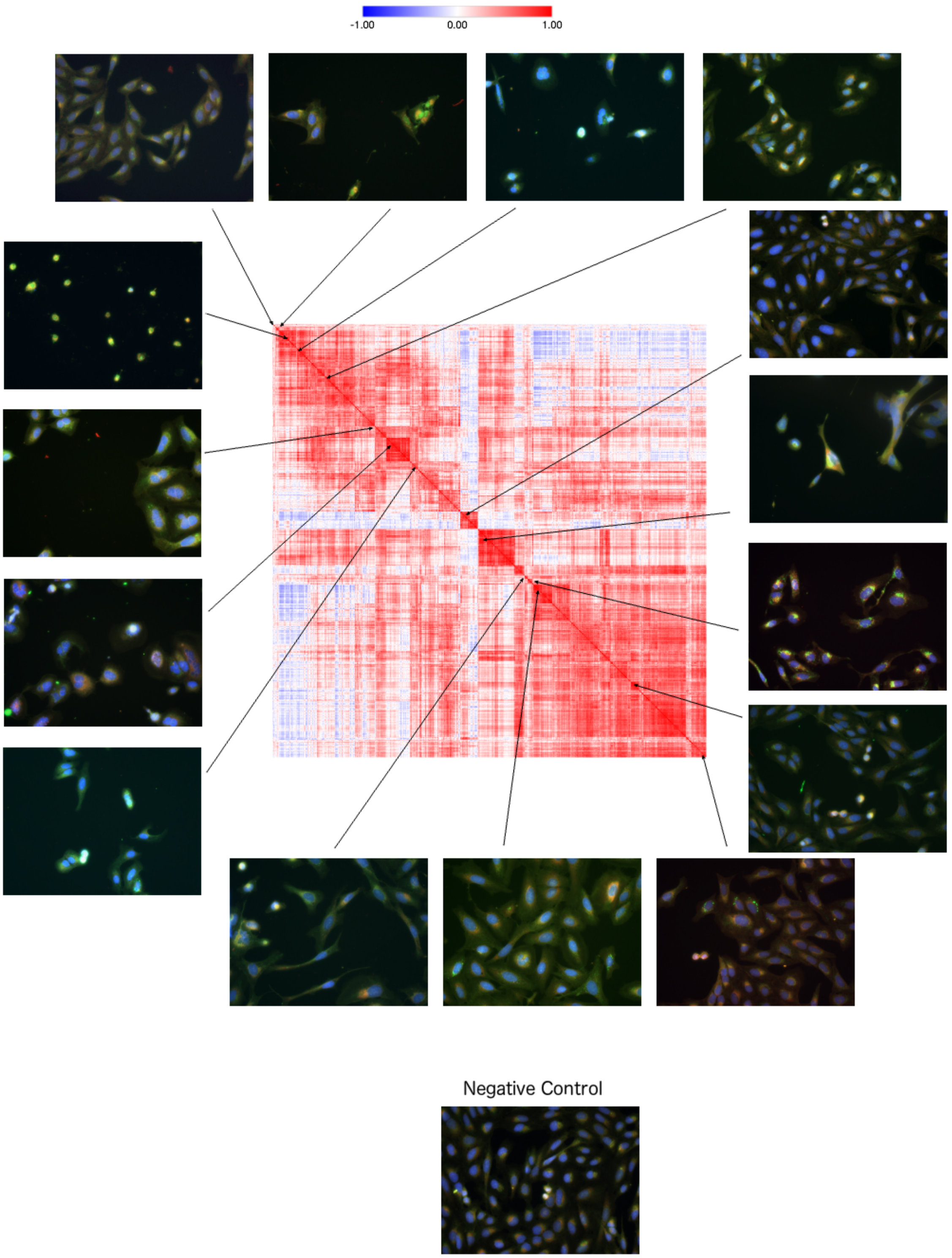


***Supplementary Figure S2: Compounds yielding a low cell count may be toxic or proliferation-impeding but they display many distinguishable phenotypes.*** *Low-cell-proliferation or potentially toxic compounds (with the z-score for the sum of DNA content less than -3) are clustered, and show many different types of toxic phenotype. Various tight clusters mean the assay is specific and has sufficient resolution to distinguish types of toxicity.*

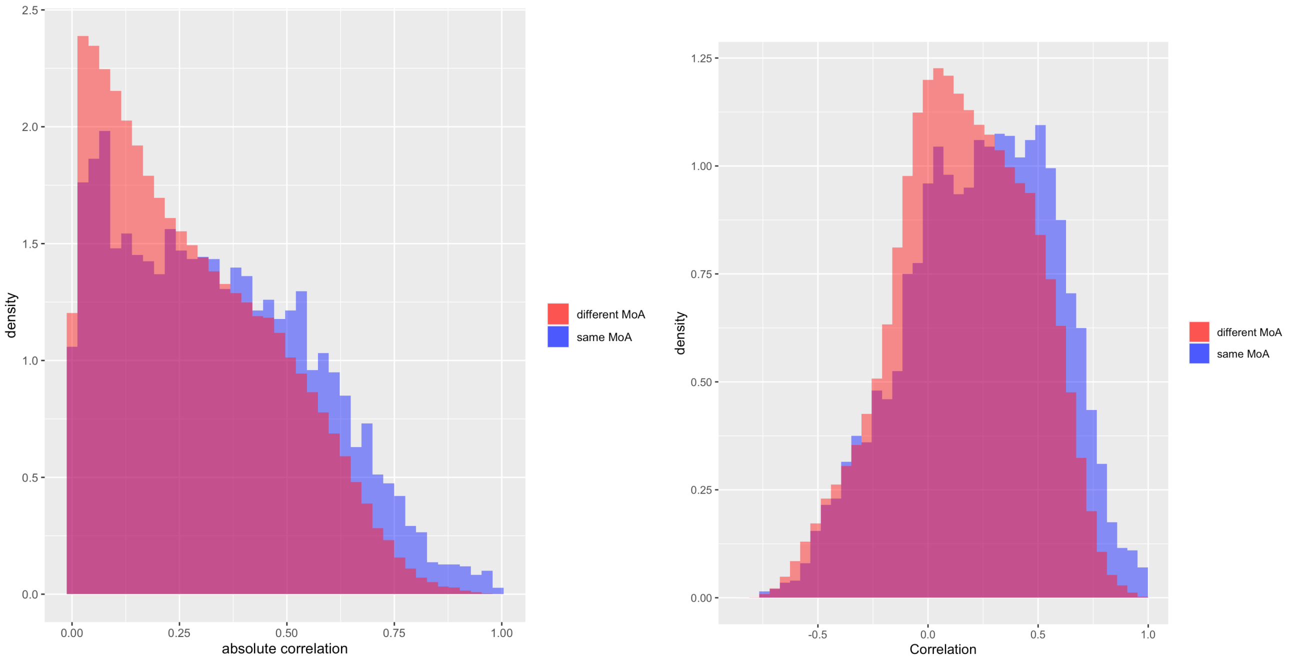


***Supplementary Figure S3: Correlation values of same-MOA compound pairs are shifted slightly compared to the different-MoA compound pairs.*** *Shown are the absolute correlations (left) and the correlations (right), with the latter keeping the directionality of correlation intact (showing almost entirely positive correlations, as expected for compounds sharing the same MOA).*

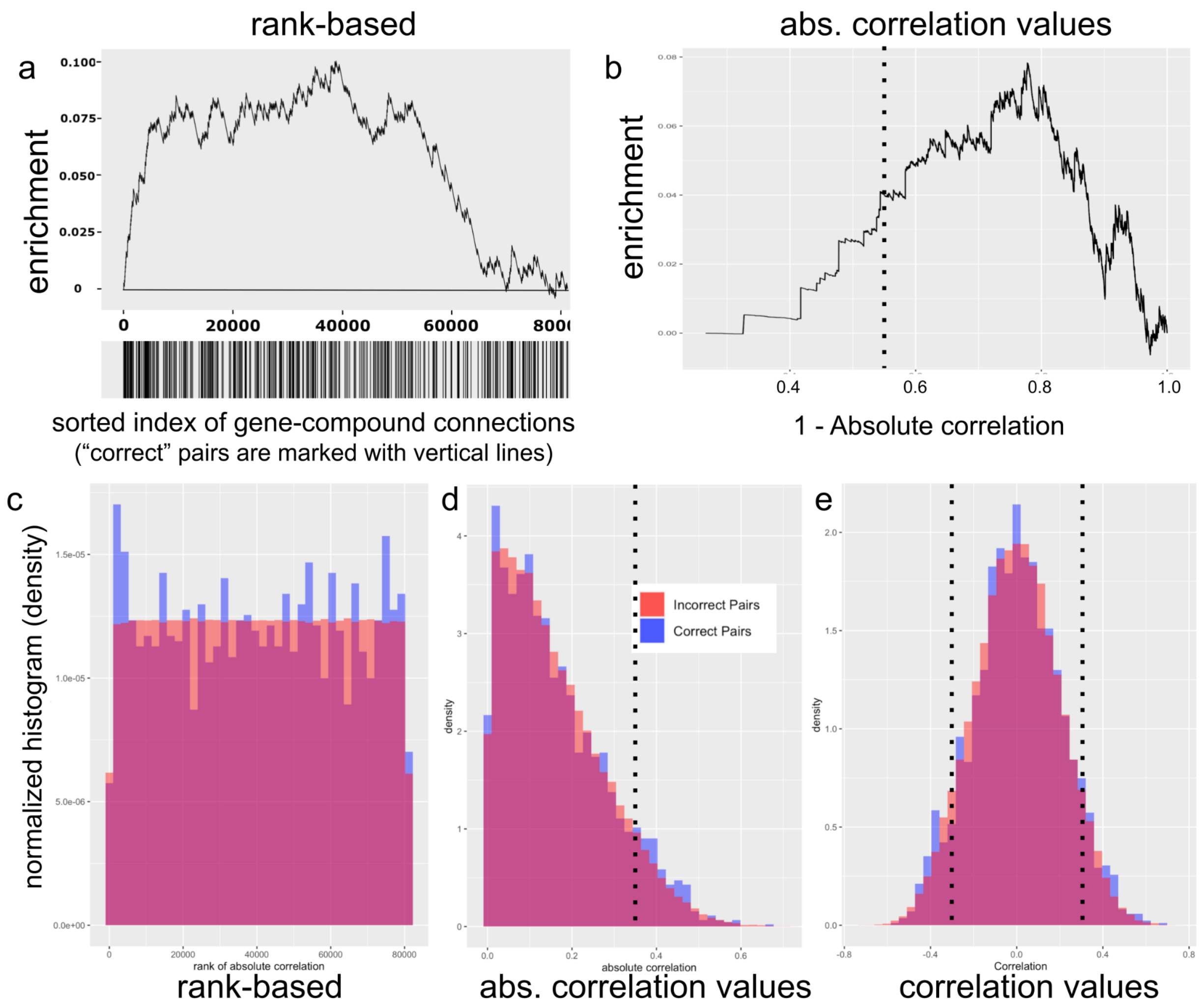


***Supplementary Figure S4: Enrichment analysis and histograms of gene-compound matches in the validation set.***

*The rank order or correlation values of "correct" gene-compound pairs (based on existing annotations) is shown relative to "incorrect" ones (note: there may be many not-previously-known gene-compound matches in the set that are actually correct but just unannotated). In contrast to the gene-centric analysis that yielded 20 successful genes out of 63 in Figure 1c, here we examine* ***all*** *gene-compound pairs together and limit matching to the 1177 compounds in the set that are annotated as targeting at least one of the 69 genes. We also separately plot rank orders and correlations. Although most correct gene-compound pairs did not correlate highly, as expected for the reasons described in the main text, nevertheless we care most about the strongest connections (correlation ≥|0.35|), where correct gene-compound pairings were 2.5-fold overrepresented among the strongest gene-compound pairings (p-value = 0.007; details of the analysis are in Supplementary Table S1). Note that in a screening context, false negatives are routine and false positives are readily filtered at a later stage so long as their rate is not overwhelming; the goal is to obtain at least some true positives among the top-ranked compounds. Defining correctness differently – based on whether the Gene Ontology biological processes intersection of two proteins is greater than 50% of their union – is less successful (odds ratio of 1.24 with p-value = 0.0148 in the one-sided Fisher's test). When correlation values are shown, the threshold of |0.35| used in the study is marked with a black dotted line.*

*a) Enrichment plot of all gene-compound pairs rank-ordered based on their absolute profile correlation. Starting from the left, the curve rises a unit if the gene-compound pair is a known/annotated connection, and goes down a unit otherwise. The units are normalized to the number of possible known connections, so the maximum height is one and ends in zero. The steep increase of the curve indicates enrichment of correct connections towards the top of the rank-ordered list of pairs. Correct pairs are also noted as black vertical lines on the x axis. More details in "Enrichment analysis plots" in Methods. b) Same as (a) except using sorted gene-compound absolute correlation values rather than their rank. c) Using a histogram rather than an enrichment plot, the best-ranking gene-compound pairs (e.g., the first three bars with highest absolute correlation) contain more correct pairs than incorrect ones. d) Same as (c) except using sorted gene-compound absolute correlation values rather than their rank. e) Same as (d) except using actual correlation values rather than absolute values (keeping the directionality of correlation intact and showing a mixture of negative and positive correlations above the threshold of |0.35| used in the study).*

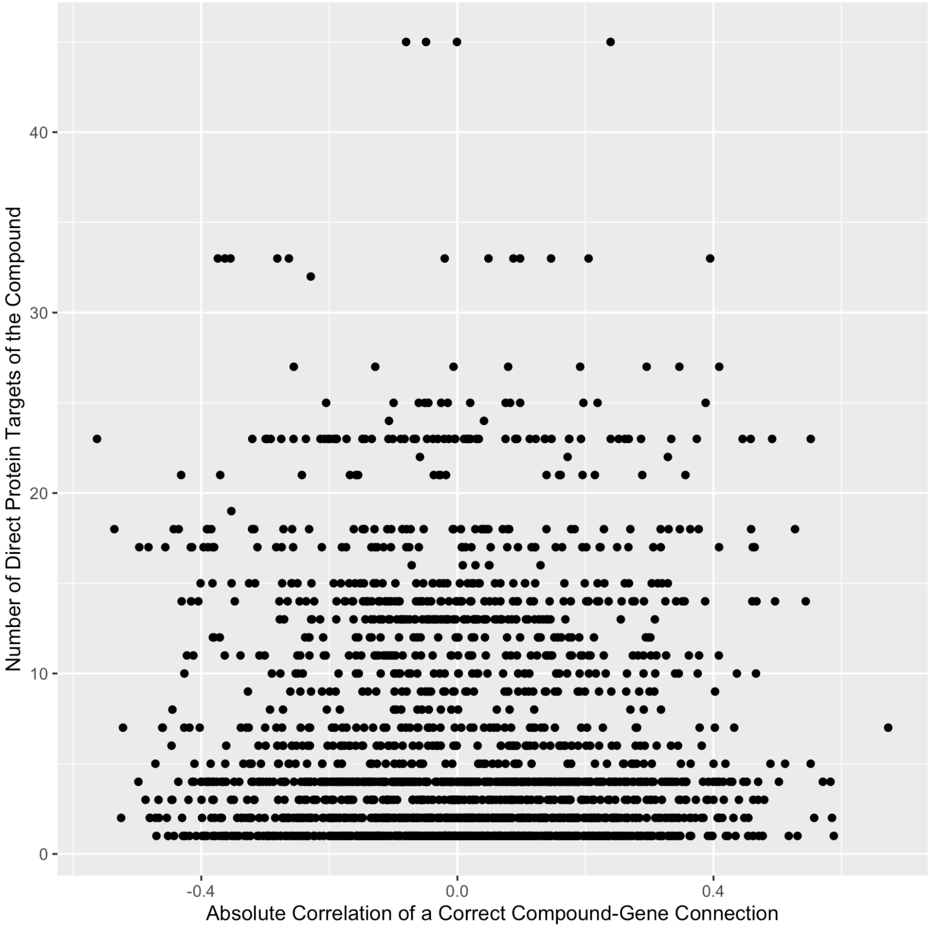


***Supplementary Figure S5: Gene-compound matching is uncorrelated to the number of compound targets for correct matches.*** *In our validation analysis testing known gene-compound pairs, a gene is considered as a correct match for a given compound if its protein is one of the compound's known (annotated) protein targets, or is reported to interact with one of those targets, per BioGrid. The x-axis shows the absolute correlation of compounds to their correct matches. Each point represents a true compound-gene connection.*

*The y-axis shows the number of direct protein targets of a compound. Spearman correlation of x-axis and y-axis is ~ -0.02.*

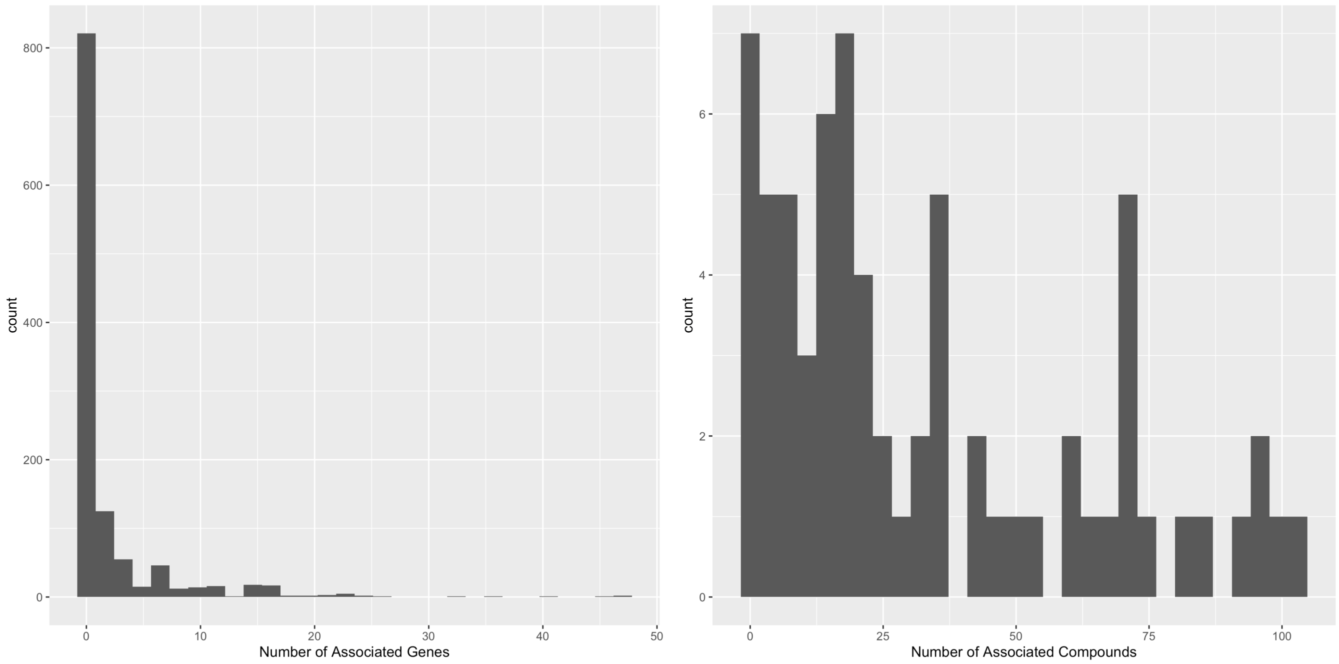


***Supplementary Figure S6: Polypharmacology analysis of the compounds.*** *a) The majority of compounds in the experiment are associated with fewer than four genes in the genome (including links via BioGrid interactions). Note that only ~1000 compounds total are annotated with a gene in our set; many are uncharacterized. b) The majority of genes in the set of 63 validation genes are annotated as associated with fewer than 32 compounds (including links via BioGrid interactions).*

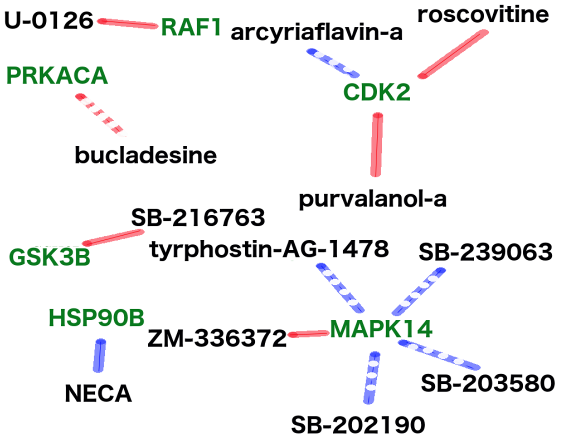


***Supplementary Figure S7: Known gene-compound connections with morphological matches.*** *Of the 63 genes that have a bioactive compound annotated as targeting it in the dataset, we analyzed the six genes (green text) that most strongly matched or opposed the correct compound(s) (black text), where correct here includes only the direct annotated targets of the compounds (in contrast to the analysis in the validation section of the paper which included interacting proteins of the targets as 'correct' matches). The lines represent positive (blue) and negative (red) morphological correlations to compounds. They also show whether the morphological correlation is the expected (solid) or unexpected directionality (dotted) based on previously described positive or negative impacts on gene function.*

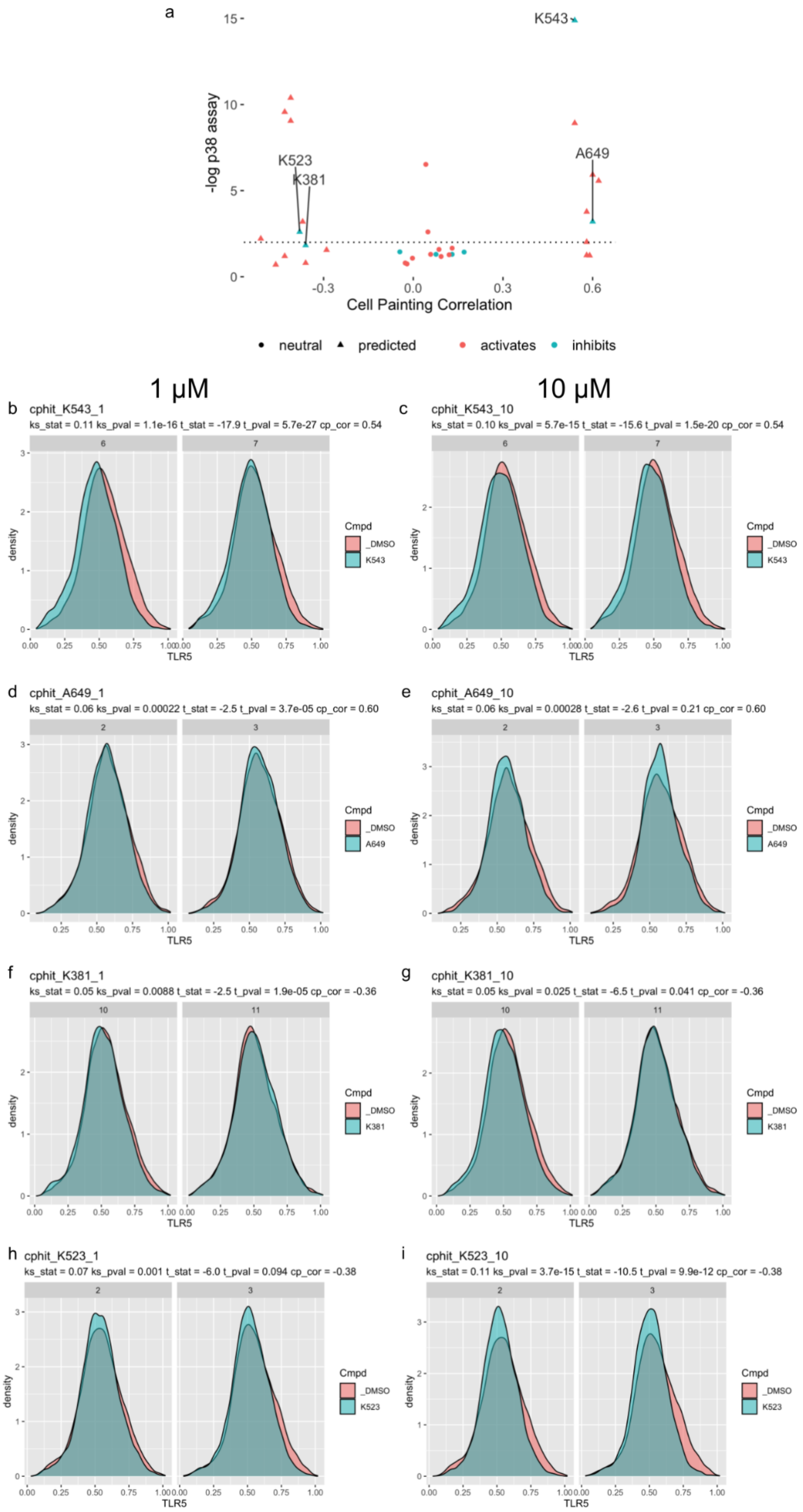
a
K543
K523
K381
A649
-log p38 assay
Cell Painting Correlation
neutral
predicted
activates
inhibits
1 µM
10 µM
b
cphit_K543_1
ks_stat = 0.11 ks_pval = 1.1e-16 t_stat = -17.9 t_pval = 5.7e-27 cp_cor = 0.54
c
cphit_K543_10
ks_stat = 0.10 ks_pval = 5.7e-15 t_stat = -15.6 t_pval = 1.5e-20 cp_cor = 0.54
d
cphit_A649_1
ks_stat = 0.06 ks_pval = 0.00022 t_stat = -2.5 t_pval = 3.7e-05 cp_cor = 0.60
e
cphit_A649_10
ks_stat = 0.06 ks_pval = 0.00028 t_stat = -2.6 t_pval = 0.21 cp_cor = 0.60
f
cphit_K381_1
ks_stat = 0.05 ks_pval = 0.0088 t_stat = -2.5 t_pval = 1.9e-05 cp_cor = -0.36
g
cphit_K381_10
ks_stat = 0.05 ks_pval = 0.025 t_stat = -6.5 t_pval = 0.041 cp_cor = -0.36
h
cphit_K523_1
ks_stat = 0.07 ks_pval = 0.001 t_stat = -6.0 t_pval = 0.094 cp_cor = -0.38
i
cphit_K523_10
ks_stat = 0.11 ks_pval = 3.7e-15 t_stat = -10.5 t_pval = 9.9e-12 cp_cor = -0.38
density
TLR5
Cmpd
_DMSO
K543
A649
K381
K523

***Supplementary Figure S8: Predicted compounds impact p38 activity in a single-cell reporter assay.*** *a) The same experiment as shown in Figure 2 is shown here, except using a Kolmogorov-Smirnoff (KS) analysis to detect differences in distribution instead of shifts in the mean. This raises an additional hit, K523. b-i) Single cell distribution plots show the shifts induced, at both 1μM and 10μM, by a known inhibitor of p38, SB202190 (b-c), by the two hits from the t-test in Figure 2 (d-g) and by the hit from the KS test (h-i). Note that the biological effect size here is relatively small, even for the known p38 inhibitor; this is typical for the assay.*

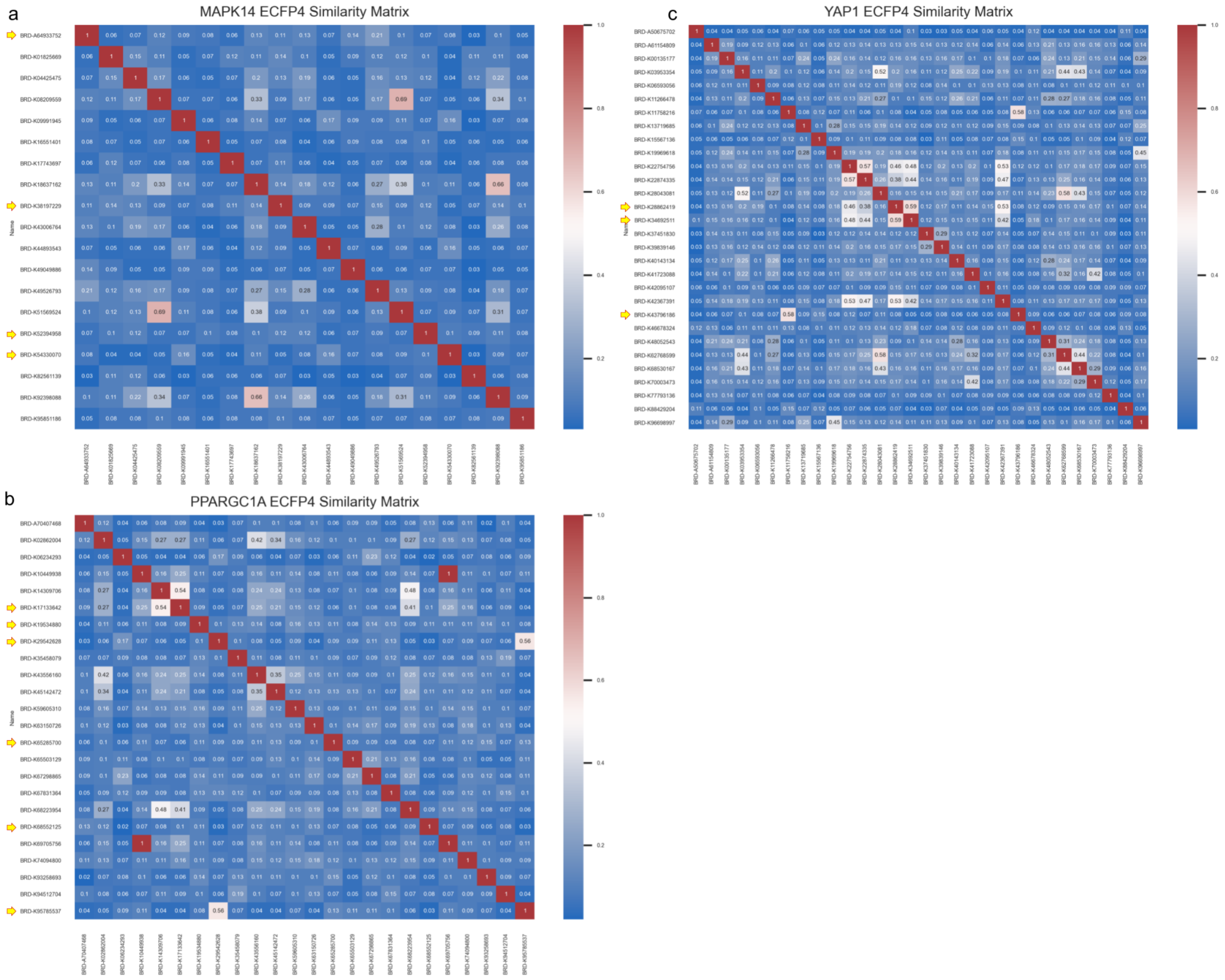


***Supplementary Figure S9: Similarity among chemical structures of compounds matching/opposing each gene.*** *Although hits by morphological matching share a Cell Painting activity profile in common, their structures can be quite diverse. (a) MAPK14: the four compounds marked with a yellow arrow are those highlighted in Figure 2. (b) PPARGC1A: the six compounds marked with a yellow arrow are those mentioned in the legend of Supplementary Figure S12. (c) YAP1: the three compounds marked with a yellow arrow are those heavily studied in the text.*

a

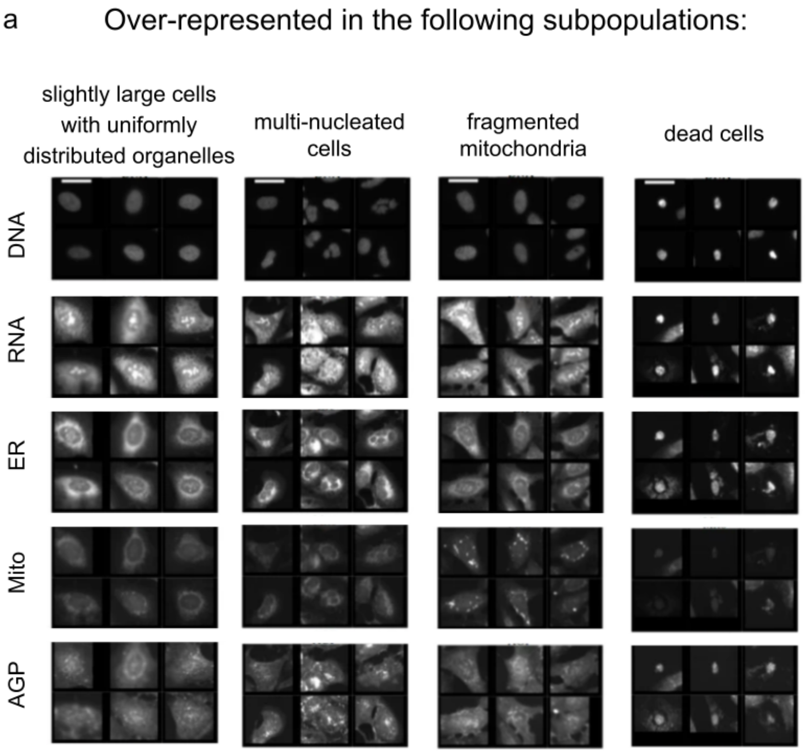


b

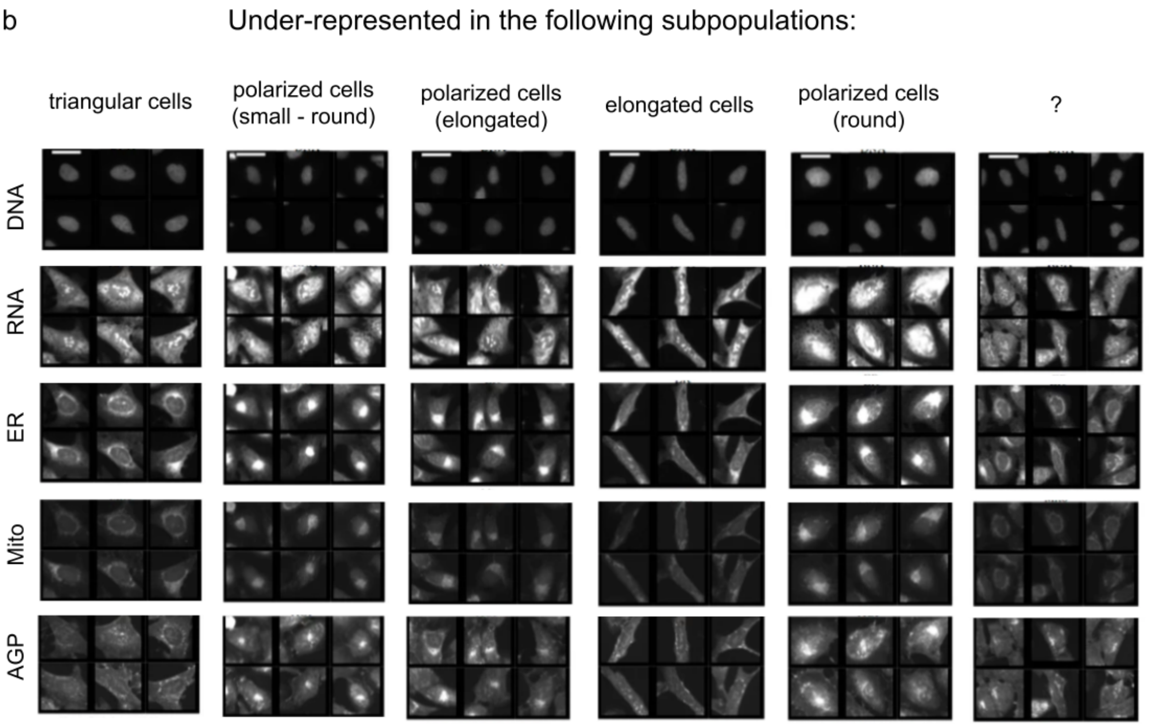


***Supplementary Figure S10: Certain subpopulations of cells are over- or under-represented when PPARGC1A is overexpressed.*** *Following the procedure described previously[1] we clustered cells based on their morphological profiles, then identified which subpopulations were (a) over- or (b) under-represented when PPARGC1A is overexpressed. Scale bars = 39.36 $\mu$m.*

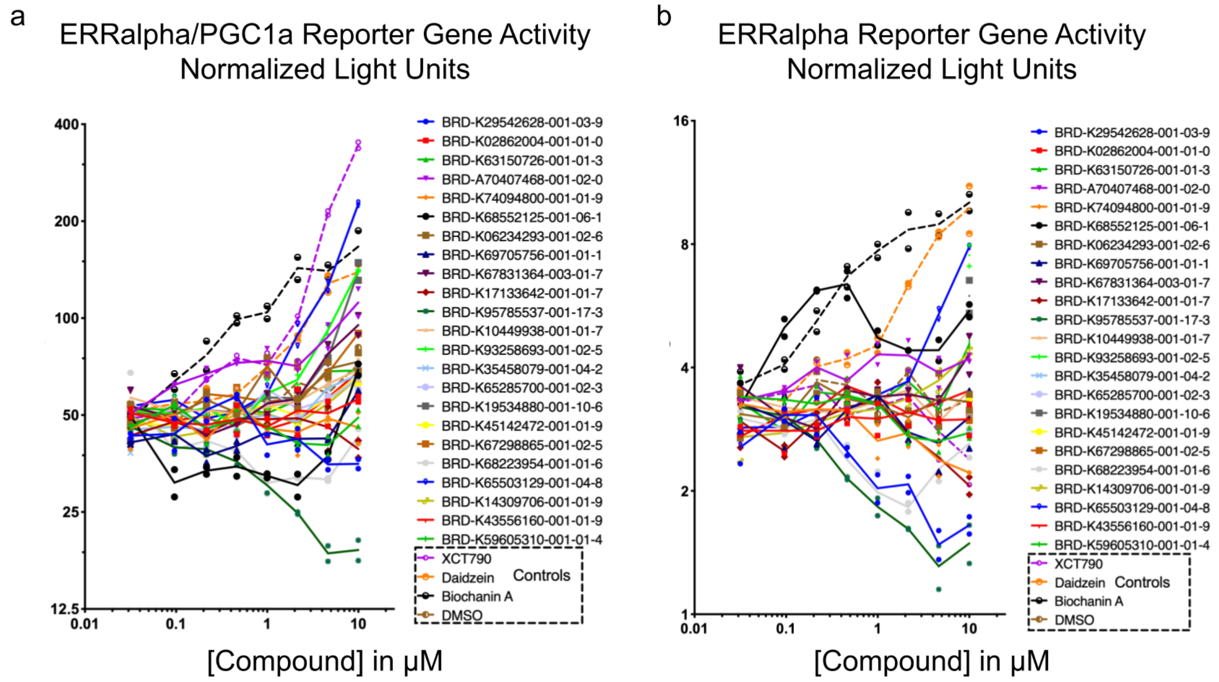


***Supplementary Figure S11: Compounds predicted to influence pathways containing PGC1α impact an ERRa reporter assay in 293T cells.*** *In this reporter system, a mammalian one-hybrid fusion protein containing the Gal4 DNA binding domain and the ERR alpha ligand binding domain is co-expressed with the Firefly luciferase gene under control of the Gal4 Upstream Activating Sequence. Renilla luciferase was included for normalization. The assay was performed in the presence (a) or absence (b) of ectopically expressed PGC1α; their behavior being similar in these two conditions suggests, but does not prove, that the compounds do not directly target PGC1α but instead modulate other targets in the relevant pathway, consistent with having been discovered by the morphological matching approach which assesses impact on the cell system rather than a particular desired target.*

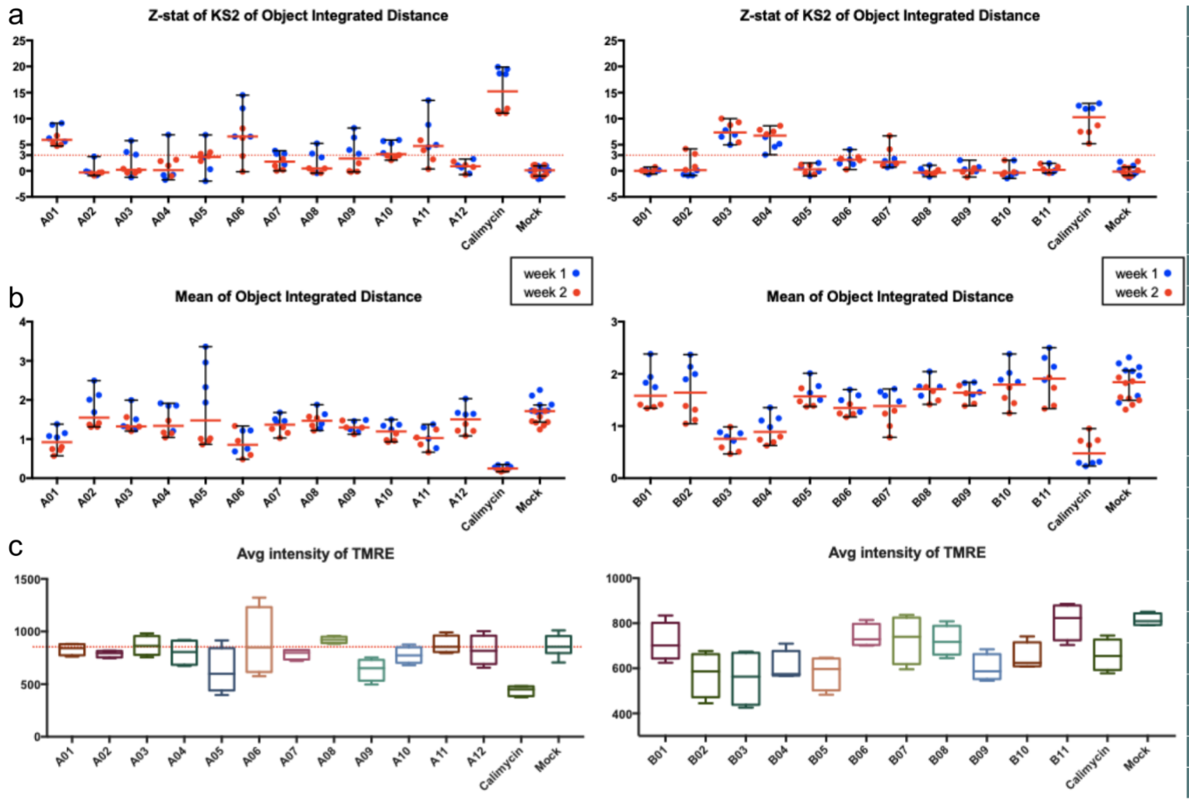


| Well | Broad ID |
|---|---|
| A01 | BRD-K29542628-001-03-9 |
| A02 | BRD-K02862004-001-01-0 |
| A03 | BRD-K63150726-001-01-3 |
| A04 | BRD-A70407468-001-02-0 |
| A05 | BRD-K74094800-001-01-9 |
| A06 | BRD-K68552125-001-06-1 |
| A07 | BRD-K06234293-001-02-6 |
| A08 | BRD-K69705756-001-01-1 |
| A09 | BRD-K67831364-003-01-7 |
| A10 | BRD-K17133642-001-01-7 |
| A11 | BRD-K95785537-001-17-3 |
| A12 | BRD-K10449938-001-01-7 |
| **Well** | **Broad ID** |
| B01 | BRD-K93258693-001-02-5 |
| B02 | BRD-K35458079-001-04-2 |
| B03 | BRD-K65285700-001-02-3 |
| B04 | BRD-K19534880-001-10-6 |
| B05 | BRD-K45142472-001-01-9 |
| B06 | BRD-K67298865-001-02-5 |
| B07 | BRD-K68223954-001-01-6 |
| B08 | BRD-K14309706-001-01-9 |
| B09 | BRD-K65503129-001-04-8 |
| B10 | BRD-K43556160-001-01-9 |
| B11 | BRD-K59605310-001-01-4 |

***Supplementary Figure S12: Predicted compounds impact a mitochondrial motility assay in rat cortical neurons.*** *(a) For most compounds, the integrated distance traveled for each motile mitochondrion (the length of travel, or the sum of all movements, including changes in direction) is comparable to the negative control (Mock), but a few (A01, A06, A10, A11, B03, and B04) consistently have a z-score >3, as does the positive control, Calcimycin, a calcium ionophore that arrests mitochondria [18]. Two separate experiments are plotted (week 1 in blue and week 2 in red), and the values are the Z-prime factor of the Kolmogorov-Smirnov (KS) statistic calculated for each compound. The red line indicates the median +- 95% confidence interval. (b) Mean values of the mitochondrial distance; these are the values that underlie the statistical analysis in (a). The red line indicates the median +- 95% confidence interval. (c) The average intensity of TMRE reflects the mitochondrial membrane potential, a measure of mitochondrial function. Boxplots show the median and 25th/75th percentiles, with whiskers showing the most extreme observation less than or equal to the upper hinge + 1.5 * inter-quartile range. Interestingly, A01, A06 and A11 all show normal levels of TMRE staining, suggesting a specific effect on mitochondrial motility rather than a more general decrease in neuronal or mitochondrial health. This cannot be said for B03 and B04 (and A10 to a lesser extent), which apparently reduce membrane potential, although additional validation with TMRE is needed to conclude that they are in fact detrimental to cell health. Of note, four of these compounds were also active in the PPARG reporter assay (Figure 3c): A01 and A11 are structurally related molecules of the pyrazolo-pyrimidine family, 1-Naphthyl-PP1 and PP2, which are Src family kinase inhibitors with additional targets including TGFbeta receptors and others. A06 is Phorbol myristate acetate (aka TPA, PMA). B09 is annotated as an HSP-90 inhibitor CCT-018159. 23 compounds were tested because one of the original 24 tested in Figure 3c became unavailable.*

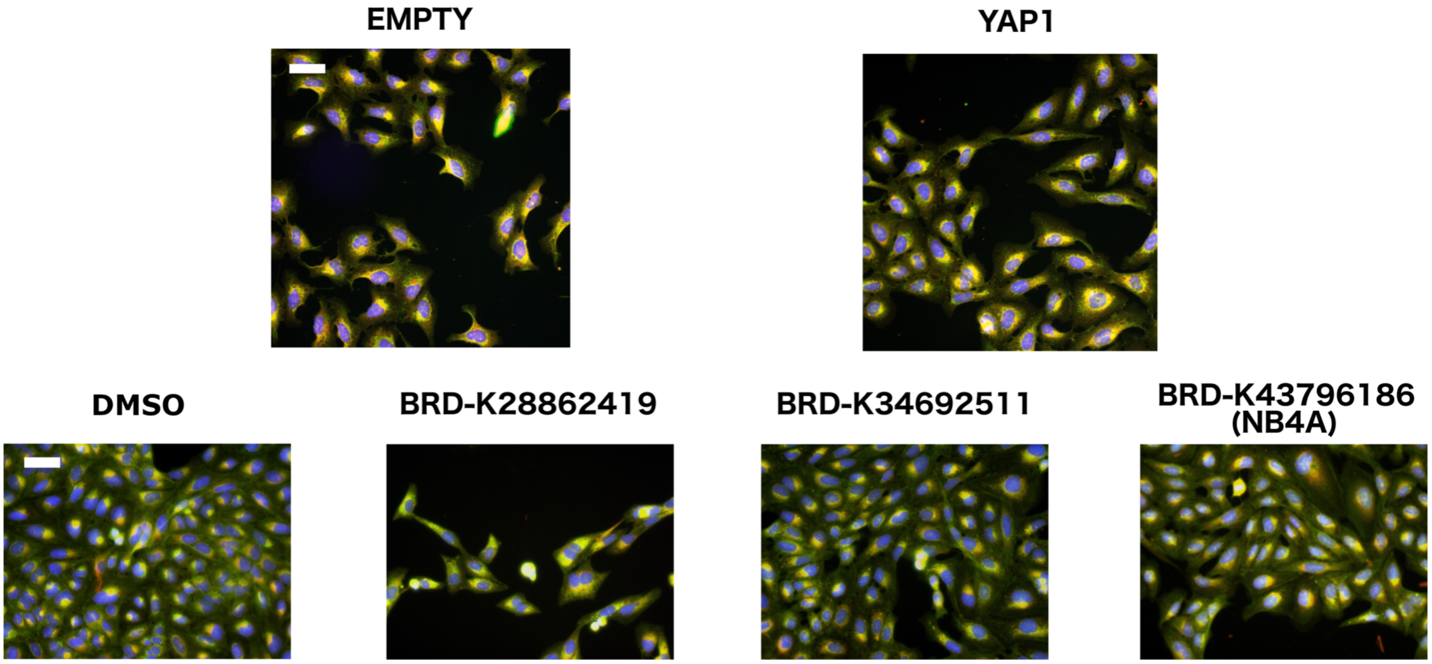


***Supplementary Figure S13: Cell Painting images related to the YAP1 pathway in U2OS cells.*** *Top: Cell Painting images for YAP1 overexpression compared to negative control (EMPTY, same image as in Figure 1b). Overexpressing YAP1 produces elongated cells with more cell protrusions, lower RNA staining, and disjoint, bright mitochondria patterns. Bottom: Cell Painting images for the negative control (DMSO, same image as in Figure 1b) and three compounds that correlated negatively or positively to the YAP1 overexpression profile. NB4A (BRD-K43796186) was positively correlated and the other two negatively correlated. Scale bars = 60* $\mu$*m.*

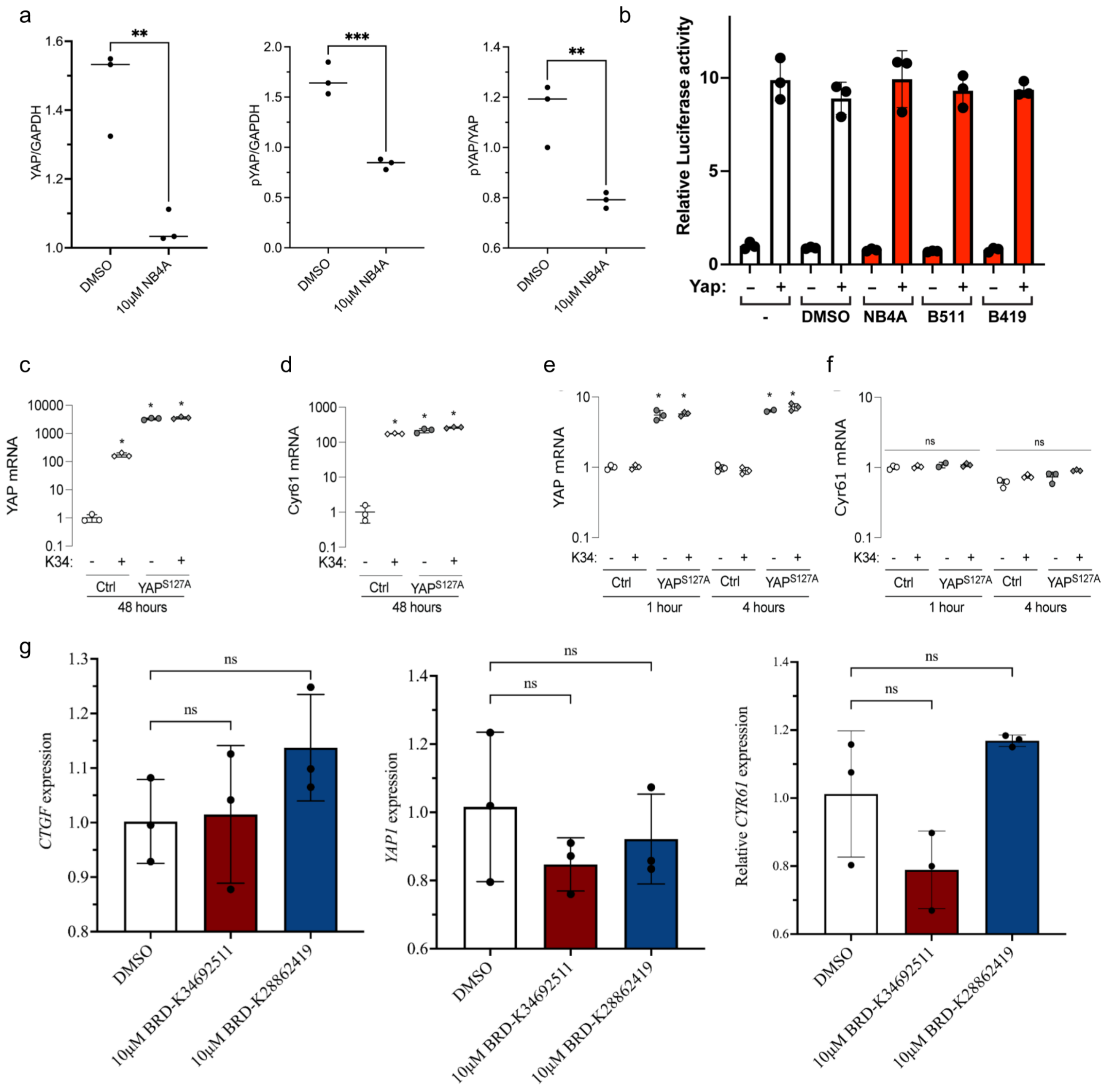


**_Supplementary Figure S14: Analysis of selected compounds in various YAP-related contexts._**

*a) Quantification of relative levels of total YAP1 and phospho-YAP1 in H9 hPSCs after treatment with DMSO or NB4A for 24 hours. **P<0.01; ***P<0.001 (Two-tailed student's t-test). Mean ± SD. n = 3 biologically independent experiments. A representative example western blot is shown in Figure 4c. b) A TEAD luciferase reporter was co-transfected with or without a Yap expression construct into HEK293T cells followed by treatment for 48 hours with DMSO or the indicated compounds, which appear to have no effect. The data shown are the average of three samples within a representative experiment ± SEM. c-f) BRD-K34692511 upregulates YAP1 and target-gene mRNA levels in murine periosteal cells: c, d) YAP1 and Cyr61 mRNA levels* *in murine periosteal cells after 48 hours of treatment with BRD-K34692511 (K34) in the presence or absence of doxycycline-induced YAP$^{S127A}$. e, f) YAP1 and Cyr61 mRNA levels after 1 and 4 hours of treatment. Gene expression was evaluated by one and two-way ANOVA with Tukey post hoc test n=3/group/time-point. * indicates $p<0.05$ compared to untreated controls. g) BRD-K28862419 and BRD-K34692511 did not dramatically impact mRNA levels of Hippo pathway members in hPSCs. Relative transcript levels of YAP1, CTGF, and CYR61 from H9 hPSCs treated with DMSO, BRD-K28862419, or BRD-K34692511 for 24 hrs. Error bars represent mean + SEM, from n=3 independent biological replicates (one-way ANOVA with Dunnett multiple comparison test).*

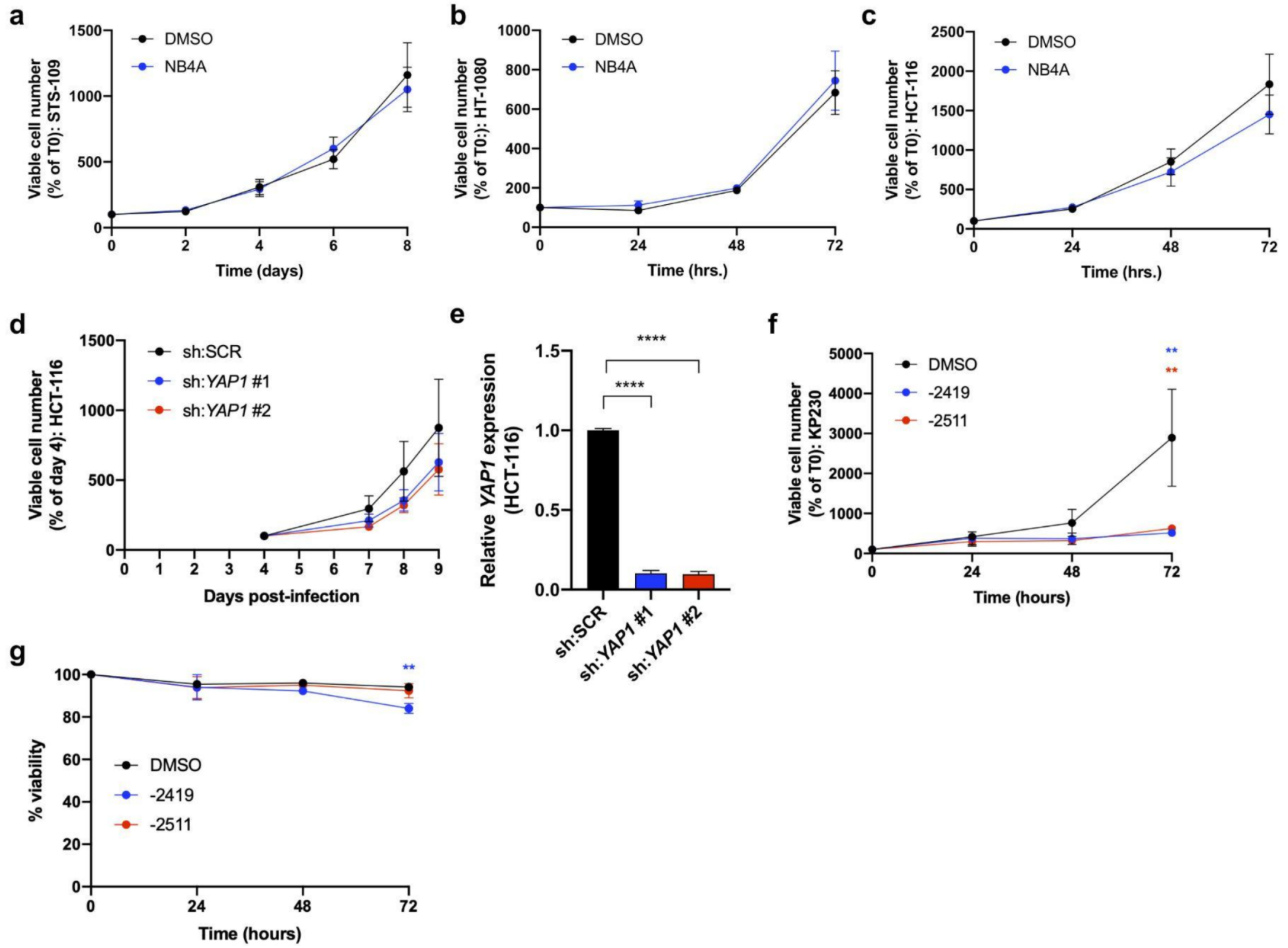


***Supplementary Figure S15: Predicted YAP1-related compounds impact proliferation in a cell type-specific manner.*** *a, b) Growth curves of YAP1-dependent human sarcoma cells[14,19] treated with 10 µM NB4A or DMSO control. c) Growth curve of HCT-116 colon cancer cells treated with 10 µM NB4A or DMSO control. a-c are not significantly different at any time point (2-way ANOVA with Sidak's multiple comparisons test). n = 3. Mean ± SEM. d) Growth curve of HCT-116 cells infected with YAP1-targeting shRNAs or scrambled shRNA control (sh:SCR); no conditions were significantly different at any time point (vs. sh:SCR; 2-way ANOVA with Dunnett's multiple comparisons test). n = 3. Mean ± SEM. e) Relative YAP1 expression in the cells depicted in panel d ****P<0.0001 vs. sh:SCR (1-way ANOVA with Dunnett's multiple comparisons test). n = 3. Mean ± SEM. f) Growth curves of KP230 cells treated with 10 µM BRD-K28862419, BRD-K34692511, or DMSO control. **P<0.01 vs. DMSO (72 hrs.; 2-way ANOVA with Dunnett's multiple comparisons test). n = 2 Mean ± SEM. g) Percent viability of KP230 cells depicted in panel f **P<0.01 vs. DMSO (72 hrs.; 2-way ANOVA with Dunnett's multiple comparisons test). n = 3. Mean ± SEM. For panels a, b, c, f, and g, cells were treated with 10 µM of the indicated inhibitor daily for 72 hours.*
